# Dynamic evolution of retroviral envelope genes in egg-laying mammalian genomes

**DOI:** 10.1101/2022.12.20.521333

**Authors:** Koichi Kitao, Hiyori Shoji, Takayuki Miyazawa, So Nakagawa

## Abstract

Independently acquired envelope (*env*) genes from endogenous retroviruses have contributed to the placental trophoblast cell-cell fusion in therian mammals. Egg-laying mammals (monotremes) are an important sister clade for understanding mammalian placental evolution, but the *env* genes in their genomes have yet to be investigated. Here, *env*-derived open reading frames (*env*-ORFs) encoding more than 400 amino-acid lengths were searched in the genomes of two monotremes: platypus and echidna. Only two *env*-ORFs were present in the platypus genome, whereas 121 *env*-ORFs were found in the echidna genome. The echidna *env*-ORFs were phylogenetically classified into seven groups named env-Tac1 to -Tac7. Among them, the env-Tac1 group contained only a single gene, and its amino acid sequence showed high similarity to those of the RD114/simian type D retroviruses. Using the pseudotyped virus assay, we demonstrated that the Env-Tac1 protein utilizes echidna sodium-dependent neutral amino acid transporter type 1 and 2 (ASCT1 and ASCT2) as entry receptors. Moreover, the Env-Tac1 protein caused cell-cell fusion in human 293T cells depending on the expression of ASCT1 and ASCT2. These results illustrate that fusogenic *env* genes are not restricted to placental mammals, providing insights into the evolution of retroviral genes and the placenta.

## Introduction

Endogenous retroviruses (ERVs) are remnants of ancient retroviral infections to host germ cells. ERVs have provided de novo genes and new functions in their hosts (1). In therian mammals, various retroviral envelope (*env*) genes have been reported to be co-opted for placental fusogenic genes during evolution, the original of which is responsible for viral entry by fusing the retrovirus and the target cell membrane. *Syncytin-1* and *syncytin-2* derived from ERV-W and ERV-FRD1, respectively, are thought to be responsible for the cell-cell fusion of human placental trophoblasts (2-5). Placental genes derived from *env* are not restricted to humans, but also to rodents (6-9), rabbits (10), carnivorans (11), ruminants (12, 13), tenrecs (14), and even in opossums belonging to marsupials (15). Moreover, non-mammalian species such as the viviparous *Mabuya* lizards were reported to acquire an *env*-derived gene, *syncytin-Mab1*, which was involved in placental cell-cell fusion (16). However, given the widespread occurrence of ERVs in vertebrates (17) and the ancient origin of retroviruses (18), the ERV invasion into the host genomes should have been occurring in egg-laying mammals (i.e., monotremes), which would be of particular interest to speculate the situation before the placental birth in mammals. Recently, the chromosome-level-assembled genomes of platypus and short-beaked echidna have been reported (19). It enables us to investigate the monotreme ERVs in more detail. Indeed, in platypus and echidna genomes, we previously reported highly conserved genes that were derived from a retroviral reverse transcriptase, although their molecular functions are unclear (20). Here, we comprehensively identified nearly full-length *env*-coding open reading frames (hereafter called *env*-ORFs) in the platypus and echidna genomes and found their dynamic evolution. Furthermore, a fusogenic Env protein identified in the echidna genome was experimentally verified. Our study demonstrates the ongoing acquisition of fusogenic *env* genes in mammals that do not utilize a placenta, providing key insights into the evolution of retroviral genes and the placenta.

## Results

### Discovery of *env*-ORFs in the platypus and echidna genomes

To obtain ORFs encoding nearly full-length envelope proteins in monotremes, we first extracted more than ORFs encoding more than 400-amino-acid length (i.e., 1200 nucleotides starting from the start codon) from the chromosome-level-assembled reference genomes of platypus and echidna (19). We then extracted ORFs encoding envelope proteins by sequence similarity search (see Materials and Methods). This search identified only two *env*-ORFs in the platypus genome. This observation is consistent with previous studies showing very few ERV-derived ORFs in the platypus (21, 22). In contrast, we identified 121 *env*-ORFs in the echidna genome. These monotreme *env*-ORFs were clustered into nine groups based on the putative amino acid sequences. We named these *env*-ORF groups as env-Oan1 and -Oan2 (Platypus: *Ornithorhynchus anatinus*) and env-Tac1 to -Tac7 (Echidna: *Tachyglossus aculeatus*) (Fig. 1; Dataset S1). The env-Oan1 and -Oan2 are found to have only one copy for each in the platypus genome. For echidna, the env-Tac1, -Tac2, -Tac4, and -Tac7 were also low copies (one copy for env-Tac1 and -Tac2, two copies for env-Tac7, and three copies for env-Tac4), while env-Tac3, - Tac5, and -Tac7 had 18, 77, and 19 copies, respectively. These high copy number *env*-ORFs were shallowly branched in the phylogenetic tree (Fig. 1), suggesting the recent expansion in the echidna genome. Proviral structures containing *env*-ORFs were identified based on their genomic sequences (Fig. 2; Dataset S2). Note that, for the high copy number env-ORFs (i.e., env-Tac3, - Tac5, and -Tac6), we generated consensus sequences of *env*-ORFs with flanking sequences and identified their proviral structures. The topologies of the phylogenetic trees of putative Gag and Pol proteins were consistent with those of the Env proteins and also suggested that these ERVs belong to either gammaretroviruses (env-Tac1, -Tac2, -Tac3, and -Tac4) or betaretroviruses (env-Tac5, - Tac6, and -Tac7) (Fig. S1). Nucleotide sequences of 5”-LTR and 3”-LTR were almost identical in the high-copy proviruses encoding *env-Tac3* (90.97 to 100%), *env-Tac5* (87.87 to 100%), and *env-Tac6* (82.82 to 99.53%) suggesting that they still actively expand in echidna genomes.

**Figure 1.**
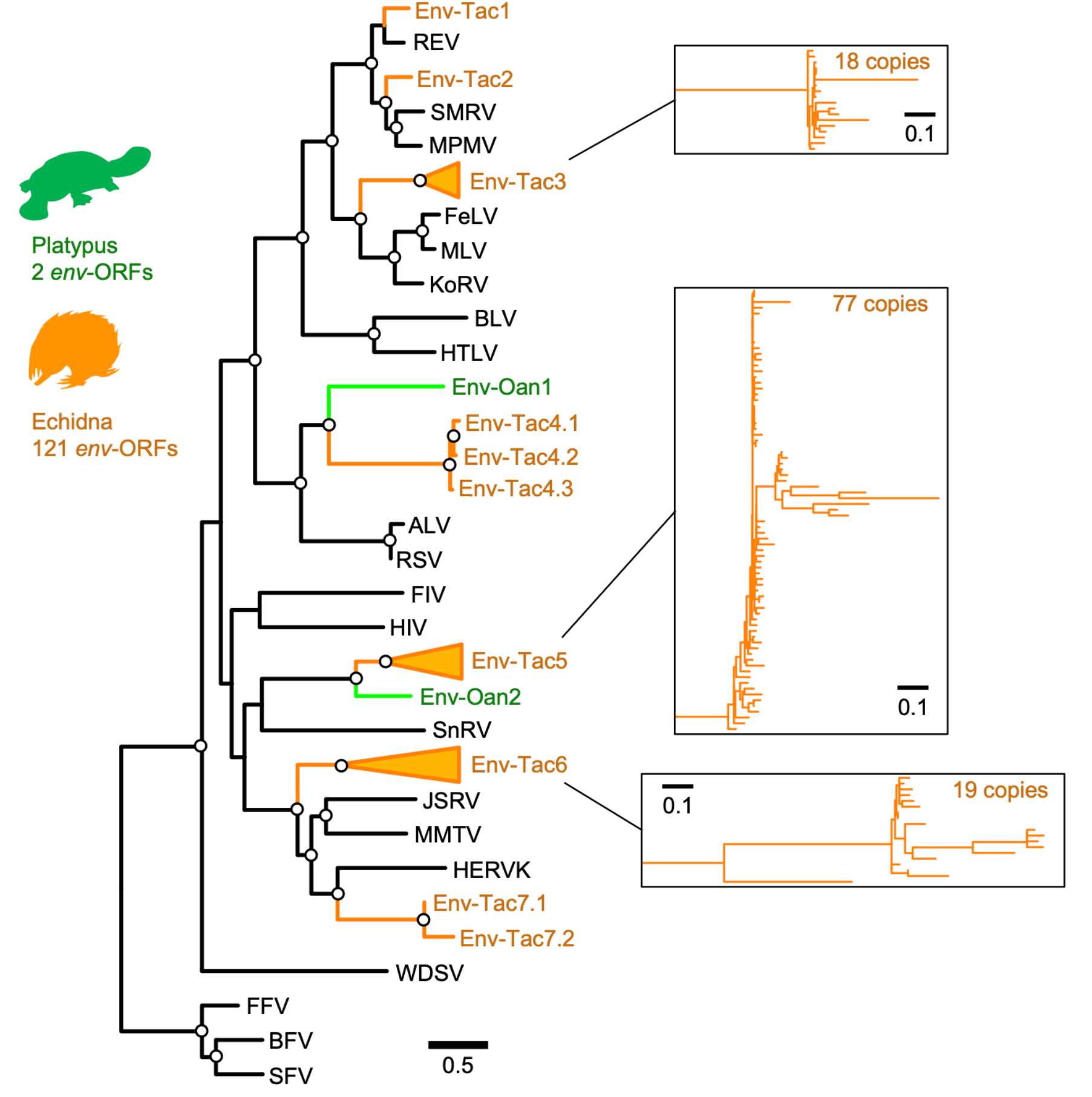
Phylogenetic tree of *env*-ORFs in the platypus and echidna genomes. Alignment of transmembrane regions of the Env proteins coded by *env*-ORFs and representative retroviral Env proteins (Table S1) are used. An open circle in an internal node indicates >95% ultrafast bootstrap support (1000 replicates).

**Figure 2.**
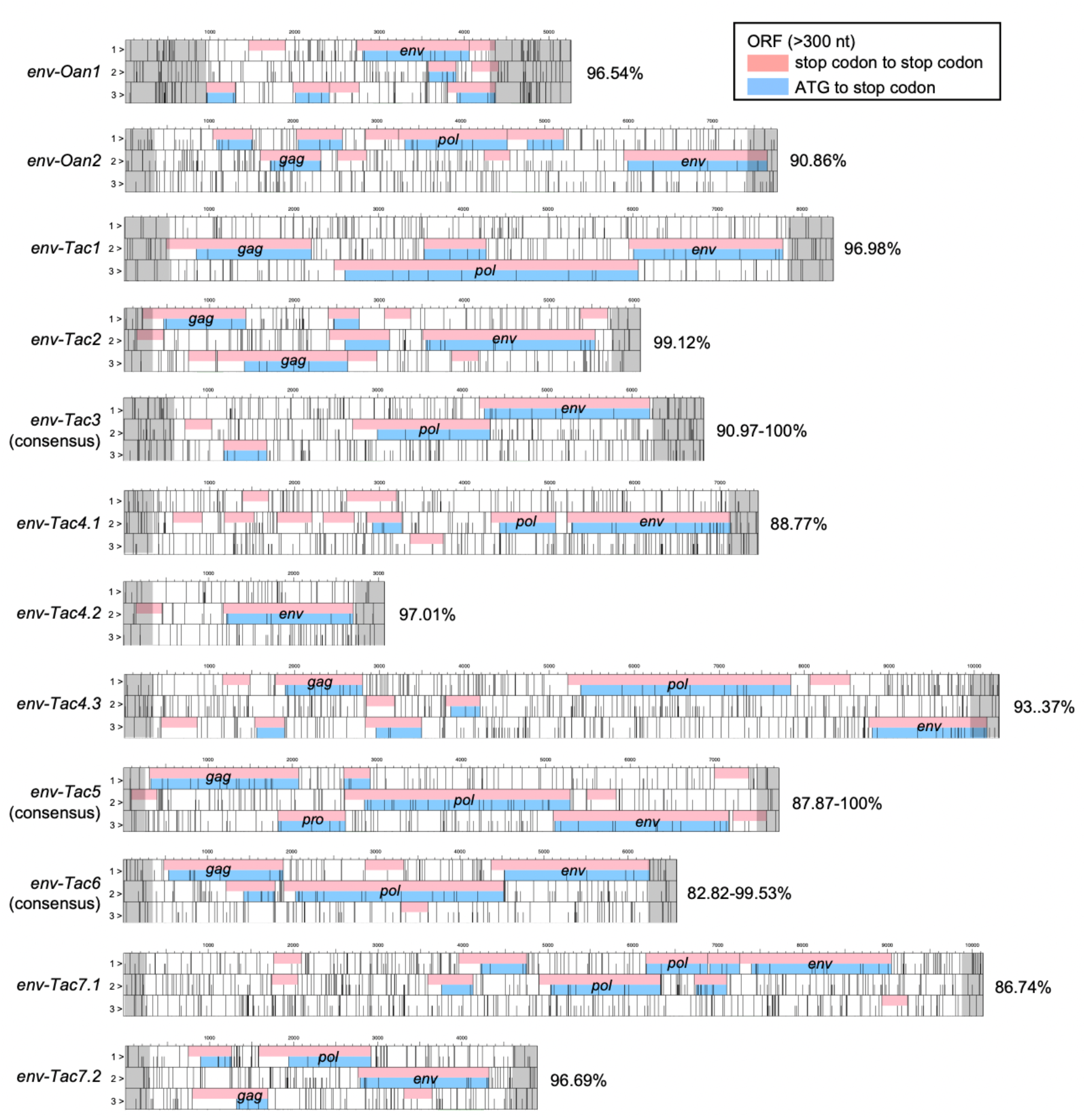
Proviruses encoding platypus and echidna *env*-ORFs. For each *env*-ORF, its retroviral structure, including *gag, pro*, and *pol* genes, was illustrated. Short bars indicate start codons, and tall bars indicate stop codons. ORFs of more than 300-nt were filled in magenta (a stop codon to a stop codon) or light blue (an ATG triplet to a stop codon). Retroviral gene names (*gag, pro, pol*, and *env*) were also indicated. 5”- and 3”-LTRs were filled with gray. Percent nucleotide identity between 5’ s- and 3’-LTRs was indicated on the right side of the sequences. For multi-copy *env*-ORFs (i.e., *env-Tac3, -Tac5*, and *-Tac6*), consensus sequences were constructed.

### Tissue expression of transcripts encoding *env*-ORFs

To investigate the transcription of *env*-ORFs, we calculated the expression levels of genes consisting of transcripts encoding *env*-ORFs from the platypus and echidna tissue transcriptome data (Fig. 3A). In the platypus tissues, we could not detect the transcripts of *env-Oan1* and *-Oan2*. In contrast, *env-Tac1, -Tac2, -Tac4*.*1*, and eight *env-Tac5* loci were expressed in echidna tissues (Fig. 3B). The *env-Tac1* showed the highest expression among these *env*-ORFs, whereas the *env-Tac2* was expressed only in male samples. When multi-mapped reads were included in the analysis, the expression levels of *env-Tac1* were increased, suggesting the presence of *env-Tac1*-like sequences in the echidna genome (Fig. 3C). In contrast, the expression level of *env-Tac4*.*1* did not change when uniquely mapped reads were used. All expressed sequences in *env-Tac5* genes showed specificity to the ovary (Fig. 3B). The expression levels of the *env-Tac5* genes, such as *env-Tac5*.*29*, were greatly increased when multi-mapped reads were used, suggesting that their sequences were similar (Fig. 3C). These transcriptome analyses revealed that *env*-ORFs are actively transcribed in echidna tissues and have different tissue specificities among *env*-ORF groups.

**Figure 3.**
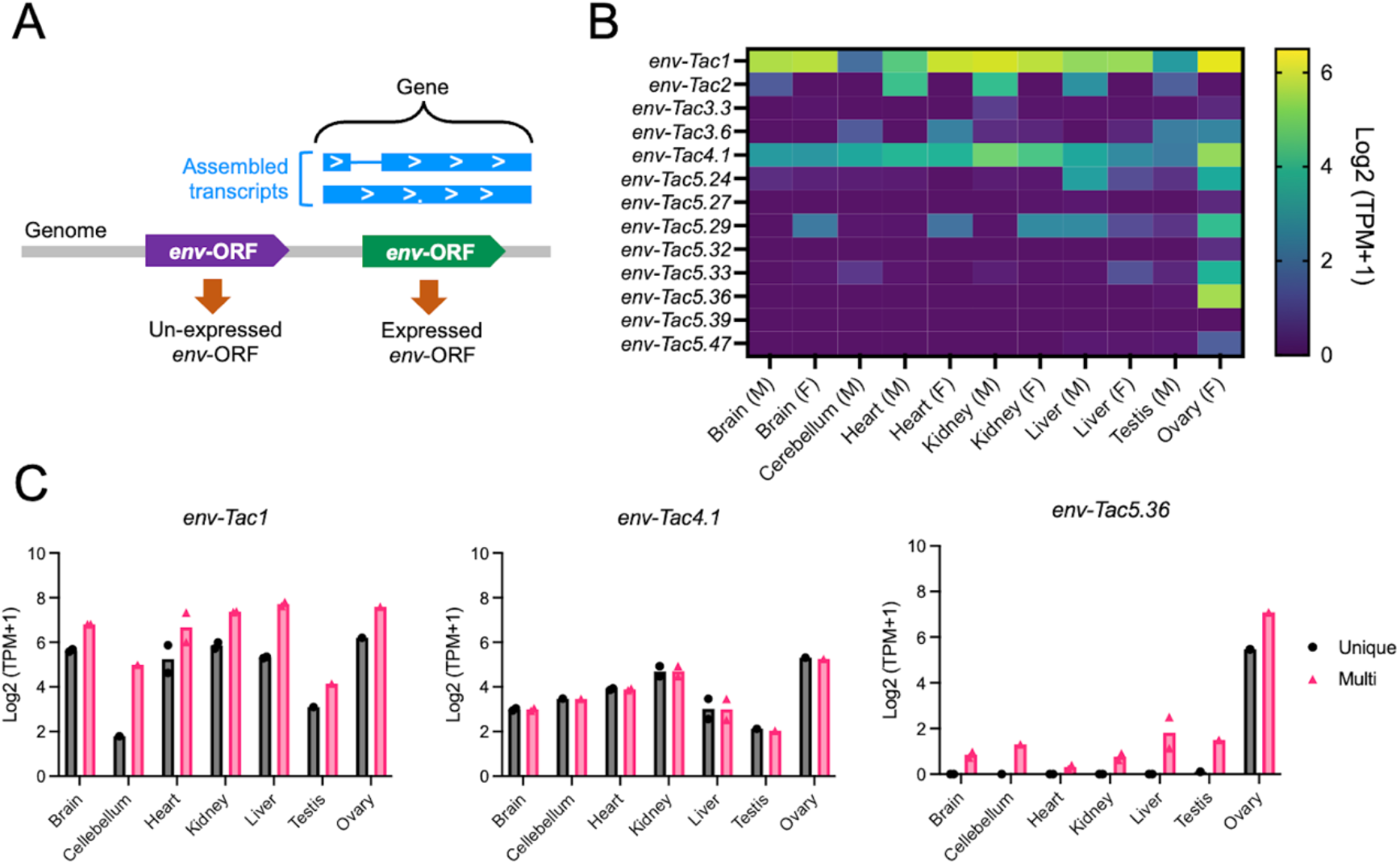
Tissue expression of echidna *env*-ORFs. (A) Tissue expression was calculated based on the genes of the assembled transcripts that overlapped with *env*-ORFs. (B) Heatmap rows indicate the expression levels of *env*-ORFs, and columns indicate echidna tissues. M, male; F, female. TPMs were calculated using uniquely-mapped reads. (C) TPMs with or without multi-mapped reads. Dots show individual values, and bars show the means.

### *env-Tac1* potentially encodes a functional Env protein

To further characterize the Env proteins identified in this study, we investigated *env-Tac1* which was highly expressed in various echidna tissues (Fig. 3B). The Env-Tac1 protein is closely related to reticuloendotheliosis virus (REV), which belongs to the RD114/D-type retrovirus (RDR) interference group (Fig 1; Fig. S2A). Previously, Niewiadomska and Gifford reported a REV-like *env*-ORF lacking the SU domain in the echidna genome, as detected by PCR (23). The *env-Tac1* was thought to be a full-length copy of this ERV group, which was suggested by our phylogenetic analysis of REV Env proteins (Fig. S2A). A multiple sequence alignment of the Env-Tac1 protein with the Env proteins of REV and spleen necrosis virus (SNV), which is a subgroup of REV, showed that the Env-Tac1 protein retains functionally important amino acids (Fig. S2B). In particular, the SDGGGXXDXXR motif is important for the binding to receptors, amino acid transporter ASCT1 (also known as SLC1A4) and/or ASCT2 (SLC1A5) (24). The predicted topology of the Env-Tac1 protein suggested a typical Env protein structure with a signal peptide at the N-terminus and a single transmembrane domain at the C-terminus (Fig. S2C). Together, the Env-Tac1 protein retained amino acid motifs for functional RDR Env protein.

### Env-Tac1 protein interacts with ASCT1 and ASCT2 to induce membrane fusion

The fusogenic activity of the Env-Tac1 protein was experimentally characterized under the assumption that the Env-Tac1 uses ASCT1 and/or ASCT2 as receptors. First, we conducted a cell-cell-fusion-dependent Lac Z assay to verify the cell-cell fusion activity of the Env-Tac1 protein in the presence of ASCT1 or ASCT2 (Fig. 4A). The Env-Tac1 protein was revealed to mediate cell-cell fusion in 293T cells in the presence of both human and echidna ASCT1 and ASCT2 (Fig. 4B). The fusion activity was the highest on echidna ASCT2. These experimental data support that the Env-Tac1 protein is the RDR Env protein that utilizes both ASCT1 and ASCT2 as receptors. To investigate whether the Env-Tac1 protein enables retroviral entry, we generated murine leukemia virus (MLV)-based pseudotypes with the Env-Tac1 protein carrying the LacZ marker (Fig. 4C). 293T cells stably expressing echidna ASCT1 and ASCT2 (293T-tacASCT1 and 293T-tacASCT2, respectively) were generated using a piggyBac system. We found that the Env-Tac1 protein mediated the infection of both 293T-tacASCT1 and 293T-tacASCT2 cells (Fig. 4D). The infectious units were higher in 293T-tacASCT2 cells than those in 293T-tacASCT1 cells, which is consistent with the higher cell-cell fusion activity on tacASCT2. Env proteins of SNV mediated infection only in 293T-tacASCT1 cells, suggesting that there are differences in amino acids that determine whether they are directed toward ASCT1 or ASCT2. We examined the expression of *ASCT1* and *ASCT2* in various echidna tissues and found that *ASCT2* was particularly highly transcribed in the kidney (Fig. 4E), in which *env-Tac1* was also expressed (Fig. 2). These data indicate that the Env-Tac1 protein is a potential fusogen in echidna tissues utilizing ASCT1 and ASCT2 as receptors.

**Figure 4.**
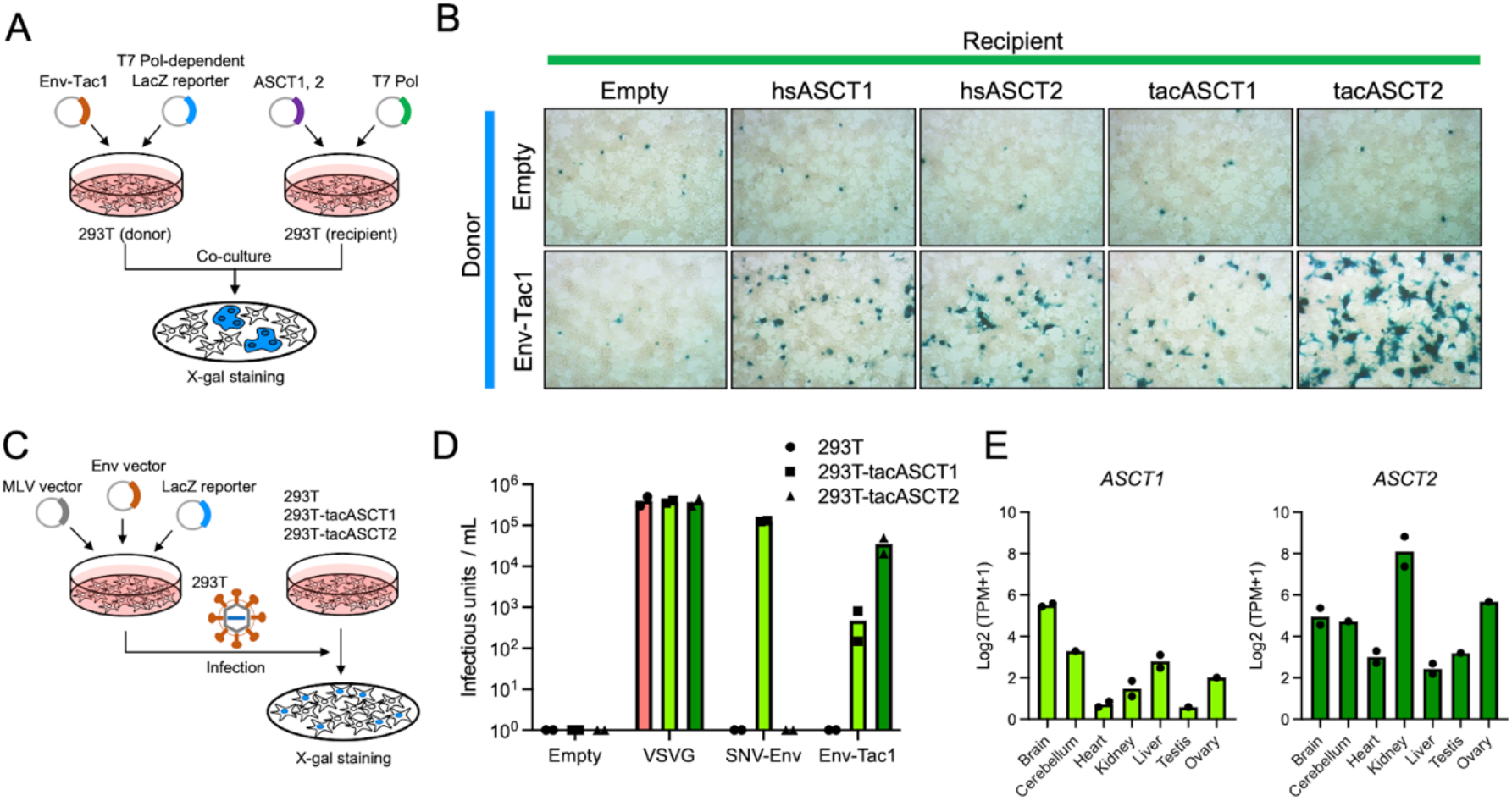
Fusogenic activity of the Env-Tac1 protein. (A) The cell-cell-fusion-dependent-LacZ assay. In the donor 293T cells, Env-Tac1-encoding plasmid or empty plasmid was transfected with the reporter plasmid, which expresses lacZ in the presence of T7 polymerase. In the recipient 293T cells, the plasmid encoding ASCT1 or ASCT2 of human or echidna was transfected with an expression plasmid for T7 polymerase. After co-culture of donor and recipient cells, fused cells were visualized by X-gal staining. (B) The Env-Tac1 protein caused cell-cell fusion in the presence of ASCT1 and ASCT2 of human (hsASCT1 and hsASCT2) and echidna (tacASCT1 and tacASCT2). The LacZ-positive fused cells were stained by X-gal 48 hours after transfection. (C) Human 293T cells expressing echidna ASCT1 and ASCT2 (293T-tacASCT1 and 293T-tacASCT2) were generated using the piggyBac system. The expression plasmid of VSVG, SNV-Env, Env-Tac1, or empty plasmid was transfected to 293T cells with the MLV virion and LacZ-reporter plasmids. Supernatants containing pseudotyped viruses were infected with 293T, 293T-tacASCT1, and 293T-tacASCT2 cells. (D) LacZ-positive focuses were counted and viral titers per milli-liter were calculated. (E) Tissue expression of echidna ASCT1 and ASCT2 from RNA-seq data.

## Discussion

The acquisition of the ERV-derived *env* genes has been associated with the evolution of placentation in therians (25, 26). The very small numbers of ERVs in the platypus genomes arose the hypothesis that intensive invasion of therian genomes may facilitate the co-option of ERV-derived placental genes (27). However, our study revealed that the echidna genome contained more than one hundred nearly full-length *env*-ORFs. The number of echidna *env*-ORFs was not unusual compared with that in other mammals. For example, 37 and 255 Methionine-starting *env*-ORFs longer than 400 codons were found in the human and mouse genomes, respectively (21), and 162 nearly full-length *env*-ORFs have been identified in nine-banded armadillo (28). Thus, the copy number of *env*-ORFs varies greatly among mammalian species, including the monotremes, during evolution.

Among echidna *env* genes, we demonstrated the cell-cell fusion activity of the Env-Tac1 protein. The systemic expression of *env-Tac1* mRNA raises the question of how such potentially harmful fusogenic genes are suppressed. One explanation is that the Env-Tac1 protein is not translated because *env*-ORFs, including *syncytin-1* and *syncytin-2*, require specific RNA motifs to be translated (29). Another possibility is that restriction protein(s) suppress cell-cell fusion. One candidate is defective Env proteins interacting with ASCT1 and ASCT2. Suppressyn, a truncated form of the RDR Env protein encoded in ERV-Fb1, inhibits cell-cell fusion by the Syncytin-1 protein (30). The balance between Syncytin-1 and Suppressyn expression regulates human placental formation (31). Therefore, suppressive truncated env genes may give tolerance to the fusogenicity of the *env-Tac1* protein.

Identifying the evolutionary conservation of retroviral genes is necessary for determining their co-options on host functions. Previously, we reported retroviral reverse transcriptase-derived genes *RTOM1, RTOM2*, and *RTOM3*, which are conserved in platypus and echidna (20). The dN/dS analysis revealed that these genes were under purifying selection between the two species. In this study, however, no orthologous *env* genes were found in platypus and echidna, indicating that *env* genes are not conserved in monotremes. Given that most of the ERVs encoding *env*-ORFs still retained retroviral ORFs other than *env* genes (Fig. 2), these *env*-ORFs could be under the process of mutational decay. One exception is *env-Tac4*.*1*, whose provirus has lost ORFs of *gag* and *pol* genes and showed low LTR similarity, suggesting purifying selection on the encoded *Env* protein (Fig. 2). Future comparative analyses of the evolutionary conservation of individual *env* genes in other echidna species, whose genomes have not yet been sequenced, will reveal the evolutionary conservation of individual *env* genes. In conclusion, we demonstrated the diversity of *env* genes in monotremes and identified an *env*-derived gene in echidna showing cell fusion ability, which indicates the dynamic evolution of *env*-derived genes in mammals.

## Materials and Methods

### Detection of *env*-ORFs

The platypus genome (mOrnAna1.p.v1, GCF_004115215.1) and echidna genome (mTacAcu1.pri, GCF_015852505.1) were used for the *env*-ORF screening. The ORFs of more than 400 codons from start to stop codons were retrieved using the getorf program in the European Molecular Biology Open Software Suite v6.5.7.0 (32). To obtain ORFs similar to *env* genes, hmmscan in HMMER3 v3.2.2 (33) was used (expected threshold: 1E-10). The profile-HMMs of retroviral Env proteins retrieved from the GyDB collection (https://gydb.org/index.php?title=Collection_HMM) were used as queries (34).

### Phylogenetic analysis

To infer the phylogenetic relationship of monotreme *env*-ORFs obtained in this study, amino acid sequences of 20 retroviral Env were retrieved from representative retroviruses (accession numbers are listed in Table S1). In total, 153 amino acid sequences were aligned using the MAFFT v7.487 with the L-INS-i method (35). Conserved transmembrane (TM) domains of the Env proteins (regions after furin cleavage RXR/KR motif) were extracted. For the Env-Tac1 phylogeny belonging to the RD114/D-type retrovirus (RDR) interference group, we generated a multiple sequence alignment of full-length amino acid sequences of the Env-Tac1 and RDR Env proteins (Table S1) using the same procedure. IQ-TREE2 v2.0.8 (36) was used to construct the phylogenetic tree with 1000 replicates generated by an ultrafast bootstrap approximation (37). The tree was visualized using FigTree v1.4.4 (https://github.com/rambaut/figtree/releases).

### Characterization of proviruses of *env*-ORFs

The nucleotide sequences of *env*-ORFs with 10 kbp downstream and upstream extensions were retrieved using BEDtools v2.30.0 (38). 5”- and 3”-LTRs were identified using LTRharvest (39) and LTRdigest (40) included in GenomeTools v1.6.2. To obtain consensus sequences for the multicopy *env* genes (*env-Tac3, env-Tac5*, and *env-Tac6*), the proviral sequences were aligned with MAFFT as previously described. Then minor insertions were removed using T-coffee v11.0.8 (41) with the “rm_gap 60” option. A consensus sequence for each *env* group was generated using the cons program in the European Molecular Biology Open Software Suite v6.5.7.0 (32). The ORFs in proviruses were visualized using ApE v3.0.6 (42). Characterization of proviral ORFs was based on the protein motif search in the HMMER web server (43).

### Analysis of transcriptome data

RNA-seq data of platypus (20 samples from 6 tissues) (44) and echidna (11 samples from seven tissues) (19) were obtained from the SRA database whose Accession IDs were summarized in Table S2. Low-quality reads were trimmed and filtered using fastp v0.19.5 with default options (45). The filtered reads were mapped to each reference genome using HISAT2 v2.1.0 (46). For each RNA-seq dataset, we conducted genome-guided transcript assemblies using StringTie2 v2.1.6 (47) with “--rf” options to specify the strandedness. The resultant GTF files and the RefSeq GTF file were merged using StringTie2 with the “--merge” option to generate a merged GTF file. To quantify the expression level of each *env*-ORF, we calculated transcripts per kilobase million (TPM) for 20 platypus and 11 echidna RNA-seq samples using the Stringtie2 program v2.1.6 with “-e --rf” options (47). Exons in the merged GTF file overlapped with *env*-ORFs were identified using BEDtools intersect with “-s” option to specify the strandedness (38), and TPM of genes whose exons were overlapped with *env*-ORF were regarded as expression levels of *env*-ORFs. For extraction of unique-mapped reads, we collected reads that had “NH:i:1” flag, which indicates they were uniquely mapped, in aligner-generated SAM files. A heatmap of TPMs was generated using GraphPad Prism v9.4.1. Source data are included in Dataset S3.

### Cell culture

293T cells (#RCB2202, Riken BioResource Research Center, Tsukuba, Japan) were cultured in Dulbecco”s modified Eagle”s medium (#D5796, Merck, Darmstadt, Germany) supplemented with 10% heat-inactivated fetal calf serum (#10270106, Thermo Fisher Scientific, Waltham, MA, USA), and penicillin (100 units/mL) and streptomycin (100 mg/mL) (#09367-34, Nacalai Tesque, Kyoto, Japan) at 37°C in a humidified atmosphere of 5% CO2 in the air.

### Plasmids

To construct the Env-Tac1 expression plasmid (phCMV3-Env-Tac1), dsDNA encoding the *env-Tac1* ORF was synthesized by Eurofins Genomics (Tokyo, Japan). The synthesized dsDNA was inserted into EcoRI and BamHI sites of phCMV3 (#P003300, Genlantis, San Diego, CA, USA) using NEBuilder HiFi DNA Assembly Master Mix (#M5520AA, New England Biolabs, Ipswich, MA, USA). For the construction of the SNV-Env expression plasmid (phCMV3-SNV-Env), the SNV-Env coding region was amplified from pPR102 (48) using PCR and inserted into the EcoRI and BamHI sites of phCMV3. To construct the human ASCT1 and ASCT2 expression plasmids (phCMV3-hsASCT1 and phCMV3-hsASCT2), human ASCT1 and ASCT2 were amplified from 293T cDNA and inserted into the the EcoRI and BamHI sites of phCMV3. For the construction of the echidna ASCT1 and ASCT2 expression plasmids (phCMV3-tacASCT1 and phCMV3-tacASCT2), dsDNA fragments coding the ASCT1 and ASCT2 were synthesized based on the RefSeq sequences of XM_038750847.1 (ASCT1) and XM_038766835.1 (ASCT2) (Eurofins Genomics K.K., Tokyo, Japan) and inserted into the EcoRI and BamHI sites of phCMV3. For the construction of the echidna ASCT1 and ASCT2 expression piggyBac plasmids (pPB-tacASCT1 and pPB-tacASCT2), the GFP coding region of the piggyBac plasmid (#VB900088-2265rnj, VectorBuilder, Chicago, IL, USA) was removed by inverse PCR and was replaced with the echidna ASCT1 and ASCT2 fragments using NEBuilder HiFi DNA Assembly Master Mix. All PCRs described above were carried out using KOD One Master Mix (#KMM-101, TOYOBO, Osaka, Japan) with a C1000 Touch thermal cycler (Bio-Rad, Hercules, CA, USA). Primer sequences are listed in Table S3.

### Construction of echidna ASCT1 and ASCT2 expressing 293T cells

293T cells were seeded in 24 well plates at 2.5 × 105 cells per well. The next day, 0.4 µg of pPB-tacASCT1 or pPB-tacASCT2 and 0.1 µg of the hyper PBase expression plasmid (#VB900088-2874gzt, VectorBuilder) were transfected into 293T cells. Forty-eight hours after transfection, 293T cells were selected with 1 µg/mL of puromycin. After passaging for more than 7 days, the selected cells were termed 293T-tacASCT1 and 293T-tacASCT2.

### Pseudotyping assay

Pseudotyped viruses were produced by transfection of 2 × 106 293T cells in a 35-mm dish with 0.7 µg of the plasmid encoding MLV Gag and Pol proteins (pGag-Pol-IRES-bsr) (49), 0.7 µg of retroviral plasmid for nuclear localization signal-attached LacZ (pMX-nlsLacZ), and 0.7 µg of Env expression plasmid (phCMV3, pCAG-VSVG, phCMV3-SNV-Env, or phCMV3-Env-Tac1) using 2.8 µL of Avalanche Everyday Transfection Reagent (#EZT-EVDY-1, EZ Biosystems, College Park, MD, USA). Forty-eight hours after transfection, supernatants were filtered through 0.45 µm polyvinylidene difluoride (PVDF) membranes and transferred to target cells, which were seeded in 96-well plates the day before infection, with 8 µg/mL Polybrene. X-gal staining was performed two days after infection.

### Cell-cell fusion-dependent-LacZ assay

Donor 293T cells, which were seeded at 2.5 × 105 cells per well in 24-well plates, were transfected with 0.5 µg of phCMV3 or phCMV3-Env-Tac1 and 0.5 µg of the LacZ reporter plasmid (pT7EMCV-LacZ), which expresses LacZ with internal ribosome entry site from encephalomyocarditis virus in the presence of T7 polymerase. The recipient 293T cells were transfected with 0.5 µg of receptor expression plasmid (phCMV3, phCMV3-hsASCT1, phCMV3-hsASCT2, phCMV3-tacASCT1, or phCMV3-tacASCT2) and 0.5 µg of expression plasmid for T7 polymerase (pCAG-T7Pol). For transfection, 0.8 µL of Avalanche Everyday Transfection Reagent was used. Donor and recipient cells were co-cultured six hours after transfection, and the LacZ-positive fused cells were stained by X-gal 42 hours after co-culture.

## Acknowledgments

This work was supported by Grant-in-Aid for JSPS fellows 20J22607 to K.K. and JSPS KAKENHI 20K06775 and 20H03150 to T.M. and S.N. The super-computing resource was partially supported by the NIG supercomputer at ROIS National Institute of Genetics.

