## Supplementary Figures and Tables for "Dynamic evolution of retroviral envelope genes in egg-laying mammalian genomes"

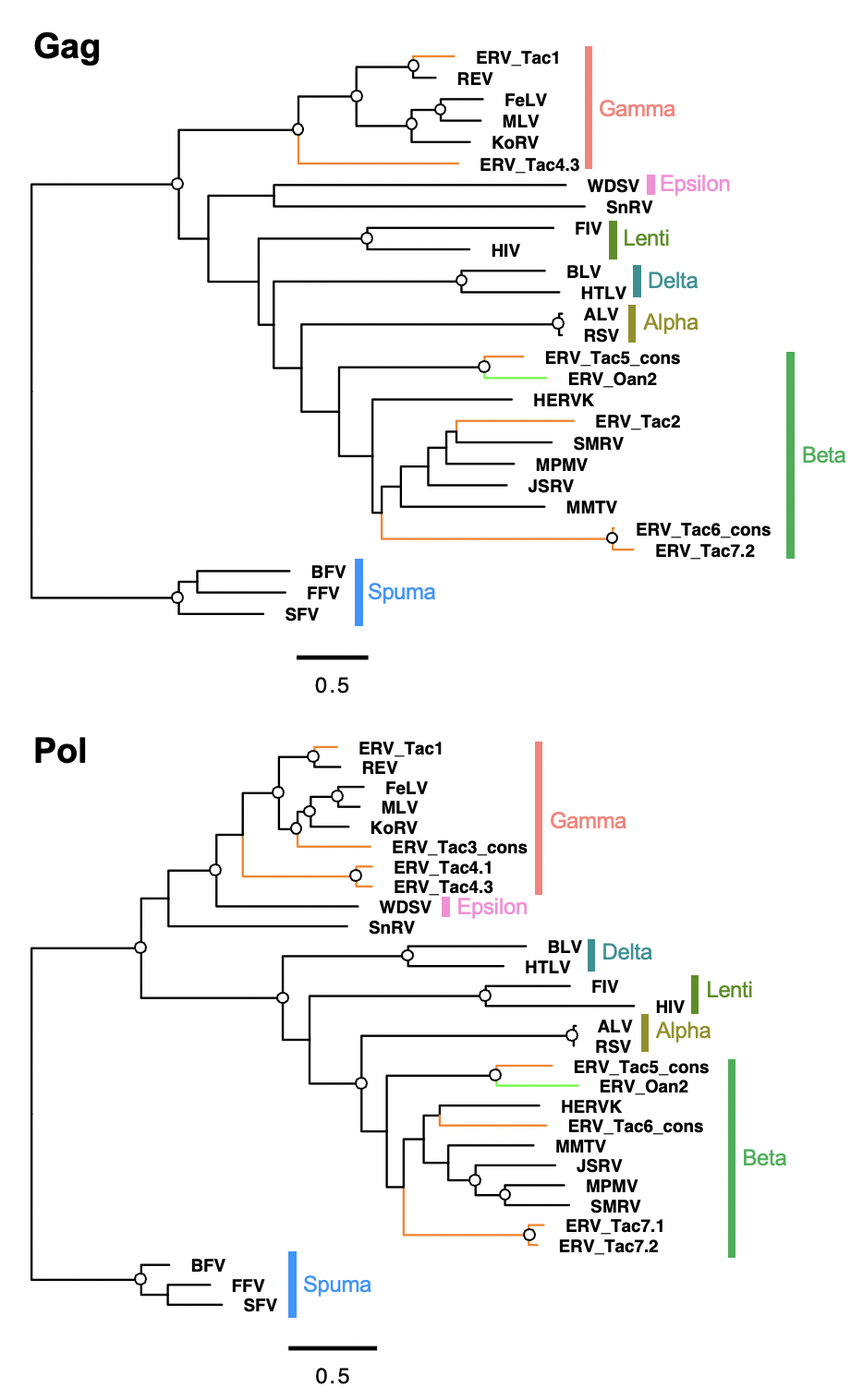

Fig. S1. Phylogenetic trees of Gag and Pol proteins encoded in proviruses. The alignment of Gag and Pol proteins encoded in provirus (Fig. 2) and representative retroviruses (Table S2) were used for tree construction. An open circle in an internal node indicates >95% bootstrap support (1000 replicates).

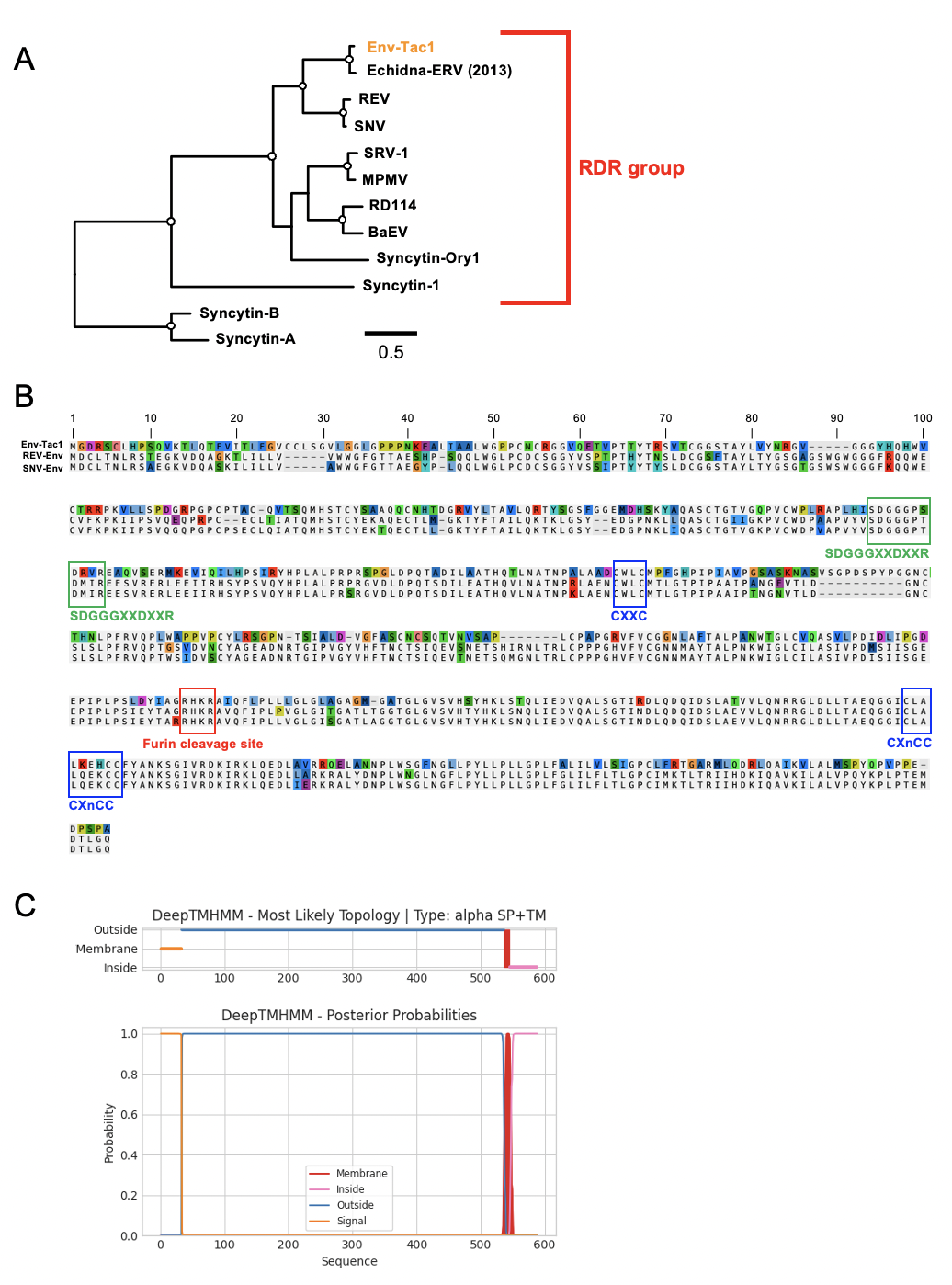

Fig. S2. Characterization of the Env-Tac1 protein. (A) A phylogenetic tree of RDR Env proteins with mouse Syncytin-A and B as an outgroup. An open circle in an internal node indicates >95% bootstrap support (1000 replicates). (B) Multiple alignment of amino acid sequences of the Env-Tac1, REV-Env, and SNV-Env. Functionally important motifs were indicated with captions. (C) Predicted topology of the Env-Tac1 protein. The Env-Tac1 proteins have the N-terminus signal peptide and a transmembrane domain like other retroviral Env proteins.

Table S1. Representative retroviruses used for phylogenetic trees.

| Name | Abbreviation | Accession number | Phylogenetic tree |
| --- | --- | --- | --- |
| Rous sarcoma virus | RSV | NC_001407.1 | Fig1, Fig S1 |
| Avian leukosis virus | ALV | NC_001408.1 | Fig1, Fig S1 |
| Jaagsiekte sheep retrovirus | JSRV | NC_001494.1 | Fig1, Fig S1 |
| Mouse mammary tumor virus | MMTV | NC_001503.1 | Fig1, Fig S1 |
| Human endogenous retrovirus K113 | HERVK | NC_022518.1 | Fig1, Fig S1 |
| Reticuloendotheliosis virus | REV | NC_006934.1 | Fig1, Fig S1 |
| Squirrel monkey retrovirus | SMRV | NC_001514.1 | Fig1, Fig S1 |
| Mason-Pfizer monkey virus | MPMV | NC_001550.1 | Fig1, Fig S1 |
| Koala retrovirus | KoRV | NC_039228.1 | Fig1, Fig S1 |
| Moloney murine leukemia virus | MMLV | NC_001501.1 | Fig1, Fig S1 |
| Feline leukemia virus | FeLV | NC_001940.1 | Fig1, Fig S1 |
| Human T-lymphotropic virus 4 | HTLV | NC_011800.1 | Fig1, Fig S1 |
| Bovine leukemia virus | BLV | NC_001414.1 | Fig1, Fig S1 |
| Feline leukemia virus | FIV | NC_001482.1 | Fig1, Fig S1 |
| Human immunodeficiency virus 1 | HIV | NC_001802.1 | Fig1, Fig S1 |
| Walleye dermal sarcoma virus | WDSV | NC_001867.1 | Fig1, Fig S1 |
| Snakehead retrovirus | SnRV | NC_001724.1 | Fig1, Fig S1 |
| Simian foamy virus | SFV | NC_001364.1 | Fig1, Fig S1 |
| Feline foamy virus | FFV | NC_039242.1 | Fig1, Fig S1 |
| Bovine foamy virus | BFV | NC_001831.1 | Fig1, Fig S1 |
| Echidna ERV | - | AGV92857.1 | Fig S2A |
| Simian retrovirus 1 | SRV-1 | AAA47733.1 | Fig S2A |
| Mason-Pfizer monkey virus | MPMV | AAA47712.1 | Fig S2A |
| RD114 retrovirus | RD114 | YP_001497149.1 | Fig S2A |
| Reticuloendotheliosis virus | REV | YP_223872.1 | Fig S2A |
| Spleen necrosis virus | SNV | AAZ57419.1 | Fig S2A |
| Baboon endogenous virus | BaEV | YP_009109691.1 | Fig S2A |
| syncytin-1 | - | NP_055405.3 | Fig S2A |
| syncytin-A | - | NP_001013773.1 | Fig S2A |
| syncytin-B | - | NP_775596.1 | Fig S2A |

Table S2. RNA-seq datasets used for expression analyses.

| Species | Tissue | SRA_ID |
| --- | --- | --- |
| *Tachyglossus aculeatus* | Liver | SRR10530483 |
| *Tachyglossus aculeatus* | Kidney | SRR10530484 |
| *Tachyglossus aculeatus* | Testis | SRR10530485 |
| *Tachyglossus aculeatus* | Heart | SRR10530486 |
| *Tachyglossus aculeatus* | Brain | SRR10530487 |
| *Tachyglossus aculeatus* | Kidney | SRR10530488 |
| *Tachyglossus aculeatus* | Ovary | SRR10530489 |
| *Tachyglossus aculeatus* | Liver | SRR10530490 |
| *Tachyglossus aculeatus* | Cerebellum | SRR10530491 |
| *Tachyglossus aculeatus* | Heart | SRR10530492 |
| *Tachyglossus aculeatus* | Brain | SRR10530493 |
| *Ornithorhynchus anatinus* | Brain | SRR5412222 |
| *Ornithorhynchus anatinus* | Brain | SRR5412223 |
| *Ornithorhynchus anatinus* | Brain | SRR5412224 |
| *Ornithorhynchus anatinus* | Brain | SRR5412225 |
| *Ornithorhynchus anatinus* | Heart | SRR5412226 |
| *Ornithorhynchus anatinus* | Heart | SRR5412227 |
| *Ornithorhynchus anatinus* | Heart | SRR5412228 |
| *Ornithorhynchus anatinus* | Heart | SRR5412229 |
| *Ornithorhynchus anatinus* | Kidney | SRR5412230 |
| *Ornithorhynchus anatinus* | Kidney | SRR5412231 |
| *Ornithorhynchus anatinus* | Kidney | SRR5412232 |
| *Ornithorhynchus anatinus* | Kidney | SRR5412233 |
| *Ornithorhynchus anatinus* | Liver | SRR5412234 |
| *Ornithorhynchus anatinus* | Liver | SRR5412235 |
| *Ornithorhynchus anatinus* | Liver | SRR5412236 |
| *Ornithorhynchus anatinus* | Liver | SRR5412237 |
| *Ornithorhynchus anatinus* | Ovary | SRR5412238 |
| *Ornithorhynchus anatinus* | Ovary | SRR5412239 |
| *Ornithorhynchus anatinus* | Testis | SRR5412240 |
| *Ornithorhynchus anatinus* | Testis | SRR5412241 |

Table S3. Primers.

| Purpose | Template | Primer sequence (5' to 3') | Direction |
| --- | --- | --- | --- |
| SNV-Env for phCMV3-SNV-Env | pPR102 | CGAGCTCAAGCTTCGCCACCATGGACTGTCTCACCAAC | Forward |
|  |  | ACATCGTATGGGTAGTCATTGACCTAGGGTATCCATC | Reverse |
| Human ASCT1 for phCMV3-hsASCT1 | Human cDNA | CGAGCTCAAGCTTCGCCACCATGGAGAAGAGCAACGAG | Forward |
|  |  | ACATCGTATGGGTAGAGAACCGACTCCTTGGATTCCAG | Reverse |
| Human ASCT2 for phCMV3-hsASCT1 | Human cDNA | CGAGCTCAAGCTTCGCCACCATGGTGGCCGATCCTCCT | Forward |
|  |  | ACATCGTATGGGTAGGCCATGACTGATTCCTTCTCAGAG | Reverse |
| piggyBac backbone | pPB-EGFP-Puro | ACCCAGCTTTCTTGTACAAAG | Forward |
|  |  | GGTGGCAGCCTGCTTTTTTGTAC | Reverse |
| Echidna ASCT1 for pPB-tacASCT1 | Synthesized DNA | AAGCAGGCTGCCACCATGACCTACGTTAGTC | Forward |
|  |  | ACAAGAAAGCTGGGTTTAAGCGTAATCCGGAACATC | Reverse |
| Echidna ASCT2 for pPB-tacASCT2 | Synthesized DNA | AAGCAGGCTGCCACCATGGAAGCCAGTCC | Forward |
|  |  | ACAAGAAAGCTGGGTTTAAGCGTAATCCGGAACATC | Reverse |

Dataset S1 (Excel file). Envelope-derived ORFs in the platypus and echidna genomes.

Dataset S2 (Word file). Annotated provirus sequences.

Dataset S3 (Excel file). TPMs of env-ORFs (source data of Fig. 3).
