## Supplementary material for "Dynamic evolution of retroviral envelope genes in egg-laying mammalian genomes": Dataset S2

**Dataset S2. Annotated provirus sequences.**

LTRs were underlined and env-ORFs were colored in blue.

＞ERV_Env-Oan1

TGTGGGAGTGTGCCAAGGTGGACCTGGGAACTGGCCACCGAAAGGTCAAAATGCCATCAGTGACGTAACCTCTCTGGCCAGAGGAGGCAGGACGGGAAGATAGGTGTCCCTCACCTCCGTGCCCACCCAATAAGGTATGGAAGGGACGGGAGCGGAAGGGGGCGGAGATTGAACAGGAAGGAAGTGGGGACCACCCAAACCAAGGTAATAAATACCTATTGCCTCTGGTGTTCGGGTCTCTCTCTGGGAAGATCACAGCAATAGCAACGGCCACCCACGTGTCTCTCCCTGTCGCGAGCACCAGACGCTATTTTGCCACCAGGAACAACGCACAGAGACAGGGGCCGCGGGACGGGTGAGTGTAACACATAACTGGGTTAAATACATAAGTGGGTAATGTATGAGTGTAACGCATATGTGAGCTTAGTGTATAAGCGAGCAATGTATAAGTGAGTATAATGCACATGTGAGTTTAGTGCGTAAGTGGGTAGTGTACAAGCGAGTGTAACGCGCAAGTGAGTTTAACGCACGAGTGTGTAACGCATAAGTGAATCTAAAGCCTAAATGGGGTCGGCGCATAAGTAGATTGAACGCTTAAGTGGATTCCTCCCAAGTGGGTGCGCACATGGTGGGAACTTGCCGCCTCACCACGTTGTTAGTAAACCATTAACGAGCTCTTCCTGGGCGGGCCCTGGTCTGCCCACACTCGAGAAACATACTAGTGTAATCCTCCCATCAGGGAAGCTTTAACACCAAATTCCTCCCAAAGGGGAAGGTTTAACTCCATACTTTCCCCAAAAAGGGAAGGCCGACTGTCGCGTCTAAATAATTAATTAATTCAAACCGCCCCGTGGAATAAATTCTATACAAAACTCAGGTTTTCCATCTCCAGCCTCTCTCTCTCTCTCTCCCGCGCCGATTCCGAAAGAACCTGTCCCCGGCAACGGGTGACATAAATGGCGACGAGGATGGGATCGGCGATAGGCGATAGAGAGCCGGGAGGGCTATTTGAATTCCTTGCGAATCAGGGATGGAAAACCCCGAGATGGGACAGCCGCTGGGTGGAGGACCATTGGGAAGATCCGGCCAATCTGGTGAGGGAGTTTTGTAATTGGCGTGAGGCTACCCAAATCAGGAAAAGGAAAGGAAAGACCACCCTTCTCGGCCTCCTAGCAACAGCGCTGCAGAGCGCTGCAGGCCGCCGCCCAGGATCGGCGGCGGTTGCGGCAGGAAAGGGACGCCCTGGAGGCGAAACTAAGGGAACGGGACACGAGATGCCTCTTATTGCAGACAGCGGCTGAGGTTAAAGCACAGCAGAACACCCTTTTGGAGCAGAAATTGGCACCCCGGGTGGCCAAGGAGTTAGAGGAGAGAAGAAACCTTCCGGTAACCGAGCAGGAGGTTAAGCAAGTCATGTACTCGGGAATTTGCCAGGGGAATTGCTCAGGGGAAGGGTAGGGAGATCTCGGCCCTTTACCAGAAGTGGAACCGCCCCCTCCCCAGCAGGATCGGGTGGTGCCCCCGCAGAATCCCGAGTGCCAACCCAAGCCCATCTACCCTATTATCAAGGAGGAGCGGAACCCTGAGGGAAAGGTCACCACCCAGACTATGGGCTTCACAGCCCCTGAGCTCAGGCAGATCTCTAAGGAGTTTTCCCGGGACCCCGGGGAACCGGTGATAACCTGGCTGGCTAGGATTTGGGAGGGAGCAGCCGCTCAGGTAGACCTCAGCCCAAATGAAAGCAGCCGCCTAGATTTGGGTCCAGGGGTGGATGTATCCCACACTCACAAGGTCGAACTCTTTGGACCCTCGCGGTACATTGGGCTCGGACGATACCCTGCAGTGAACAAGGAAGGGCCGGCACTCTCCGGTGGACCGACCCAACCACACTAATCCAGCATCTCCTGGTGCTTGGGATCGAACAAGTCCTGGATGATAGGGGGAAAACTCAGGAACCTAGTCCATGGTCCCTACCCGCCTAAGAGGGTTTACTTCATGCCATAGTAATGGGTGCACCCCCGGGGCCAGAGCATGCTCTGGCTCACCGCTTGAGGGATAGAGCCACAGGAGCAAGCACCTGGAGCACATTGGCCAATGTGGTCAATAGTGACTGGCCCCTATTTCAGTCCTCAGAGGTACCCATTACCCGGCGTGTAGCAACCATGATTGGGAAGGGCCTAGGTTTGAGGAGGACAAAGTCAGCCCCAACATATACTAGAAGGAACACCCGCCAACCTCCGAAGTGGATCGGCTCAGGAAGACTGGGCTTAGGACGGGTAGCTATTTGGCGGGAGTTGAGGGAACGGGGCTTTCCCATGGATGAGTTAGATGGACTACCCACCCATATCCTGAAACTGCTCACTCTGGAAGGTAGACCATGGGTCCCAACCATCCCGGAGGAGGAAGACTAGGAGGGAGATCCGGCAGGCCGGGCTAATGGACGGTGGCCGCGGCCCCCTTACAGGCCTATGACCGTCCTAATGGATGACCGCCAGGAAGTCTCTTTCCTGGTAGATACCGAGGCCCAGATCCTCCACCTGGGATGGGGCTCAGCACATCTTTACATAGCTCCTACAGGGATACCCTCCCCTGAGAAGGATACTGGAGACAGATTGGGAAGCGAGTCATTATCCGGCATTCCCCAAAAGGCCAGTTATGGGTTTACCCTTATGAGAGCTCTCATGGGACACATTTGGGCAGCTAATACCAATGAGGGCTGCCCACTAACCCTGGTAGAGGCTTGGACATCCTCCAAGACCTAAGCTCTTCTCTTTGGCAGATACCACACCTGGGAGAGGGGAAATGCCGACCCAGCGCTCGATGAATCTGCTCAGCCTGCTTGTCATTCTGACCCTGGAATGTGGGAGAGCCCGCGGCAACACCTTCTTGAAGGCCTTGGCCCACTACAGCCAGGACCTGAATGTTTCAAACTGTTGGGTTTGTGGGCATTCCCCATTGAATGGAGGGGGAGGGTACCCCCTTATAGCCCATCCTTATGACAACTCCACCTGGTTCAATCCAACGGGAGCCCCCCATCCCTGCAGCACCCGCCCTGTCACGGAGCCCGCAAATGCCACCACATACTTTATGGTGAGTGGGTGGACTGACCATGCCGTGTTCCCATTCTGCTTTCACAGTACTGGTACCAAATTCTGGGCAGGCAATAGTTCCCATGTATGTGACGTAGCTAAGCTCCAGGGCGGGCCCAGATCTACGGCCGGACATTGTGGCCCTCCGACCACCAAGCAGGGGGTGGATGTCCGCTCCCATGGCCTGCTCATTGCGAATGGGACCTTTGGCAACTGTGGCACCATCCCCTGTGCAGTACCTGCCTATCTAATTTGTGGCGTCCGTGCCTATTGGTGGCTCCCTGCCGGCTGGGGAGGGACATGCTTCCTTGGTTTTGTACTCCCAGCTATCCATCACACCCTGAGTCGCCCGGAAGATCAACTCCGAAACCGTCGGGGGGCCCCAATCTCAGAGTCTAAATGCTTCTTCGGGATTATGTTTCCTGCATATGGGTTTGCACGGGCGGCTCATGAAATACTTAACCTAGCTAAACTAATTGAACAGATAACCAATGACACTGCTGTCAGCCTTATGGCCCTCAGGGACGAACAGAAAGCCATCTGGACGACCGTCCTTCAGAACCGAATGGCCCTTGACTTCCTCCTCGCCAGCCAAGGAGGGGTCTGCAAGCTTATCGGGAAGGAGTGTTGCACCTTTATCCCTGACAATTCTGGACATGTGGACGCAATTGTTGCTGATATGTACCACGCAGTTAACCAGTACCGGAATGATGACACTGCTGGAGGAGTCTGGGACTGGTTCCAAGGACTCTTTACGAACTGGGGGAGCTCACTATTCCATGGGTTGCTGCTGCTTTCGCTCTTGTTGGTGGGCATAGTGGGGGGATGCTGGTTCCTAGGTTGCTGTTGCTCGGTCCTCTCTACATGCATGCGACGAACACTGAAGTCATCCCCGACTCCTCTCTCTGTGCATTACCTCAAGAATGTTGTAGTCCACCAGTCACCCTCCCTACCTCTGGTCACCACACCTTAAAGGCTCCGCCTCCCCCCCCCCGTCCTAAGTCCCTGGCCAGAGCACGGGAGCTGTCCTCTGCCCCGACCTATGGATCTACTCTCTGGACTTCCGTGGCCAGCCAGGAGAGGATATCAGTTCCTTGTGGTCAGGGGAGGGACAATGTTGGAGGGCACGGCCTCCGGCATCACTACACCACCACGCGCCCCGGAGGGGATGCAGGGAAGGTTGGGGGTCAAATGGGCACGAATGGCGAGGGCCAAGGGGTGGTATGTGCCAAGGTGGACCTGGGCACTGGCTACCCAAAGGTCAAAATGCCATCAGTGACGTAACCTCTCTGGCCAGAGGAGGCAGGATGGGAAGATAGGTGTCCCTCACCTCCGTGCCCACCCAGTAAGGTATGGAAGGGAAGGGAGCGGAAGGTGGCGGAGATTGAACAGGAAGGAAGTGGGGACCACCCAAACCAAGGTAATAAATACCTATTGCCTCTGGTGCTCGGGTCTCTCTCTGGGAAGATCACAGCAGTAGCAACGGCCACCCACGTGTCTCTCCCTGTCGCGAGCACCAGACGCCATTCTGCCACCAGGAACAACGCACAGAGACAGGGGCCGCGGGACGGGTGAGTGTAACACATAACTGGGTTAAATACATAAGTGAGTAATGTATGAGTGTAACGCATATGTGAGCTTCGTGTATAAGCGAGCAATGTATAAGTGAGTATAATGCACATGTGAGTTTAGTGCGTAAGTGGGTAGTGTACAAGCGAGTGTAACGCGCAAGTGAGTTTAACGCACGAGTGTGTAACGCATAAGTGAATCTAAAGCCTAAATGGGGTCGGCGCATAAATAGATTGAACGCTTAAGTGGATTCCTCCCAAGTGGGTGCGCACATGGTGGGAACTTGCCGCCTCACCACGTTGTTAGTAAACCATTAACGAGCTCTTCCTGGGCGGGCCCTGGTCCGCCCACACTCGAGTAACATACTAGTGTAAACCTCCCGTCAAAGAAGCTTTAACACCAAATTCCTTCCAAAGGGGAAGGTTTAACACCATACTTTCCCCCAAAAGAGAAGGCCGACTGTCGCATCTAAATAATTAATTAATTCAATCCGCCCCGTGGAATAAATTCTATACAAAACTCAGGTTTTCCATCTCCAGCCTCTCTCTCTCTCTCTCCCGCGCCGATTCCGAAAGAACCTGTCCCCGGCAACGGGTGACAGGGA

>ERV_Env-Oan2

ATGGCTGGCGTGGGGGGCCATGGGAGCACAGGGAATGGCAAGTGGCCGAGCCAAAAAGCGAGTACCTAGCAAATAGGTCCAGGCTGAGAAAAACAAGTCAGCCTTGGGAGGCCCCCAGGGGTATGCAGTCACGCACGGCATCCGAGGTCAGAGATGTTCCCACAGTATCCGAGGTCAGAGATGTTTTCATCTGGTCTCCCTCTGATAAGGAAGTCGGGGTGCAGGTGTATGCCAAATAAACTGGGACGTGTTCTCTTCGCTTATTGGTGAAAGCTGGTTGCCATGCCACATGACCCGCTTGGCGGATAAAAGGGAATGCGAGGGGAGAACGGGGGCTGCCCCCACCGCTTTCGGTGCAATGTAACGGGAACTAATAAAGCTGCTTCAAGTTTGGTGCCTCTGGCTGACTCTTCCTTGGGAGAACGCGCAGGCCGCGTCCAGAGACGTCCAAAGACCTGAGATGGTAAGACCCTAGCATTGCGGGAAGCGGAACGCAACGAGTGGCACCCAAAGTGGGGCCCGAGGGAGAGGGGACCCCATAAAACAGGAAGCCAAGGAGGAGTGCCCCGTAGACTAGGGAGCTCATAGTCAGGCGGGAAGATGGGAGTGGAAAAGTCGCTGCCCCTGTTTAAGCCAGATGAGATGGAACTGTATGTACCATATTGTTATAAGATTCTGAAGGAACACAGGATGAACGTACATCTGAAGATTAGGGAATTTATCGGGAAGCTGACGATGATCTGCCCGTGGATCTCTCAGTCTGGACTGACTGGGGATTAGTGGAGGATAGCGGGGGAACAGATGACAGCGTATGAGGATCCTTCCCCGGGAGAGCTTACCGACCATGATTTTCTGATTTATGGCATCCTAGGGGCTGCGCTCAACGGGCCCGAGTGAAAGCCATGACACTTTAATGTCACCAGCCAGACAGAGGGTGCAGGGGAGATAGTACAGCGCCCCCCCACCAGCGACCCCGACACCGAGCTATCACAAGCTATACCCCGATCTGTCATCCGTAGGCGCGCCCTCCAGTGTAGTCCTTGAGACGCGAGATACAGCAGAACAATCGGGGCGGCTAAAAGTGATAGTATGGCGCAGTCAGAAAAGGAGGCAGAAATGACACGGCTTCGAAGGCGGCTAGCTGAACTAAGTATGGGTAGCAGGCTTAAGCACGATTCCCGAGAAAGAGGGGTGGGAGCAGAGGCTTGCGCAGGTCCGCCCGGAACGGCGATGCAATTGGCGCTGGAGGAAGGCAGAAGATTGGGGGAGGACACAGGGAGTTGGGACGCATACCCCGTGATACAGGATGGGGACGGGCAATGGGCTTTCCAGCCGATAGCGTGGATTAAACTAAAGGAACTAAAATCGGCCTGCTCCTCCTACGGGCCGGACTCTCCCTATGTGCAGCAATTGCTCGAAAATCTGACCTTAGAATCAGTCCTCACGCCAAATGACTGGAAATCTCTGGCTCGAGGTTGTTTAGACCCGGGTCAATCTATAATCTGGATGTCAGAATAAACAGCTGTGGTGAAGGATATGGTAAAAACAGCACGAGTTTCATAACCCCGCCGAGGACTTCTCAATATTAGCAGGAATGAGTAGGAATTAGCAGTATGAGACCACCGAAGCTCAATTATTGTACGACCCCCCGACGTACGTACTAGTGGGACAATTAGCCTTGACAGCCTGCAGCAAGGTCCCCCAGAAAGGGGACCGGCGCTTGCCCATGACACAGATCATGCAAGGATCTGGAGAACCTATACATGACTTTGTCTCCAGAATGCAACAAGCAGTATTGAGGTCTATAGGCGACAATGCCGGAGCGGAGATAGTCCTGAAACAGATGGTCAGGGAAAACGCTAACCCTGCTTGTAAAAAGGCGATGATGGGCCTCCCAAAGGATGCGCCCTTGGAAGATATTCTTAGCCGGTGCGAGGGATTGGGGGGCGAGGAATATAAAGCACAGGTGCTAGTGGGGGCCATAGTGAAGGGGTTACGTGGAGGAATGTCTGAGCAGAGGAAGTGTTTCTGCTGTGGTAAACAGGGACACTTGAAGGCACAATGCCGTGTGGGAGGATCCCTGAGCGTTGGGGCCCAGGAAAGGGGCGCCCCAACCTGTTTTCAGTGCGGGAAGCCAGGGCACTATGCGAAACAGTGTCGGCAGAAAAGGCACAGACAACAGTCGGGAAACTGGCGGGGAGGCCCCGCACGGGCCCCGCACGCCTACCCAGTGACGCAGTCAGAACCAAAGGACTCGCTACTCCAGGAATATCGAGAGCTCCAGACACAGGAACCCCGTGGGCAGCTGTCCGCTGCATGGACAATGTTAAAATAAACCCGGGGGACACTGTGGCCATCCCCATTTGGCCATTGCCACGGGACGTGCTTGTCACCGGTCTCGAGTCCCGAACATCAGGGGTGGTTCACACCACCACAATGGGGGGAGGCTCAAGCCAAGTCCCCCTAACAAATCCTTACCCATACCCTATATTCATTGAAAGATCGAGACCCATAGCAAAGGCCACACCGCTAGAGGCGGGACCCCCGGACCATAACCTTAGCCAAACCTTGGAAGACGGTGATTGTTGAAGGGATTCCTTTTCGAGGGACTCTCGACACCGGCACTGACCGCTCGGTGGTCACCCACTGCGATTGGCCAGCCGCGTGGCCAACGACAGAGGGACAGATGGGCATAAAAGGGATAGGCGGAACACAAACCGCCCACGAGGCTGACAGATCCCTAGCCTGGAAATGCGAGGGAAAACAAGGACAATTTGTGCCCATTGTGGTCCAAGACCTGGGAACAAACCTGTGGGGGAGAGATGTTATGCAGGGTATGGGACTAAAATTGACTGATCAGGCTGAAAATTTTGGATAGGGGCCATTGGGGTGTGGCGCCTGGAGACACCCGCGATAGTCTGGAAATCTTCCGAGCCTGTGTGGGTGGACCAGTGGCCCCTCACCTTAGAAAAGCTTCAGGCGCTCACGGAGATAGTAGAGGAGCAATTGAAAGCAGGGCACTTGGAGATCTCGTTTAGCCCCAATAATTCTCCCGTGTTTGTGATAAAGAAGAAAAACACTGGGAAATACCGCATGCTTATGGACCTCCGAAAGATTAATGCCCTTATACAGCCTATGGGGCCGCTTCAGGTGGGTTTGCCCTCCCCAAATATGATCCCAAAGGGGCAGTCTATCAGGATCCTTGATATTCGTGATTGCTTTTACAGTATCCCACTGCACGAGCAGGATAGACCCTGATTTGCCTTTACGGTACTGTCCATTAATTTTGCTGCACCCACTAAGAGATACCAATGGAAGGTCCTGCCCCAGCGCATGGTAAACAGCCCGACACTTTGTCAATGGTACGTCGGGAAGATACTGGAAGATATCCGATCACAGTACCCCGACGCCACCATGTTACACTATATGGATGATATCCTTTTGAGTCACCCATCTCCGGGTCAGCTTCACTCCCTAATGGTCGCAGTGATCGAGCACCTCAAGCACTACAGCTTGGTGGTGGCCCCTGAGAAAGTCCAAGACAAGGAACCATTCTTGTATTTGGGATTCTCCCTCCTGGGTGATCGAGTAACACAACAGACACCTCAGATCGAATTCGGTCGATATACCACTTTGAATGACATACAGCGCTTGGTGGGGCAAATCCAATGGCTCAGAGCTAGATTGCCCATTCCTTCGGGCTTGATGGAGCCCTTGTATGATCTCTTAAAAGGGGACCCAAACCTCCGGTCACCCAGGGAATGGACGACCACCGCCAAGGATGCGGTCCAGACAATTCTCCAGGTAGCTACCCGGGGCTCCACTGACCGGGTGGATCCGACGGCATGGTTGGAGGTAACTGTCTTTAGAGACCATGAACTGTTTGCCGCCTTCCACCAAGACAATCGGGTGCTCGAGTGGAGCTACTCTGGTCGCACCTCTAAGGTGCTGGAGCGGGAGGAGGTGATTCTTGGACGTTTCTGCCTCACCCTAATACGGCGGGTTCGAGTGCTGATGGGGAGCACTCCGATTATCTACTTGGGCATATCCCAGGGGGAGGTTGATGCCCTAGCACAAGAAGATCTCACCTGGGCGACAATCACGCAGACGGCTCATGTCGGGGAGAGGTCGACCCTCTTGATGGGTCCCCTCCTCCGGAATACCTGGCTGTTGCGCGCACCTGTGGTTCAGACTGAACCAGTAGCGGGCGACAATATTTTCACGGACGCAACCAAAGATAGGAGAGCCGCGGTGTATAACCAGACCTCAGGTGCCTTGACCGTGCTCTCTACCCTGCATGAGTCCACGCAGAGGAACGAGCTTTATGCCATCAGCTGGGCGTTGGAAAAATATTCTCAGCCTATTATTTCTGACAGTTTATATGCAGTTAGCCTTGTCAACAGGTTGGAGACGTCGATCCTCTTTGACCGCCGCTCAGAAATCGGGGAACAGCTGTGTGACTTGCAACGCAGGCTGCTCTCCCGGACAAGTCCTATATATGTTATACACATCCGGTCTCATACCGACAACCGCGGGCCGGTGTTTGCGGGTAACTGAGTGGTGGACGCCAGCCTCTATCAGGTACTAGCGGGGGGATCCTTCCCGGAGTCAGCGCAGAAGGCGGACGCCCTTTTTCACCTCCCCGCTGTTTCCCTCCGCCGCCTGTACGGTTTGACGAAAGCGGAAGCCCGGGGCATCGTACGCCATTGCACCCGGTGCCTCCCGTTCCTTACACGCCCACCCGGCCAGAGCGGAGTGAATCCGCGCGATTTGGTCCCGAATGACATCTGGCAGATGGATGTCACCCATTGGGGGCGCGACTGCGTCCATGTCTCCGTGGACACTCACTCCGGATTCCTGATGGCTACGAAGCAGCCAGGTGAGGCGGTTCGGCACGTGCAGAGCCACCTGTACCACTGGTTTGCCAGTAGCGGAGTGCCAAGGGAGATCAAGACAGACAATGGTCCGGCGTACACGTCGCAGACCATGGCCAAGTTTTTCACGACCTTTGGCATTAAACATGTCACCTGCATTGCCTATAACCCTAATGGTCAAGCCATCGTGGAGCATGCCAACCGGATGCTCAGGACCCTCTTGACCAAACAAGGGGTGGGATGGCGGGTGGGCCAGAGGGAGCTGGACACGGCAGTCTATACCCATAAGTTTTCTGAGTGTGGACCGAGTCTCTGGCCTCACCCCGGCTATGTGGCATCTGTGCGTCGCGGATAAGTCGGGGTTGCCCTCGCGCTGGACGCAAATGGATGACGTGGCTCAACAGGCTAAGTGGAAGGACGCCCAAGGCGGGTGGCAGGGACCACATCAGGTTCTTATCCGTGGTCGAGGATATGCTTGACCAGTCATATTTGGTCAGAGGAGAGGGGCCAGTCTGGGTCTCAAGCCGAAACTTCCGGGTCCTCGAGGGGCGACTGACGATGCTGACGGCAGTGACCCGGCCTCGGTGACCTCGGCCCCCTCGGAGGGCGGTGCACTCGATGCTACAACTTCTGCTGCTGCTCTCGCTGGGCAGTGGCGTCGGAGCAGTGCCAGGCCCAGCAGGAGGCCACCAAGTCACCTGGCAGGCCTGGTTGACTCCGCGTTGCTGGCACAGGAGGAGGTGAGCGGAGAATGGTGATGGGTGCTGGTACCACACCCTCCGGTGGTGCGGCTAGTCTCATGGCGGGAGGACTCGCCAGACCAGATCCTGTGGGGTAATATGTCCAAACTGACGGGCCAACCACGGGACACTCGTCAGAACCGTGGGGAGGAGGCGGTTGCATTGTGGAACGTTACCCTCCGGAGCCTGGGTTTGCCTATTTGTTTTTATGATGCATCCCGGAATGTGTCTGCATCGATATGACGGTACTGCTTGGCCCTCCGCAGCTGGCCGGTCACGCAGTCACCCTTCTCCGGGGCGAGGGGCATCCCACTCACCGGAACCTCACCATGATGGGTTTACACGGGGGACTTCGGTGTACATGGCCACGCCTTAATGTCACTGTTACCGATCACTCCGAGGGGAAATTTGGGCCAATTCGTAACTCCTCCCTTATCCCTTTTCCACCCCCTAAGGCGCTGCGCTGAGAGCTGTGTGGGGGTGTGGGTTCCACTGACCCGATGCTCTACGAGCATCTTCCGATGTGTAGTCTTGGGGGACAATGTCACCACCCTTATTACTTGGGACAATCCCATTCGTAACCACTCCTATAGTGAGGCCGCCGCCCCGGTCTTGTTTGCATCTCTGGCCGGTACCCAACATCCCGGCCTCTGGAAAGCCTTCACTACCTTGAGGACCATGAACGTGACGGTTCATTCATGTCCGTGCAAGGGGCCCTGTCAGGATTACCCGTATCGGGTGTGGCGGGCGGTCTTCCGAGAGGGCCTTAACATCACCCTGGAATCCACGCCGGAGAACTGTACTGATTACCGGTACCATTTCACGGGCCGTATCTTGATCCCATCCCCTTATGTTTTTTTGCATTGTCAGCTTGGCTCGAACCGCTCGGTCGAGGTGAGCGCCACGTGCGCCTCTGACTCCGACTATGCTCTCTTCGCCTCAGATTGTGACCTCACGAATGATTCCCCCTGTCAGGGACATGGGCAAGGGATAGTCACATTTCTGGTCCGGCGGCCCGGATACCTCCTGGTTCCGGTCCGCTCGGCACACCCGTGGTTCTCTTCCCTGTCCGAGCGCGACTGGTATCACGTGTCGCAATGGATCAAGCAACAGAGGCGGGAATTGTTTGAGGCTGTTTTGGGTATTGCCAGCCTGGCTTTCTCTGCCTTTCAGGAGTGGCAGATCCAAAACTTGTTTGATGGGATAGGGTATCTCGGCTCGCAGCTTCAGGCTTTTATGCATAGCACTTGTCAGGCCTTCGTGGTACAGCAGCAGCTTGATGTCTCGCTTCAGCAAGAGGTAGTGGGCTTGGAACGCGTCCTCGAAATGCTTGGGGATGAAGTGCGCCTCCTGTCTTTGAGACAGGAAGTTCAATGCGATTACCGTTACCGGCATGTATGTGTTCTTTCCCTCGTTTGTAACGCGACTAACTCCTCTTTCCCAGGCGACTGGCTCTCCGTTAAGCGTCACTTGGAAGGCCTCTTCTTGGCGGCGAACGTTTCTTCCGAGCTACAGGAGTTGCGGCTTCTGGTCAACAAATTGCGCGCTCAGGAGCTGAACTTCACCTTGTCCGGCCCGGCCAAGGACTCTCACCAGCTGGCTGTCTCGACAGGCGCCGGGGGGCCTGCCCCCTTGGCTGTGGCATGTGGTGACTGCCGCTGGTTTGTTCCTGCTTTGTCTTCTATGCCTCCCGTGCTTCGTTTGCCTTATTCTTTGGTTCCTGCGCCATGCAGTCCGGGACGTGGAGACCGAATTGTTCGCTCTTCACATGCGACGGCCCTGTGCGGTCCCCGCATAAGGGAAAGGGGGAGACGTGGGGGGCCACGGAAGCACGGGGAATGGCAAGCGGCCGAGCCAAAAAACGAGTACCTAGCAAATAGGTCCAGGCTGAGAAAAGCAAATCAGCCTTGGGAGGCCCCCAGGGGTATGCAGTCACGCACGGCATCCGAGGTCAGAGATGTTCCTACGGTATCCGAGGTCAGAGATGTTTTCATCTGGTCTCCCTCTGATAAGGAAGTCGGGGCGCAGGTGTATGCCAAATAAACTGGGACGTGTTCTCCTCTCTTATTGGTGAACGCTGGTTGCCATGCCACATGACCCGCTTGGCTGATAAAAAGAGAATGCGAGGGGACAACGGGGGCCGCCCCCGCCGCTTTT

>ERV-Env-Tac1

TTGTGAGGCTTGGAGAGGCTTGAGTCTCTACAACTAGACAGACCCCTATCTTGGGCAACCGGGCCAGTAACACAAGGAAGGAACATTTGCATAACCGCTCCTCAGACCCCTATCTTGGGCAACCGGGCCAGTAACACAAGGAAGGAACATTTGCATAACCGCTCCTCAGACCCCTATCTTGGGCAACCAGGCCAGTAACACAAGGAAGGAACATTTGCATAACCGCTCCTCAGACCCCTATCTTGGGCAACCAGGCCAGTAACACAAGGAAGGAACATTTGCATAACCGCTCCTCAGACCCCTATAAAGGGTGTAAAGTTGTGCTCGTGGGTGCTGCCGCTTCCCAAGCCTTACTATAAGGCGGTAGCCCAGATTTAAATCTGCAATAAAGCTGCGACCTGCCCTCTTTGGCGCTGGGCAGGTCATCTAAATTTTGTCCGGTTGTCGTGCGATCCAATTTGTTGGTGGGTGGGATCCCGGGCCCTGGAGACCTTGATCAGGTTTCCTTACAACATTTGGGGGCTCGTCCGGGATCCCCACCATTTTAGGCGGACCCCTTCGCTCCGCCATTCGGGATCGGCGCAACAGACCCGGTACCGGTGAGTCACTTTGTCAGGCCTCGCGAGGGTTTGGGAGTATAGGAACGGCAGGACGCTGCCTTTGTTCCGACTCCACTCGGATCAGGGGACGCTCTGATCTCGAGTTTGGGTCTCATGGTCGAGTACAGTTGTCCGTGAGACGTGTATGTTTTGGGCTGTTTGTTTGGTTGTTGGTCCGTCGTTTTTGTTCTTTGTGCGTCTTGCGAATTGCCGGTGGACCCCGGCTAGAGCCGAGGGAGAGGGGGCGCATGCGTAATGTTTTAATCTTCGGGCGAGTACGGGGTACAGGAAAGGGCCCTTCAAGCCCGTTGGAACATCTTCTTCGGTATTTTTGTCTCCCAGTTTTTCTGCTCGCACTTGCCGTCCTCTCCCTCTCGGCCAAGATGGGACAGGGCAGGTCAAAGGGCCCCTTGAGCCCGTTAGGGTGTTTGCTCAAACACTTCTCTGATTTCCAGCGGCGAGCTGATAACTATGGCGTGTCTGTTAATAACTTTGACTTACGCAGGTTTTGCGAGTTGGAATGGCCCACCTTTAAGGTTGGCTGGCCTGACACTGGGACCCTAGACATAGGGGTGGCGGCCGCCGTCCGCCGAGTTGTGGACGGGAACCCAGGCCACCCGGACCAAATCCCATACATTACCATTTGGATAGATATTATAGTAGACAACCCTAAGTACTTAAAGGACTGCGGGTGCCGGCCCCTCAGCGCTTCTAAGGTCCTCGTATCCAGTACTCTAGGGTCCAAAGCTGCTCGCAAGCCTCCCGTGCTCCCCACGCAACCGGAGAGCCCGAGGAGGGGGCGAGCGCCTCCTCCGCGAGCGCCCCCTCCCCCCTATAGGGAACCCTCAGCCCCGCCCGAGGAAGAGGCTTTTCCCCAACCAGATTCCACCGGACCTCCAAGCCCCCCCCACACCCGAAGTGGGACTGAATTTGGGCCGACAGGAAGGGCACCGGGGGTCTCGGGAATATATCCCCTGAGGGAAACAGGAGAAAGGGATGAAACGGGGCGGCCTGTGCGGACATATGTTCCTTTTACCACGTCAGATCTGTACAATTGGAAGAACCAGAATCCTTCCTTTTCCCAAGCCCCGGAGGAGGTAATCAACTTACTAGAGTCAGTCTTCTATACCCATCAACCTACTTGGGACGACTGCCAGCAACTCCTCCGCGTCTTGTTTACGACGGAGGAAAGGGAAAGAGTAAAGGCAGAGAGCAAAAAGGAGGTCCGAAATATTCGCGGTGAACCAAGCACTGACGCAGGGGAAGTGGAGGCCCAGTTCCCCTCTGGCAGGCCTGATTGGGACCCCAACACCCCGGGAGGGGAGGCCAATCTGAATCAATACCGCCAGATCCTCTTACGGGGGCTACGGGCGGCGGCCAGAAAGCCGACTAATCTCTCTAAGATAACCGAGGTCCGGCAGGGGCCAACGGAAAGTCCTACGGCCTACCTGGAACGACTATATCAGGCCTACCGGACCTGGACCCCCATAGACCCTGGGAGTCCTGATAATCAGGCAGCTATAGTAATTCAATTCGTGTCGCAGTCGGCCCCAGATATCCGAAAAAAAGATTCAAAAAATGGATGGGTTTCAGGGAAAGCCTCTCTCTGAGCTGGTAGCCATAGCCCAGAAGGTTTTTGACCAACGAGAGGACCCCACCAGAACAACTTATGAATTAACCCAAAAAATGGCGAGGGTCCTCCTAGCTCGAGAGGAACATTCAGAGAATAGGCGACGGGGAGGCAGGTCAGGTCAGAAGAGGCTGCCCCTGGGAAAGGACCAATGTGCCTACTGCAGGGAGAATGGGCACTGGAAACGGGACTGTCCCAAGTTAAAAGGGGGCGCAGCTCCGGTCCTGGTAGAGGAGGAGACTCAATAGGGCCGTCGGGGTCCCTCAGCCCTCCAGGAACCCAGGCTAAAGTTAAAAGTCGGGGGGCAATTGATTGATTTTCTGGTTGACACAGGGGCAACCCATTCAGTAGTGCAGAAACCCGTTGGTCCAATGACAAGGGATACGGTGACTATTGTAGGGGCCACCGGGGCCACGTGCAGGTACCCTAAATCGGAAGGTCGAATTGTTGATCTAGGGAAGGGATTGGTAACACACTCCTTCCTAGTTATTCCCGAATGCCCTGACCCCCTGTTGGGACGGGACCTCCTGCACAAGTTAAGGGCCACCATTATATTCCCCGAAGCGGGGACCCCTGAAATTAGAACTGAAGGCAAGTTACTGCTGTCCTCACCCTTGGTGGAGGAGTATCGTCTGTTCACTGAACAACCTGCACAAAACCTCGCCCTCTTAGATTTATGGAGGGAGGAGATCCCCGGAGTATGGGCAGAATCGAACCCTCCGGGACTCGCTACTACCCAGGTCCCCGTGCATGTCCAGCTTACCAGCACGGCCCTGCCGATCAGAATAAGGCAATACCCTATAAGTCTGGAGGCTAGAAGGAGCCTCAGGGGGAGTATTCGGAAATTTAAGGAGGCAGGAATATTGAAACCCGTCCACTCCCCTTGGAATACCCCCCTCCTACCCGTCCGGAAAACTGGGACCTCGGAATACCGCATGGTACAGGACCTGAGGGAGGTGAATAAGCGAGTGGAAACCATACACCCCACTGTTCCCAACCCTTATACCCTCCTCAGCCTTCTGCCACCTGACCGAACCTGGTATTCGGTCCTAGATCTTAAGGACGCATTCTTCTGTATACCTTTGACTTGTCAATCACAGCTCCTGTTTGCATTCGAATGGATAGACATGGAGGAGGGGGAGTCGGGCCAATTGACCTGGACCAGACTGCCCCAGGGATTTAAGAATTCCCCCACCTTGTTTGACGAAGCTTTGAGTAGAGATTTGCAGGGATATCGATTTGACCACCCAACAGTAACGCTCCTCCAGTACGTAGACGACCTTTTGATTGCCGCCGGGAGTCGAGATGAATGCCTCCAAGCTACCAGGGACCTGCTGGTCACTCTAGGATCAATGGGGTACCGCGTGTCAAGCAGCAAGGCCCAGCTGTGCCAGGAGGAGGTCACTTACTTGGGATTCAGGATCAAGGACGGGACCAGGACGTTGGCCCAGAGCCGGGTCCAGGCCATCCTGCAGGTCCCAGCCCCGAAGACCAAGAAGCAGGTACGAGAGTTCCTGGGCACGGTCGGCTACTGCAGGCTCTGGATCCCCAGCTTCGCGGAGTTGGCACAACCCCTATACGCCGCCATCCGAGGGGCCGATGCCCCCCTACGATGGACCAGTACCGAAGAGGAAGCCTTCCAGCGGTTGAAAACGGCCCTGCTGCAGCCACCTGCTCTGGCCCTACCCGACCTGGACAAGCCCTTCCAGCTTTTTGTAGACGAGGCAGAGGGTGTTGCCAAGGGGGTGCTCATGCAGACTCTCGGCCCCTGGAAGAGACCAGTGGCGTATCTCTCCAGGAAACTGGACCCCGTGGCCGCCGGATGGCCCCGCTGTCTGCGGGCCATTGCAGCCGCCGCCCTCCTGTCCAAGGAAGCGTCGAAGTTAACCTTCGGGCAGAGTTTGGAGATCACCTCGTCTCACAACTTGGAGGGTCTCCTGCGCACGCCCCCGGACAAATGGCTGACCAATGCTCGAGTAACCCAATATCAGGTCCTGCTCCTGGACCCACCCCGGGTGATCTTCAAGCAAACTGCGGCACTTAATCCCGCAACCCTGCTGCCAGCAACTGACGACTCCTTGCCCCTGCATCACTGCGCGGACACCCTGGATGCCCTAACCACCACCCGCCCGGATCTGACCGACCAACCCCTTGCCGACGCTGAGGCCACGCTCTTCACTGATGGGAGCAGTTACGTGAAGGAAGGCCTGAGGTATGCGGGGGCGGCCGTGGTGACAACGGACTCCATCGTCTGGGCTGAGGCACTCCCGAAAGGGACGTCGGCCCAGCGGGCTGAACTTATAGCCTTAACCAAGGCGCTGGAATGGAGCAGGGGTAAGACTGTGAACATCTACACCGACAGCCGTTATGCGTTTGCTACCCTGCACGTACATGCAATGATCTACAAGGAAAGGGGACTGCTGACTGCCGGGGGCAAGGCCATCAAAAACGCCTCTGAAATTTTAGCTCTTCTAACGGCCATCTGGCTGCCAAAGCGTGTCGCCGTCATCCACTGCAGAGGACACCAACAAGGTGAATCGTTGGAAGCATTGGGAAACCGGCTGGCTGACAAGACAGCCCGGGAGGTCGCTAAGAAGTCACCGGCAATTCAGGCCTCCCTGTGCGACTCGCCCCGTACCCCAGTTGACTGGGTCCCAGTGGACACCCCACAATATACAAAAGAGGAAGAGGCTCTCGGCCAACGGCTTGGCGGAACCACTGACTCGACCGGCTGGTGGCGACTCCCTGACGGGCGGATCCTACTCCCAAAAGCAGTAGGGAGGCGGGTAGTCGAGCAGACCCACCGTGCTTCCCATCTTGGGGAATCCAAACTGGCCGCGGTCATACGAAAGCACTACCTCATCTGTGGCATCTACGGGGCAGTAAAAGACGTGGTGCGCAGGTGCGAGGCCTGCGCTCGGGTAAATGCACAATCCGTCCCTACCAGCTCGGCCGAGAACGTCCGCGACCGAGGACTGGCCCCCGGGGAACATTGGGAAATTGACTTTACCGAGATGACTCCGGCCCGGGGCGGCTACAAGTATTTGTTGGTCCTGGTGGATACCTTCTCCGGATGGGTGGAGGCTTACCCGGCGAAGGGGGAAACGGCTCAGATTGTCGTCAAGCACCTGGCAAATGATCTAGTCCCGCGATTTGGACTGCCACTTCGTATTGGGTCTGACAATGGTCCGGCTTTTGTCGCAAAGATAACTCAGCAGCTGGCCTCCGCGCTCCGGATCACCTGGAAACTACACTGTGCGTACCGGCCCCAGAGCTCTGGGCAGGTGGAAAGGATGAATCGGACTTTGAAAGAAACTATCACCAAATTAAAGATGGAAACTGGGGGTGATTGGGTCGCGCTTCTCCCCCAGGCCCTCCTCCGGGCCCGGTGCACACCAGGGAGGGAAGGCCTGTCCCCCTTTGAGATTGTCTATGGTCTGAGGCCCCCCCTGGTGCCCCGAGTCGGCCTTGACCAGCTTGCCGAAGTCACCCATCGGTCTTTGCTTAAGTCCTTACAGGCGCTGCAGGCTACGCGGTCCCTCGCCCGGACGACCTTGGCAGATCAGCAACCTGGGGCAGAGGTCCATCGAGGAAGGGAGCCCCTCTTCCATCCTGGGGACCTTGTCTACGTAAAGAGACTCGACCCCCGGCAGCTCGCTCCCCGCTGGGACGGGCCCTTCACTGTCGTTTTGAGCACTCCCACTGCCGTGAAGGTAGCTGGTAAGACCCCGTGGATCCACCACACCAGGTTGAAGGGTGCCCCAGAGTGCAGTGGAACATGGGGGATCGATCCTGCCTCCACCCCTCTCAAGTTAAAACTCTCCAGACGTTTGTCATAACCTTGTTCGGAGTCTGCTGTCTCTCTGGGGTTCTGGGGGGCCTCGGGCCCCCCCCTAACAAAGAGGCACTGATAGCGGCCCTGTGGGGACCCCCCTGTAACTGTCGAGGGGGCGTCCAAGAAACCGTACCCACCACCTATACGCGATCAGTTACTTGTGGCGGCTCGACGGCTTACCTAGTGTACAACAGGGGAGTGGGTGGAGGATATCATCAACATTGGGTATGCACTCGCCGACCTAAGGTACTACTGTCCCCGGACGGTCGACCTGGGCCCTGCCCAACGGCATGCCAGGTCACCTCCCAGATGCACTCCACTTGCTATAGCGCGGCCCAGCAGTGTAACCACACGGACGGGAGGGTCTATTTGACTGCCGTCCTACAGAGAACTTACAGTGGCTCCTTTGGGGGAGAAATGGACCACTCCAAATACGCCCAGGCCTCCTGCACGGGGACCGTCGGTCAGCCCGTCTGTTGGCCACTCCGGGCCCCCCTCCACATCTCTGATGGTGGGGGGCCGTCCGATCGCGTCCGGGAAGCGCAGGTGTCGGAACGCATGAAGGAGGTAATCCAGATCTTACACCCCTCGATCCGGTATCACCCCCTGGCGCTACCTAGGCCCCGGAGCCCCGGTCTGGACCCTCAGACTGCGGATATCCTCGCCGCCACCCATCAAACTTTGAATGCCACTAACCCTGCGCTGGCGGCAGATTGCTGGCTCTGCATGCCCTTCGGCCATCCTATACCCATTGCAGTGCCCGGGAGCGCGTCCAAAAACGCTTCCGTGTCCGGGCCCGATTCTCCTTACCCCGGAGGAAACTGTACTCACAACCTTCCCTTTCGTGTGCAGCCGCTTTGGGCCCCCCCTGTTCCCTGTTACCTTAGGTCAGGCCCCAACACCAGTATCGCCTTGGACGTAGGCTTCGCCTCTTGTAACTGCTCTCAAACCGTCAATGTCTCCGCCCCACTGTGCCCGGCCCCCGGTCGAGTCTTTGTGTGCGGTGGGAACTTGGCTTTCACGGCCCTTCCCGCCAACTGGACGGGTCTTTGTGTCCAAGCCTCCGTACTTCCTGACATCGACCTTATTCCAGGTGACGAGCCTATTCCACTCCCCAGCCTGGATTATATCGCCGGTAGACATAAGAGGGCCATTCAGTTTCTCCCCCTACTCCTGGGCCTTGGGTTGGCCGGTGCGGGCATGGGAGCAACGGGTCTAGGGGTGTCGGTCCACTCCTACCATAAATTGTCCACCCAGCTCATTGAGGATGTCCAGGCTCTTTCAGGCACCATCCGCGATCTACAGGACCAGATTGACTCCCTTGCTACAGTGGTCCTACAAAACCGGAGGGGCCTGGACCTGCTGACAGCTGAACAGGGCGGGATCTGCCTAGCCCTAAAGGAACATTGCTGCTTCTACGCTAACAAATCCGGGATCGTTCGGGACAAGATCCGCAAGCTCCAGGAGGACTTGGCCGTGCGGCGGCAGGAGCTGGCCAACAACCCCCTCTGGAGCGGCTTCAATGGACTCCTCCCTTATCTGCTGCCACTTCTGGGCCCCTTGTTTGCATTGATCCTTGTGTTGTCTATCGGCCCCTGCCTGTTCAGAACTGGAGCACGTATGCTCCAGGATAGGCTGCAAGCTATTAAAGTCCTGGCCCTGATGTCCCCGTATCAACCAGTGCCCCCTGAGGACCCCTCCCCCGCGTAACCTTGCGTTTGGCTTCTGTACCCACGCTTTGCTGAGCGGTCAAAGATTTGCCCTTCACTGACAAAAAGCAGTGGGGAATGTGAGGCTTGGAGAGGCTTGAGTCTCTACAACTAGACAGACCCCTATCTTGGGCAACCGGGCCAGTAACACAAGGAAGGAACATTTGCATAACCGCTCCTCAGACCCCTATCTTGGGCAACCGGGCCAGTAACACAAGGAAGGAACATTTGCATAACCGCTCCTCAGACCCCTATCTTGGGCAACCAGGCCAGTAACACAAGGAAGGAACATTTGCATAACCGCTCCTCAGACCCCTATCTTGGGCAACCAGGCCAGTAACACAAGGAAGGAACATTTGCATAACCGCTCCTCAGACCCCTATAAAGGGTGTAAAGTTGTGCTCGTGGGTGCTGCCGCTTCCCAAGCCTTACTATAAGGCGGTAGCCCAGATTTAAATCTGCAATAAAGCTGCGACCTGCCCTCTTTGGCGCTGGGCACGTCATCTAAATTTTGTCCGGTTGTCGTGCGATCCAATTTGTTGGTGGGTGGGATCCCGGGCCCTGGAGACCTTGATCAGGTTTCCTTACAACA

>ERV-Env-Tac2

TGTTGGGGACCACCTGTCATGTAAGGCCCCTGGCCTACCCCTGTAAAGGCACCGCACCCAGCCTGCCAGTTCCAGGAAGGGCCTAACCACGAGATGTCCTCATCAGCAGATGTTCCAACGGCGATAAGCTCCGGATAGCCCCTGAAGCAAACTTCCTGATTGCCGCACCTCCCCCCACGCTCTATATATACATGTAGTGTGCAAAAATAAAGTTGACTCTTGCCTCGCACCCACCTCGGTCTCCCTTCTCTTCTTCACCCGTCCCCTTTCAGCCGCTGGCGGCTTACTCAGGCAGGGTCCTCCTCGGCCCCCACTCGACCCGGCCCCGGTGGCACGGGCAAGTGGCGCCCAACGTGGGGCCCGAGGCACGGGACCCTGCCAGGCGGACCCCTGCATTCGTTATCTGAATCGACTCGTCGCGACCCCACATTTGACCGCTCTACGGAGCCCCTGCGTTTAGGTATGTTGAGCTTCGTCCGCCTCTGTATTCTCATGGGTTCCACGCTTTCTAAGGAGCAGGCCTTTATACTTGACCTCAAACAAGCTCTTAAGGAAAGGGGGGTCAAGGTTAAGAAGAAGGACCTTATAAATTTCTTTCTCTTCATTGATGAGGTTTGCCCTTGGTTCCTTGTAAGCGGACCAGAAATTCATCCAGGCAAGTGGCAGAAGGTTGGGAGGGATCTCAACAAAAAATTACAGACTGAGGGCCCGGAGGCAGTGCCCACCACGGCCTTCTCTTATTGGAGCCTCATTAGGGACATAGTGGAGGCCGCCTCTGGGGACCCGGACAAACGCCAGCTCCTTTCAGTGGCCGAGGTTTGTCTACGCCCTTTATCGCGGGCGGCTTCTGTCAGGTCATTGGCGTCGGGGGACCAGGATCCCCCCAGGCCTCCGTCCGTGGTCATCGATATCCCGCCAGCCCCAGCGAAACCACTCTACAACCCCCTCCCTGTGGAAGCGCAATCTATGACCGACCCCCTGACCCTCCCCATCCTTGCAAGACACGACCTCCCACCAACAAAAAACTCCGACCCCGATACCCTGGACCCTGGGGAGGAAGCCGAACTGGAGGATGAGGCCGCCCGTTATAATAATCCCGACTGGCCCCCTCCTCGGCTTCTCCCCCCCCCCCTTACCCAGTTCAAACCTGTGCTCCCGCATTTCTGCCCCCTGTCCCCACCCCCTCAGCCCTTGCGGACGCAAGAAACCAATTGTCCACTCAGGTCACTGAACTCCGAGAGGTCCTTGAGCTGCAGAAACAGTATGTCCAGCTCTCCTCGGAACTCTCCTTGTTACAAAAGACGCTCCGCAAGTCTGTCATCCTTTCCCCTGACGCCCCTCAGGCATCAAAAACCCCTACAAAGAGGAATAGCCCTAGCTCCCGGGATAAGAAATCAAGAGCGTTGCAGCCAATGGCATTTCCCGTCGTAACACGCTCCCAACAGTCTGAGCCCCCACGGCCTCCAGCAGGTGACCCCAAAGAATCCTCCGCCAGTGAGGATGAGGAGGGGGAGGAAGAAGGAGAGGGCGGGGGAGAGAGCGGTGAGGAAGCCTCGACAGGGCAGGAACGACCGGAGTACCGAAAAATGCAGTTTAAAACGTTAAAAGACCTTAATGCCGCCGTCAAATCTTATGGACCCAATGCCCCCTTTACCCTCTCCGCCCTCGAGGCCATGTCTCGCGGGGGATATCTCCTACCGGCAGAATGGCTCCGGGTGGTGCAGGCTGTCCTCACACGAGGCCAGTTTCTAACCTGGAAGGCAGACTTCTTTGATCGCTGCCAGTCCATAGCGGCAGTCAACCTTAAATCCCCGAGCACTCCTGCTGCTAGGTGGACCTATGAGAAACTGAGCGGGCAAGGGAGATACGCAGGAGAAAACAGGCAGCGCCATTTTCCCATAGGCCTCCTAGCCCAAACCACCAATGCTGCGCTTGCAGCTTGGCGTGCCCTGCCAACCGCTGGTTCCCCTCTTGCCCCCCTTAACAAAATAATGCAGGGAGCGCAGGAGGACTTCTCAGAGTATGTCAGCAGGTTACTGGAAGCTACTGAGCGGACCCTAGGGCATGAAGCTGCCAGTGATCAGCTTGTAAGACGTCTAGCATATGAGAATGCGAACAATACCTGCCGCTCCACATTACACGGTAAATGGAGAGATAAAACCCTGGACGAAATGATACGCTTGTGCAGGGACATTGACCCCTTTGCTTCAAGGGTGTCACAGGCCGTGCATCTGGCTATTGGGGCAGCCCTCCAGACAGGGGACCCGCAGAGAAACTGCTTTCGGTGCGGCCAGCCCGGCCATTTTGCCCGCCAGTGCCCGGAATCCTCCCCTACTACTCCTACGCTGTACCCAGATTCCCCCCCTGCTCCCGCTCCGTATAGGTCCCCTCAGCCGTTGACCCTCTGTCCTAGGTGCAGGAAGGGGAAACATTGGGCAAACACTTGCCGCGCCATTACGGATGTCGATGGTCGTCCCCTACGGGGAAACGGCCGGAGGGGCCAGCCCCGGGCCCCTCAGCCAATCCCCTTCGTTCGGGCCTCGGGGGACAGCCCGCCTGCACCCCGCCCTACAGAGCCACCTCGGGAAGTGCGGGAATGGACTTGTGTGCCACCTCCACCACAATACTAACTCCCGAGGGCGGGGTTCAGGTCGTCCCTGCGGGGGTGTACGGTCCCCCACCCCCGAATTTGTACTTCCTCATTCTTGGTCGCGCCTCGGCTGCCATCGGCGGCCTCGTTATTCATCCCACTGTGGTGGACAGTAATTACACGGGGGAGATCCACTTGCCTGTCTCGGCCCCGAAGGGCCCCATTGCTATTTCTCAAGGACAGCGCCTGGCTCAGGCCCTGCCGCTCCCCATACAGACACGCCTGTCTCGGCCCCGAAGGGCCCCATTGCTATTTCTCAAGGACAGCGCCTGGCTCAGGCCCTGCCGCTCCCCCTACAGACAACCTATCCCGCCCCCGTTACTAAACGAGGGCGGTCTCTGCCGGGCTCCTCGGACATATACTGGGCCCAGAGCCCCTGTATTCGTCGCGACCCCTCCCCTTTTTCTTTCTGACCCCGCTTCGGAACTCCTGCATTTGGGTTCCCGGGTAGCGATTGGATCCTCCCGTTCCTGATATCGGTCGTCACGGGCACAGCGGTGCGGGCTCCGGGAAGAGGTCGGCCCCGCGTTACCAGAAAGGACCCGCCCGAAAACCGTTGGCGTGAGTTGTCACCCCCTGCGCTCCTGATCGTTTGTAGACCACCATCCGGAGAGGACCCGGAGAGATGGATAACCCTCGAGAATGAGCATGCACTGATAACCCCGATTTGGGGGTTGCCTCGATCATCTGAATCCAGAGACTGCGGGCCGCATGGACCCTGAAGGTTACGTTCAGTGGGGACAGTGGGTTCTGGGAGGTTGTTTCGGTTTACCTCTTGTGCTTGGGATTGTGATATTCTTGCTGTACTTGTGCCATACGAAGTGCAGACCCCTTAGACAGGACTAGGTGACAAGATTGTAAAGAGAACCAAGCAGCAGAGGTGAGGGCACAGACACCATGATTTCGTGGCGCATGGTCCCTGTGCTGACCCTCATCTTCCTTGCCATGGCAACCGTGCTGTCTCTTCTCCTGCTGCTGCTTCCTGGGACCCAGGGCGGCTTCGAGACCCCCGACCGCGCCGCGCTCAGCCGTACCCTCTTTGGCAGCCCCTGCGACTGCAAGGGCGGTATGCTCTCCGTGCGCCCCGCGTCGTATACACGTTCGGTTGATTGCACCACAAAAATGGCCTACCTTGCCTACCGCCACACCATTACAGGGACCTCCAAGCAATCGTGGGAATGTGTAACCAAGCCCCGGGTAATCCCTGCAATTGGCGACCAGCCTGGGCTCTGCCCCTCCGGCTGCGTCTACCTCCAGGCCCTACACTCCACCTGTTACGATTCTGTCCAACGGTGCACGGGCCCGCAGGATCAGCCCTTGCCCACTGCCATACAGCAAAGGGACTATGCTGGCACCTTCGGAGGTGAGTGGGGGTCTGACCCCCATTCCAGCAAATATGCTCAGGCCCCCTGCGACAGGAACAATGTCGGAAAGACCGTTTGTTGGCCCCTCCAGGCCCCTATCCACCTCTCTGATGGGGGTGGCCCTACGGATCAGGTCAGAGAATCCCGGGTAAGTGCTCGTGTGGAGGAGATAATTAAAAGTCTTTATCCCTCCTTGCACTATCACACCCTCGCCTTGCCAAAACCCCGGGGTACGGACCTCGATGTCCACACCTCGGAGATCCTAGCGGCCACGCTCCGGGCCCTTAACCATACCAACCCAGACCTGGCTGCTAGCTGCTGGCTATGCATGACCCTCGGCACACCCATGCCCCTGGCCCTTGTGTCCGAGAACGCCTCCCTCTCGGAGAATTGCACGCTCAGTCCGCCTTTTAGGGTGCAGCCCGTTAGCCTCGACTCCCTCCCTCCCCCCTGTACCCAAGCCCCTTTTCAAAATTCCAGTTTTGACATCGATGTTGGGCGCGCCCCCTTTGTCAACTGCTCGGTGACTGTCAACCTCTCCTCCTCGGCTATGCGCTGCCCCAGACCGGGCCAGGTTTTCGTGTGCGGGGGTAATCAGGCCTTCACCGCGCTACCCCAAAATTGGACAGGCCTGTGTGTTCAGGCATCGCTTCTGCCGGACATTGACATTATTTCGGGTACTGAGCCCGTCCCCCTGCCCAGCTTGGATTACATAGCTGGTAGATCCAAGAGGGCTGTCGTACTCATCCCCTTGCTCGTCGGCCTCGGAGTTACAGGGGCTTTGGCCACTGGCTCCGCCGGGTTGGGGGTGGCCCTAGACTCATATCGAAAGCTGTCCACCCAATTAATCAGTGATGTGCAGACCCTCTCTGAGACTATTCACGATCTCCAAGACCAAATCGACTCCCTTGCGGAGGTCGTCCTGCAAAACCGGAGAGGGTTGGATCTGCTCACAGCAGAACAGGGGGGGATCTGCCTAGCCCTGCAAGAAAAATGCTGCTTCTACGCCAATAAGTCCGGCATAGTTCGGGACAAGATCAGGAAATTGCAGGAAGACCTGGTCAAGCGTCGTCGCGAGCTCTTTGAGAATCCCCTTTGGAGTGGCTTGCGTGGCGTCCTTCCTTATCTCCTTCCTCTCCTTGGGCCCCTGTTCGGTTTCCTCCTCCTTCTTTCCTTTGGGCCTTGGGCTTTTAACAAACTAACCTCTTTCGTTAAGTCTCAGATCGAGTCCTCTCTCAGGACGCCTGTCGGTGTCCACTACCACCGCCTCGATTCACAGGATGACCCTGTGGATTCTCTCGAAGACGGCTTTCGACTCTCTACTCTCGCGCAGCCTGATTCTCGGTGCGCTAGGATGTGGCGCTCCTTTCAGGAGAAATGCCACCTGACAGCTGTGAGACGGCCGATGACGGGAATATTGAGGCCCCATGGCAGACCCCAGGCGCGGCTGCAGTCGCACCCCATGACGGGAATAGAGCGGGGCCGCCAGGGCCAGGCTGCTGCGGGACCTCCCCGCTTAGCCTAAGACAGGGGCGCTGCCTATACGCTGCAGTCGCACCCCATGACGGGAATAGAGCGGGGCCGCCAGAGCCTGGTCCTCGCGGAGCCACCCCGCTTAGCCTAAGACAGGGGCGCTGCCCGCAGCCACTCTGCCCATGGAAATCAGCCTCATAGCCTAAGCCAGGCTTTCTCTATATACTCTGAGAGGGGGAGATGTTGGGGACCACCTGTCATGTAAGGCCCCTGGCCTACCCCTGCAAAGGCACCGCACCCCGCCTGCCAGTTCCAGGAAGGGCCTAACCACGAGATGTCCTCATCAGCAGATGTTCCAACGGCGATAAGCTCCGGAAAGCCCCTGAAGCAAACTTCCTGATTGCCGCACCTCCCCCCACGCTCTATATATACATGTAGTGTGCAAAAATAAAGTTGACTCTTGCCTCGCACCCACCTCGGTCTCCCTTCTCTTCTTCACCCGTCCCCTTTCAGCCGCTGGCGGCTTACTCAGGCAGGGTCCTCCTCGGCCCCCACTCGACCCGGCCCCGGTGGCACGGGCA

>ERV_Env-Tac3_consensus

GTTTGTCTGATATCCATTTTGTTTTCCCATCCCTTCCTCCCATCCCTTTGTACACCCTTCTTCACGGAAGAAGAATGCCCGCAAAAACTTCCCGCCACAAAACCTGCCTAACCAAGGCCTCCCCCACCCTGACCTGCCTGACAAAAGCCATCCCTGTTGTCCACTAGCGGACTTGACATGACTGACTCCATTTTGTGCAGGCTGAGAGAACAACTCCTTTTTGTATTGTCTGCAACAGATGCCTTCAAGGCCATTAAACAAGTAACACTTATCTATGTAAGGTCCTAGGTCAGATGACATCCTGACAAGACAGTGACAACAGAAGATAACAGAATTTACACACCTATCTATGCAAGGCCATATGGCAGATAACAACAACCTTTACAAGGCAGTAAAAACAGAGAATTTACAGCATCCTGTTGGTCCTAATTGGATTCCTGAAATTCCTAAATGCTCCCTCCCATGTTGCTGTAACTGAATAAAAGACGCAGTGAGAATGGGATCGGGGCTGCTTGATCTGGGTCCCTGGCCCCTTGCAGCCGCGCGCTAATAAAATCCACTTCTAAATTTCTACCTGGGTCTCACTCGCTGATTCTCGGCACAACATTTGGAGGCCCCAGCGAGATGCACGGTGTCCCGTGTCACCGTGGAGACCCAACCTGGGAGGACCTCGGCCTCGGGGGAAGGAGACCATTCTGCTCTGACTTCTAGGGGGGCCGGCCCCTGAGACGTTCCAGGGCCCCGGAACTCAAGGCTGAGAGGTCTAAACTGTCCCCTGACGCCGACTGATCCTCATTTCGGCCTGGTGGATCCTGGGATCGGCAACGATCAACTACTTCCGGAGGTAACCTGGTTTCTGTTTTCTGGAGGGAACGAGTGCCGGACGCGGCAGTTAGCGCTTTGCGCCTCGATTCCCCGGGCACCCCAAGACGTTGGGTCTCTGCCCGATTCTGGTTCTGGTTCTGGTTCCGCGGTTGTCTGTGAATCTGTTCTATATGTGGAAGCGTTGCCTTTAGGAAATTCTGTTTTACGATCCTCTGTCTGAAACCTCCTCGCTCACAGACTGCCAGGCAGGTTTCCACCTGGATCGGGGACGAATATCTTCAGGGCAACTTCTCCAATGTCTACTGGAAGGATGTGTTCGCCTCCAGGTTGTTGTTTGTTTGTGTCTCTGTGTTGTATGAATGGGAGGTTCAGCCAGCACGCCAGAAACGCCTTTGAACTGTATGCTCAGTCACTTTAAAAAAGGATACCGGGATGGGTATGATTATGGGATTACCCTAAAGAAACAGAAGCTCATTCTATTTTGCATGAATGAGTGGCCCACCTTCGGGGTGGGGTGGCCCCCTCAGGGTAGTTTTGATAAGCAAATTGTAAACAAGGTTTGGAGAATTGTGACTGGGACCCCTGGCCACCCCGATCAGTTCCCCTATATTGACGTTTGGCTGGACCTCATTACCCACCCCCCTCCCTGGTTGAAAAACTGCTCCCTGAAAAGGGGCATCCGAGTTCTTTTTGCCCAACCTAAAAAAGGGCCCCCTTCCAGGGACACCCCCCCCCAAAAAAAGTCCTCCAGGAGTCCTAGGAGGATGATCTTCCCCCTCCTTGTAACCCGGGCGCCCTTCGGGACGTCCCGCAGTTTGAGAGCCCGCCCTGGACTCTCAGAGAGAGTCGAGTGGCATAGGCCCCATTAAAGGTACCCAGATTCGAGTATTGGGATGGACCTCCAAAATATGGACGGGGGAGGTTACTTATCGTTGTACGTGTTTCCATAAGCGCTTAGTACCGTGGTCAGTGCACGGTTAAAGTTCTCTAGATATTAGAAAGTGACGGGAGTGTCTGTTTGACTTGAGGAAACGAAGGTGGGAAAAGGAAAAATCTCAGTCAAAACCGAGTGGTGGGAAGTCCCTCCCGGAAATTCCGGAAGGCAGTCCGATGAACTTTCAAAGTATAAGAGAAAAGAGAAGAAATTAATTATAAAAATTATTATTAAATAATTACATATTACTGCCTAACTTGGGCTGGCGCCCACCTCGGAGGGAACGTCTTCTGGCCAAAGTACGGATCCTCCGAGGACTGGGTTTGCCAAGAATTGAACATCTATGTAATTATGAGGGAACCCCGGAATGATGGGGAATTTGTGTATGCCGCAGTCTGGCGGCCGTCGGTGGGTAAATTTGTGCTGAAGGAAATAGATGAGAAGAAAGATAGAGGGGGAAGAAAGAGAAAAACCGGCACACCAAAGTGATGTTGCAGACTCCAAAACAACAGGAAAAGCAAAGAAAGATTATGATATAAATCGTGCAGATGGTGGCAAAGTTGATCATGACTTGTTACGCCTGCGGCCGGCAGGGGCATATGAAAAGAGAGTGTCCTCAGCGGGAAAAAGAAAAGAGTTCTAGGTACATGAAGGAAAAGGGAAGGTGAAAAAATTGCAGTTGAAGGAAATTGTCTGCTCAGGCATCTACACCCTGCGAAGCAGGTGTCCCATAAGGGGGGAATGAGGGGTGGAGAGATTGCCTCTGAAGGAAATTGTATGTGTATGAAGGTTAGAGTAAGTGTGTGTGAAGGAGAAAGAGAAGGGACAGGAGATAATTTTAGTAATTCCTTCGTGGAGGGGGAAGACTAGGGGGATCAGGAGCCCATAGAACAAGCCCACCCTGAGCCCTTGACAAATTTGAGGGTGGGCAGAGAAAAACAAGATTAACAGTATTATATCCTTTTGATAGGCACGGGCGCTACCCGATCGTCTCTAACCAGACAGCCGATCAGGGCCAGAATCGAGAGGGAAATCATTAGGATCTCGGGGGTGCAGGGGGAGAACTTCCCGGTCCCTGTTACAGAGACCTTAGAACTAGAATACCTAGGGTCAAACTTCTCAGGAAAATTCCTGATCATACCCGAAGCCGGGGTAAACTTGTTGGGAGGGCTCTTAACCTCGGCGGGAAAAACCATTAAAAATAAGGAGATTTTGTCTTTACTAGACGCGGTCTGGATGCCGGCCCAGGTGGCCAGCATCCATCGCCCCGGGCACCAGCACGGGGACTCACCCGAGGCCATTGGCAATCAGGCAGCGGATGAGGCTGCCCGAGAGGCGGCAAAGTCCCCTCCCAGCGTTGCCCCCTTATGCCCTGTTTTTGACCCTAATGATACACCGGCCCCTCACTACTCCCCCTCGGATGATTCCTTCGCAAAACAAAAAGGGGGGACTAGAGACGGGTCAGGATGGTGGGTCCTTCCAGAGGGAAGAATTTTTGTCCCTGAGGCAGTAGGGAGAGAATGGATCACGCGGCTACACCAGATCACCCATCTGGGGGCACGAAAGATGGGCTTGTTGTTGAGAGATAGGTACTTTATCCCACACTTGGACAGCCCCTTAGCTAGCATAACCACCCGGTGTGGAACATGTGCACAGGTAAATGCGAAGCAGGGGAAGGCTGCTCCCTCAGGAGTCAGATTGCCGGGATTGCAGCCAGGAGAAAACTGGGAGGTAGACTTTACAGAGGTGAAACCCCCCGCGGCAGGCTATCGATACCTCCTGGTATTTGTTGACACTTTTTCAGGGTGGGTAGAAGCTTTCCCAGTTAAACACGAAACAGCCATGGTAGTGGTGAAAAAGATTCTAAATGAACTTCTCCCGCGATTTGGTCTCCCACTGGGACTCGGGTCTGAAAATGGTCCAGCATTCATAGCCAAAGTGTCCCAAGGCATAGCCAAAGCTTTAGGAATAGAATGGAAATTACACTGTGCCTATCAACCACAGAGTTCAGGTCAGGTAGAAAGGACCAACCGAACTCTTAAGGAATTCCTCACCAAACTGGTCCTTGAAACTCAGGAAAATTGGGTCATGCTCCTCCCACTGGCCCTACTCCGAAGCCGATGTACCCCCAATAAGTCGGGTCTCGCACCTTTTGAGATTCTGTTTGGTAGACCCCCGCCAATCCTCCCTCTAATCAGGGAGGAACTCAGGGCGGACGCTACTAATTCTTCCTTGATTAAGTTCCGGCAGGGTCTCCAGAAAACACAGGGAACACTCCTGAAATCTGTCCGAAACGCCCTGCCAGTTCCCACCTCTGCACCCGCACACGCCTTCCAACCCGGGGACTCGGTCCTGGTCAAGAAATTCACCGCCTCCGGCTTGGAGCCTAAGTGGAAGGGCCCTTACACCGTCATCCTGACCACGCCAACAGCCGTCAAGGTTGACTCCGTTCCTGTCTGGCTCCATCACAGTCGAGTGAAACCTGCTGCGGCCCCGACATGGAAGGCGGAGGCACAGGCCGACCCCCTAAAGCTAAGACTCTCCCGCATTTCCTCCTCCTCGCCTCCCTAATGACCCCCTCTCCCTCGATGTCATTCAACCCTCATGCCCCAAAACCCAAAAACACCTGGACGATGAAAAAGGGGGATCAGGTACTCTGGACATTCTATGCAGAAGAAGGTACATGGACCCATAACCAAGGTTTGAATGATGGCCGGTATTTCCGGTTGGATTTATGCTCCCTCTTCCCCACCTCGATCGGAGGCGCTTCCCCTTGTAGTAATCCGTTTTCTGAAGTAAGAATGTGCCCGGGGTTCCTTACAGATGGCTGGGATGCCAGGTGCCTGGATAGGTCGTCCCATTTCTGCCCGAAATGCGCCTGTGTGACCGCCATTGTAGTATCCTCGTTCCGCGGGGCGGCTTGCGGGGCCAAAGAGGGCCGCGTGGGGACGGATCCGCACTTGACTATCCAAAAGGACCCAACCCCTGGATCCACAACCTTGTTTCTCACCCTTCGCAACCCTGAGAGTAGGTTCTGGAATACTCCCCATTCGTGGGGCATGAGATTAGATGGCAGAAATAGGGCCTGGGCGGACCCTGGGATTATCTTTACCATTGCTAAGCAATCCCCCGTAACATCCTACTTGCCAATCGGCCCCCTTGGGGAATTAGCTCTGCCCCCAAAAATCCTCCCTAGACCTCGGGCCCCGTCCTCCACTGTCCAGTCCTTACCACGGGCCCCGTCCTCCACTGTCCAATCCCATCAGACACCCGCCGGGGCAATTGGCTCTCAAGAGCCAGGTGAGGGGGAGTCCTCCAGCCAGAAACCCCCTCATCCTTCAGTCATGGGCCTCTTACAGGCCGTTTATGGGGTGGTCAACTCCACTCGACCAGACCTGGGCCTAAGCTGTTGGCTATGCATGGATGCCCAGCCTCCATACTATGTTGGAGTGGCTATCAATAACTCTGTGTCTCCCACCTCCGATTCTGACAACTGTGAATGGGACCAGCCAAGGTTGACTCTTGGGGATGTCCAGGGCTCTGGGGTTTGCTTAATCTCGGATAACACGAACCTCCATGCCTCCCCATACTCGCCTGTCTGCTCCCTCAATGTGATGGTCCGGTCCTCCTCCGGTTCTGCTTACTTCCCCGCCCCACCGGGCACCTGGTGGGCGTGTTTGGACGGAATCACTCGATGTGTTTCAGCCCGAGTTTTCCTTGCTCACCCCGGTGGCCCTCTCTGTGTGCTAGTCTCCATCGTCCCCAGAGTGTCCTTGTTGCCTGGCGCTGATGGGTGGGACCACTTTTCCCTGCGGGAGGATTGGTCCCTCCGTCATAAGCGGGCTGCCCCGCTGTTCATCCCCATTCTAGTGGGGTTGGGTTTAGCGGGTTCTGCCGCCCTGGGCACTACCGCACTGGTGCGGGGGGAGGCTAGCTACAGAGAACTCAGCACCCAGGTGGATATTGACCTCACCCACCTTGAGCACTCCATTTCCACTCTGGAGCGACAGGTTGACTCCCTGGCGGAGATGGTCCTCCAGAACCGGAGGGGTTTGGACTTATTGTTTCTGAGACAGGGTGGCCTCTGTGCCGCCCTGGGAGAGGCCTGCTGCTTTTATGCGAATAACTCTGGAGTTGTTCAGGAGAGCCTCTCTCTGGTGAGGAAAAATTTAGCAGACAGGCAAAGGGAGCGTGAACGGGCCGAAACCTGGTACCAGAGTCTTTTCCGGACATCCCCGTGGTTAACCACGCTTGTGTCTGCCCTAGCTGGCCCCTTGTTTCTCCTCGTAGTTGCCCTGCTCGTCGGACCCTGCTTAGTGAATCGCCTCCTAGAATTTGTTAAGTCCCGCATCAACTCTGTTAAGCTGCTCCTCATTAGGGATCTCCACTATCAATCCCTACAAACTGAGCCCGTTGGCCGGTATGACGATGTCGCCACAAACGTGTCAAGGGTTTGACACTCTGTCCATAAGAAGTGGGGAATGTTGTGATCGGGTTTGTCTGATATCCATTTTGTTTTCCCATCCCTTCCTCCCATCCCTTTGTACACCCTTCTTCACGGAAGAAGAATGCCCGCAAAAACTTCCCGCCACAAAACCTGCCTAACCAAGGCCTCCCCCACCCTGACCTGCCTGACAAAAGCCATCCCTGTTGTCCACTAGCGGACTTGACATGACTGACTCCATTTTGTGCAGGCTGAGAGAACAACTCCTTTTTGTATTGTCTGCAACAGATGCCTTCAAGGCCATTAAACAAGTAACACTTATCTATGTAAGGTCCTAGGTCAGATGACATCCTGACAAGACAGTGACAACAGAAGATAACAGAATTTACACACCTATCTATGCAAGGCCATATGGCAGATAACAACAACCTTTACAAGGCAGTAAAAACAGAGAATTTACAGCATCCTGTTGGTCCTAATTGGATTCCTGAAATTCCTAAATGCTCCCTCCCATGTTGCTGTAACTGAATAAAAGACGCAGTGAGAATGGGATCGGGGCTGCTTGATCTGGGTCCCTGGCCCCTTGCAGCCGCGCGCTAATAAAATCCACTTCTAAATTTCTACCTGGGTCTCACTCGCTGATTC

>ERV_Env-Tac4.1

GCTGTGGGGCTGTGGGGTGGGAGTGTGAGGGATGGAACCCAGTGAAGTTCCCTCATGAGACCCCAGGAGGGAAGGCAGAAGCTATCTCCCCGGGCCAGGTTTAGGTAACACCCAGAGCAGAACTGACTTGTGCTATGGCTATGGACCGGAGGGAGATAAAAGGAAATGAAACGGCCACATTACAGAACTTGCTTACTGAAGGTAGAGCCAGTCGGGTGAGGGGCCAAACCCGTGACCCCTTTTCAAGAGAAAAACAGGATGATTCAGCTTAAACCGCAAGGCCACTGCCAGCCTCTCTCCCGCTCCGCTTCACTTCCTTGTACAGGACCGCGCGTTGGGTACTAGCTGCCGCGATTAGATTGTACTCTGCACACAGTAAGCGCTCAATAAATACGATTGAATGAATGAATGTACGCCCCAAGTACCCAGCCAATGGAGAAAGGGGTTAGTCGGTGGGAGGGGTATGCGTTAGGGTATGCGTTAGGGCTATAAAAATCGGCTTCGCCCAATCAACGGGCGCACTCCCCGGCTGACGCTGCTTAAGGGACGGGGGTGGTGCCCTTTCTCATGAGAAAGAATAAAAGCTCTTTCTATACCTGGCTCGGACTCTGATTTCGATCGAGAGGGAGTCAACATCCCACACGGTTTGGGGGCTCGTCCGGGATCCTCTCCAAGCGAGGAGAAGCCTCCCTTGGCCTGGACTGACTGGCAGGACGTCCCGCTGAAATTTTCAGCGGCCCTCGGCCCGGCCTGGGAGCGGCTCTCCTCCGACGAGAAGCGACCCGAGGAGAGACCGAAACCGGGAGTAGAAACAGCTAGCGGGGACGGGACGGAAACCACTCGCGGGAACGATTGGTCACGTAAGTCTGTGCACAGACCTTGGGTGCTGACCCCGGGTGACCGGGCTGGGCGATAAAGGAACAGACCCTATAGACCGCCTGGAACGGGGAGTTTCGGCAGAAAAATAGGAAGTCAGTTCCCGCCCCGGAGAGACACGGGTCCCGTTGAAGTCATAAATTCGGAAACGGGGGTGTCTGCCTCGGTAGAAGAATGGGAAAATCCCAGGAAAAGTCGAGTGGAGGAAGCCCCTAATAAAGGTACCTAGATTAGAAAATTGGGATGAACTTCCAAAGTATAGGAGAAAAGAGAAGAAATTAATTATAAATATTATTATTAAGTAATTAAATACTGCTGCGCAGCTTGGGCTGGCGCCCACCTTGGAGGAAACGTCTTCTGGCCAAAGCACGGATCCTCCGAGGACTGGGTTTGCCAAAAGTTGAACATCTATGTAATTATGAGGGTACTCGGAATGAGGGAATGTGCATATGCCGCAGTCTGGCGGCCGTCGGTGGGTCAATTTGTGGGGAAGGGAATAGATGAGAAGAAAGATAGAGGGGAGAAGAAAGAGAAAAACCAGCAAGACAAAGTGATGATGCAGATTCTAAAGCAACAGGAAGAGGAAACAAAGGTTATGATGCAAACCGTGCAGACGGCGGCCAAGTCGATTATGACTTGTTATGCCTGAGGCCAGCCGGGAAATATGAAAAGAGAGTGTCCTTAGCGGGAAAAAGAAAAGAATTCCATGTGCATGAAGGAAACGGGAAAAAGAAAAGGGAAGATGAAGTAATTGCAGTTGAAGGAAATTGCTCAGGCATCTACACCCTGCGAAGCAGGTGGCCCATAAGAGGGGAATGATGGGTGGAGAGATTGCCTCTGAAGGAAATTGTATGTGTATGAAGGTTAGAGTAAGTGTGTGTGAAGAAGAAAGAGAAGGGACACGAGGGAATTTTAGTAACTCCCTCGAGGAGGAGGAAGATTAGGGAGGTCTGGGGTTCATAAAATAAGCCCACCCAGAGCCCTTGATAAATTTGAGGGTGGGCAGAGAAAAACAGGATTATACCTTTTTGATAGACACGGGCACTACCCGATCGTCTCTAACCAAACTGCCGATCGGGGCCAGAATTGGGAGAGAAACCATTATGATCTCTGGGGTAAAGGGAGAGAACTTTCCAGTCTCTGTTACGGAAACACTAGAATTGGAGTACTTAAGGTCAAGATTCTAGGGGAAATTCTTGGTCATACCCGAGGCTGGGGTAAACGTGTTGGGAAGGGATTTATCACCCGGATGTCTATCCGGCTCGTCCCTATGGGGTCCCAGATTGTTCCCCAAAGGGTATTATTATTAAGAGAGGAAGATAAGAATTGAATTGATCCAAGGGTCTGGGCGGGACCTCGAAATTGGGGAAAATTGAATATACCCCCTTTGAAGATTAAGTTACGGGAACTGGGGACTATGGTGAGAGTGAGACAGTACCCCATCTCACTTGAAGGAAGGTAAGGCCTAAAACCCGTGAATCAAGGCTTATTAGAAGACGGGCTACTGGAACCTTGTCAGTCTCCTTATAATTCCCCCATACTTCCGGTCAAGAAAGGGGATGGTATTTATCGATTGGTCCAGGACCTCCGGGAAGTAAATAAAATAGTACTCCCATCCCACCCCGTGGTCCCGGACCCCTATACCATCCTGGGAAATATCCCAGCTGAGAGCAAATGGTTCAGTGTTATTGATTTGAAGGACGCGTTTTGGGCATGTCCACTGGGCACGGACAGTAGAGACCTAGCAAAAAGCTGGATCCAGCCCGAATTTCCAGAATACTGCAGATTCAGCTCCCCAAGACTAAGAGAGAATTGCGGAAATTCGTAGGGCTGGTGGGCTACTGTAGATTGTGGATTGATTCTTATGCAACCCTGACAAAACTGTTATACCAACACTTGTTAGAGGAAGAACTGGATATCATCCTATGGGATGAGGAGGTTAGGAGCAATTTTAACGGGTTGAAAGGGACCCTGAGGTCACCCCCGCTGCTGGCGTTGCCCTCGCTGGAAAAACCTTTCCATTTATTTGTTAACGTGGATGGCTCTGGGAGTGTTAGCCCAACAGTGGGGGGCGGGGGGGCAGCGGAGGCCGGTGGCTTTTCTTTCCAAGGTCTTAGACCCAGTGGCTTGGGGATGGCCCACTTGTGTGCAGCCCATGGCCGCCACGGCCATCATGGTAGAAGAGAACAGGAAACTGACATTTGGGGGCAGCCTTGTTATCAGTGTTACTCATCAAGTCAGATCTATTCTAAATCAGAGGGCTGGGAGATGGTTGACAGATTCAAGGATCCTCAAATACGAGGCCATTCTACTCGAAAGAGATGACCTGGTACTCTCCCATGACACTAATCAGAACCCAAGCCGCGTTCCTGGTGGGAGGGCCTGATGTGGAGCTTGAGGAAGGGGGACACAGTTGCATGGAACTGATCGATTTTCAGACAAAAACCCGAGAGGATCTACAGGAGTCTCCCATCCCTCCCTGACAGTGTCAATTTGTTTATAGATGGCTCTTCCCGAGTGGTGGAAGGAAAACGGAGGAATGGCTATGCAATTATCGATGGAGACAAAACGGCTGTGGTAGAACTGGTGAATTTCCCAATCCTTGGTCAGCTCAGACCTGCAAATTATATGCACCAAGTCGGGCCCTGAAACTCCTGGAAGGCGGAGAAGGGAACATCTACACTGATTCCAAGTATGCCTGGGGGGTTGTACATGTGTTTGGAAAAATTTGGGAGAAAAGGGGAATGATGAATAATCAGGGGAAAGAGTTAGCCCATACCACCCTATTACAGCAGGTACTGAAGGATTTGCATCGGCCCAAAGCGCTGGCTGTTGTACATGTAAATGGCCACCAAAAGGGGAACTCTCGAGGCTAGGGGAAACCGCCTGGCTGACCAGGAAGCAAAAAGAGCTGGAGAACTCAAACAGGAAATAACCGAACCCATGTTGGTCCTGATTCCTACATTTACCACTAATTTAAAGCCGGTATCTCTATAGGAGAAAGAGGCAAGGAGGGCCCAAGATTTGGGAGCCCAGAAGGATGAGCGGGGGAGATGGATTCTCCCATACGGGAGGGAGGTCCTTAATGAAGCCACAACGAGACAGGTGTTACAACACCTACACCAGGGTAGCCATTGGGATGTCCAGAGCCTGTGTGATGCAGTACTGGTAAAAAACATCTGTCCTGGTATCTGTACACTTGCCCGGCAAGCAGTCGATGGCTGTATTATTTGTCAAAGGACGAATAAGAACAGCCAGCGACGCTGCCCCGCGGGTGGCCAGCCTCCGGGAATTTGACCATTCCAGAGCATCCAAGTGGACTTCACTGAGGTGCCCCCGGTGGGTAGGCTGAAATACCTACTGGTGGTAGTGGATCACCTCACGTTCTGGGTGGAGGCCTTTCTCCTGGCTCAGGCCACTGCTATTGCGGTGAGTTAAAGCCCTTTTGGAACAAATTATCCCTCGATATGGATTGGTAGAGAGAATTGACTCAGATCAAGGAACCCACTTCACTGCTAGAGTCCTCCAGTCTTTGATGATAGCCCTAGAAATTTCCTGGTACTTACACACTCCCTGGCACCCTCCGTCCTCAGGCAGGGTGGAAAGAATGAATCAGGAAATTAAGAAACAGCTCACCCGATTGGTGATAGAAATCCGACTTCCCTGGACGAAATGTTTACTGTTAGCCCTCCTCCGCATTAGGACCAAACCCTGCCGGGATATAGGATTATCTCCATACAAACTCTTATACGGTCATCCCTATCCAGCTAGACTGTCTCAACCTCCCCAGTGGGAGACCAAGGACAGGTTTCCAAGGGAGTACGTGCAGTCCCTGTCAAGCTATTTGTTTTCCCTACAGAAGAAGGGGATTGTCGCTCAAACCCCTCTGCTGGGATTTCCCGTACATAAGTTCCAGGCTGGCGATTGGATCCTCATTCGGGCGTGGAAGGAGAAGAAGCTGACGCTGACCTGGGAGGGCCCATTCCAGGTACTGCTCACCACGGACACGACAGTGCGGACTAAAGAAAGAGGCTGGACTCATACCAGAGTAAAGGGACCTGTCCGAAAACCATTGGAGTGGACTGTTGCCTCCTATGATCCTGACCAATTGAGGACCACCATTCGGAGGAGACTTGGAGAGACTGATGCCCACCCCCCCCCCTTAAATGAGGAAGGACCGGTAACCCTGCCATGACTCCTGAAGATTACCTCTGGTGGGGGCAGTGGGTTCTGGAGGCTTGCTTCGGTTTGCCTTTCATGGTTGGGGGAATATTTCTGGTGTGCCTGTATATTAGGAATTGTATTTCCATTAGGCAGGACTAGGTGATAGGATTGTAAAAAGAATCAAGTAGCAGAGGTGAGAGGACAGACACACATCATGATCTCATGGTGCATGGTCCCTGTGCTAACCCTCGCTTTCCTTACCCACCTATCCATCCCTGTGCCTTTAGGAAAAAGGACCCCGCCCCAGCCAAAGGAGGAAAATACGTTTACCCAGCTAGGGGAAACGATAGCTAGCACCCTTAATGTGACCAATTGTTGGATTTGCGGTGGTCCCCAGGAGTTAGAAAGTTGGCCTTGGGTCCCTATACCCCTGGAGCCGGCCTGGATCCTCAGTAACCAGTCAGAGGTGCGTAATGGTTCTGAGTTTTGGACATCCTCAGAGAAACATCGATGGCACCTGGCGGAGTTAGTTAAGGGCCAGTATTGCCTGAATCAGTCCGGAGGAGGGGAGTCGGTGGGAGAAAGTGACTGTGTCTGGACCTATTCCTCCACAGAAAAGAAACAGGTTAACTGTACTCCTTACCAGAACTGGTGGAACGATACTCACATTAGGTGTCACAATGGGAATTGGATGGGGTGTAATGTCACCGGTACACGATCATTCTGTGCAACCAAGATGTTGACTGGTAACTCGGCCGTAGGTTGTTCCCAGTTAACTGGGGCCCAGAGCGCACTGCCAGCGTGGTGTCTTAAGTATAATGAGACTGGAGTGAACTCCAATAGCACACACGTGATCTGGGAATGGCATAATTCAACCTGGTCAGGACATTTTCCTTCTTTCTGGAGTCCATGGAATCGGAGCTCCGACCCCCTTATCAAGACTTGCGTGGAGAATGCCTCCCTAGGTCTGTGGGAGTGCTGGTTTTTAGAGGAACAGCTCGGAGGACCCCTAGGGGGAGGACCTTTTGGGGATTGGGAAGGCATCTATTACCCCGTGGCCATAAACAGCACTGAACCCTTCACCATAAATTGGACTGTAATATCCCGGAACGCGCGGCCGGCACTAAGGGGCCATTATTGGATCTGCGGCCTAACAGCATATACACACCTGCCCGCAAATTGGTCGGGCAGCTGTTATATTGGCATAATTAGACCAAAGTTCTTCTTCCTGCCAGGCCAGGAGGGGAGTCACCTAGGAATCGACCTCTATGATGATTTGAGGGATGATGGAGGGAGAGAAAAGAGGTCTCCCGATACTTCGCTAACAAGTACTGGGGACGCAAACAGGTGGGGAGACAATTGGCCTCCGGAAAGAATAATAAGGACCTACGGTCCGGCCACGTGGGCTCAGGACGGGAGTTGGGGGTACCGAACTCCAATTTATATGCTCAACCGGCTAATTAGACTCCAGGCGGTCTTAGAAATTATTACTAATCAAACGGCGAGAGCCCTTGGTCTGTTAGCAGAACAAGCCACTCAAACCAGGGAGGCTGTCTTACAGCATCGGCTGGTTTTAGACTATTTATTAGCGGCTGAAGGGGGTGTGTGTGGAAAGTTAAACCTGTCAAACTGTTGTCTAAAAATTGATGACAATGGGAGGGTAGTCATGGAAATAGCTCAGGAAATAAGGAAACTAGCCCATGTTCCTGTGCAAACCTGGAAGACTCCTTTTGGTACCTCATGTATGTCTTGGCTTGGAGGCGCCTGGTGGAGACAAATTTTATGGTTCTTGCTTCTTGCTATAAGTGGAATAATACTCCTTCCTATCTGTCTGCCCTGTATGATGCAATTAATCACTAGGATCGTACAGAGTTCAATCCAGAAGATGTTGCAAGTATCAGGGAGTGGAGAAGTTAAAATTATGATCATGAGAGCTCAAGGATGGATCCCTTTAAATACTATGGATCCATCTGATTCTGATGAGCATCCTCCGGAAGAAATGATCGAGGAGGTGTATCAGCAATGGTGTGAGGAAACTTCAGGGAGGATAAAAGAAAGGGGGATTGTGAGGGATGGAACCCAGTGAAGTTCCCTCATGAGACCCCAGGAGGGAAGGCAGAAGCTATCTCCCCAGGCCAGGTTTAGGTAACACCCAGAGCAGAACTGACTTGTGCTATGGCTATGGACCAGAGGGAGATAAAAGGAAATGAAACGGCCACATTACAGAACTTGCTTACTGAAGGTAGAGCCAGTCGGGTCGGGGGCCAAACGCGTGACCCCTTTTTGAGAGAAAAGCAGGTTGATGCAGCTTAAACCGCAAGGCCACTGCCAGCCTCTCCCCCACTCCACTTCACTTCCTTGTACACGAGCACGCATTGGGTGCTAGCTGCCGTGATTAGATTGTACGCCCC

>ERV_Env-Tac4.2

GAATGTTGGGGACCACCTGTCATGTAAGGCCCCTGGCCTACCCCTGCAAAGGCACCGCACCCCGCCTGCCAGTTCCAGGAAGGGCCTAACCACGAGATGTCCTCATCAGCAGATGTTCCAACGGCGATAAGCTCCGGAAAGCCCCTGAAGCAAACTTCCTGATTGCCGCACCTCCCCCCACGCTCTATATATACATGTAGTGTGCAAAAATAAAGTTGTCTCTTGCCTCGCACCCACCTCGGTCTCCCTTCTCTTCTTCACCCATCCCACTTCAGGCAGGGTCCTCCTCGGCCCCCACTCGACCCAGCCCCGGTGGCAGGGGCAAGTGGCGCCCAACGTGGGGCTCGAGGCACGGAACCCTGCCAGGCGGACCCCCGATACGGATCGACTCGTCGCGACCCCACTCTCGACCGCTCTAAGGAAGGAGCCCCTGCACTCGTCGCGACCCCACCTTTGACCGCTCTAAGGAAGGAGCCCCTGCACTCGTCGCGACCCCACCTTTGACCGCTCTACGGAAGGAGCCCCTCCACTCGTCGCGACCCCACCCTTGACCGCTCTACGGAGCCCCTGCGTTTAGGAACGTTGAGACACGTCCGCCTCAGTTCTCCCTCGTCGCGACCCCGCCGCTCTGAGACAGGAGCCCCTGTATTCGTCGCGACCCCTCCCCTTTTTCTTTCTGACCCCGCTTCGGAACCCCTGCGTTTGGGTTCCCGGGTAGCGATTGGATCCTCCCGTTCCTGATATCGGTCGTCACGGGCACAGCGGTGCGGACTCCGGGAAGAGGTTGGCCTCGCGTTACCAGAGGGAAAGGACCCGTCCGAAAACCGTTACCAGAGGGAAAGGACCCGTCCGAAAACCGTTGGCGTGGGTTGTCACCTCCTGCGATCCTGATCGTTTGAAGACCACCATCCGGAGAGGACCCGGAGAGATGGATAACCCTCGAGAATGAGCATGCACTGATAACCCCGATTTTCCTCCTACTGGGGGTTGCCTCGATCATCTGAATCCAGAGACTGCGGACCTCATGGCCCCTGAAGGTTACGTTCAGTGGGGACAGTGGGTTCTGGGAGGTTGTTTCGGTTTACCTCTTGTGCTTGGGATTGTGATATTCTTGCTGTACTTGTGCCATACGAAGTGCAGACCCCTTAGACAGGACTAGGTGACAAGATTGTAAAGAGAACCAAGCAGCAGAGGTGAGGGCACAGACACCATGATTTCGTGGCGCATGGTCCCTGTGCTGACCCTCATCTTCCTTACCCCCCTATCCATCCCTGCGCCGGCAGGGAAAAAGATCACACCCTGGCAGAAGGAGGAGAATACGTTTACTCAGCTAGGGGAAACGATAGCTAACACCCTTAATGTGACCAACTGTTGGATTTGTGGTGGTCCCCGGGAGTTAGAGAGTTGGCCTTGGGTTCCTATACCCCTGGAGCCGGCCTGGATCCTCAGTAACCGGTCAGAGGTGCGTAACGGTTCTCAGTTTTGGACATCCTCAGAGAAACATCGATGGGACCTGGCGGAGTTAGTTAAGGGCCAGTATTGCCTGAATCAGTCCGGAGGAGGGGAGTCAGTGGGAGAAAGTGACTGTGTCTGGACCTATTCCTCCACAGAAAAGAAACAGGTTAACTGTACTCCTTACCAGAACTGGTGGAACGATACTCACATTAGGTGTCACAATGGGAATTGGATGGGGTGTAATGTCACCGGTACACGATCATCTTGTGCAAACAAGATGTTAACTGGTAACTCAGCCGTAGGTTGCTCCCAGTTCACTGGGGCCCAGAGCGCACTGCCAGCGTGGTGTCTTAAGTATAATGAGACTGGAGTGAACTCCAATAGCACACACGTGATCTGGGAATGGCATAATTCGACCTGGTCAGGACATTTTCCTTCTTTCTGGAGTCCATGGAATCGGAGCTCCGACCCCCTTATCAAGACCTGCGTGGAGAATGCCTCCCTAGGTCTGTGGGAGTGCTGGTTTTTAGAGGAACAGCTCGAAGGACCCCTAGGGGGAGGACCTCTTGGGGATTGGGAAGGCATCTATTACCCCGTGGCCATAAACAGCACTGAACCCTTCACCATAAATTGGACTGTAATATCCCGGAACGAGCGGCCGGCGCTAAGGGGCCATTATTGGATCTGCGGCCTGACAGCATATACACACCTGCCAGCAAATTGGTCGGGCAGCTGTTATATTGGCATAATTAGACCAAAGTTTTTCTTCCTGCCAGGCCAGGAGGGGAGTCACCTAGGAATACACCTCTATGACGATTTGAGGGATAGTGGAAGGAGAGAAAAGAGGTCTCTCGATACTTCGTTAACCAGTACCGGGAATGCAAATGGGTGGGGAGACGAGTGGCCTCCGGAAAGAATAATAAGGACCTACGGTCCGGCCACCTGGGCTCAGGACGGGAGTTGGGGGTACCGAACTCCGATTTATATGCTCAACCGGCTAATTAGACTCCAGGCGGTCTTAGAAATTATTACTAACCAAACAGCGAGAGCCCTTGGTCTGTTAGCAGAGCAGGCCACTCAAACCAGGGAGGCTGTCTTACAGCATCGGCTGGTTTTAGACTATTTATTAGCGGCTGAAGGGGGTGTGTGTGGAAAATTAAACCTGTCGAACTGTTGTCTAAAAATTGATGACAATGGAAGGGTGGTCATGGAAATAGCTCAGGAAATAAGGAAACCAGCCTCATAGCCTAAGCCAGGCGTTCTCTATATACTCTGAGATGGGGAGATGTTGGGGACCACCTGTCATGTAAGGCCCCTGGCCTACCCCTGCAAAGGCACCGCACCCCGCCTGCCAGTTCCAGGAAGGGCCTAACCACGAGATGTCCTCATCAGCAGATGTTCCAACGGCTATAAGCTCCGGAAAGCCCCTGAAGCAAACTTCCTGATTGCCGCACCTCCCCCCACGCTCTATATATACATGTAGTGTGCAAAAATAAAGTTGTCTCTTGCCTCGCACCCACCTCGGTCTCCCTTCTCTTCTTCACCCATCCCACTTCAGGCAGGGTCCTCCTCGGCCCCCACTCGACCCAGCCCCGGTGGCAGGGGCAGAGGAA

>ERV_Env-Tac4.3

CTTGTGAGGGATGGAACCCAGTGAAGTTTCCTCATGAGACCCCAGGGGGGAAGGCAGAAGCTATATCCCCGGTCCAGGCTGGGTTACACCAGATACGAAGACCCGGAAGGAGATAAAGGGAAAATAAAACAGCCACATTGCAGGACCTGTTTACAAAAAAACAGGATGCAGCTAACCGAAAAACAGGATGCAGGTTAAACCGCAAGGCCACTGTCAGCCTTGTACGTGAGCGCGCGTTGGGTGCTAGCTGCCGCGATTAAATTGTATGCCCTAAGTATCCAGACAATGGAGAAAGGGGATAGTCGGTGGGAGGAAGAGGATGCGTTAGGGCTATAAAGATCAGCTTCAATCAACAATCAGCTTCAGCTACAATCAACAAGCGCACTCCGTGGCTGATGCTGTTTCAGCGACGGGGGTAGTGCCCTTTCTCATGAGAAAGAATAAAGACTCTTTCTGATCCTGACTCGGACTCTGATTTGGTGGGTAGAGGGAGCCGCTATCCCACACAGTTTGGTGGCTCGTCCAGGATCCTCTCTGAACGGGGAAAGCCTTCCTTGGCCTGGGCTGACTGGCAGGACGTCCCGCTGAAATTTTCAGTGGCCCTCGGTCTGGCCTGGGAAGAGCTCTCCTCTGACGAGAAAAGACCCGAGGAGAGACCAGAGCCGGGATCAGAAACAGCCAACGGGGGAACGACGGCGGTCACTCGCGGGAACGATTGGTCACGTAAGTCTGTGCACAGACCCAGATGGGATCTGGGTGACCGGGCAGGGTGGAGAGGTGGTCCTGTTGAGGCTGCAGAGTTGGACTCCGGCCAAGAGCACAGGAGAGAAAAGAGCCTTCTCCACCCGGGGCGGTCCTGATTGAGTCTGTTTGAGCGACGGCTCCAGGCAAGAGCACAGGAGAGATAAGAGCCCCAGCTTCCCAGAGAGTCGAGTGGCATAGGCCCCATTAAAGGTACCCAGATTCGAGTATTGGGATGAACCTCCAAAATATGGAAAAATTATACAATACATTCTATTGATATTTGCTCCCCCCCCCCCCCCCATCTAAGCTGTAAACCTGTTGTGGATGGGGGAGGTCACTGTTTATCTGTTAATCGTTGTACGTGTTTTTGTAAGCGCTTGGTACCGTGGTCTGTGCACGGTTAACGTTCTCTAGATATTAGAAAGTGTTGAGAGTGTCTATTTGACTCGAGGAAACAAAGGTGGGAAAATGGGAAAATCTCAGTCAAAGCCGAGTGGTGGGAAGCCCCTCCCGGAAATTCCAGAAGGCAGTCCGTTGAGGGCCAGGTTGGATGACTGGAATGAACTCCCCAGATATAGAGGGAAAGATAAAAGAAAAATGATTACTTATTGTTGTTTAATATGGGCGGGTGTCCAACTCGGGAAAGACGTCCTCTGGCCAAAATATGGATCCCCCGAGGACTGGGTTTGTCAAAAGCTAAATACTTATGTGAATATGAAGAAATCCCCGGAGTGAAGAGGAGTGTGCCTATGCCGCGCTCTGGATGCCATTGGCAGGTCAATTTGTGGTAAAGGGAGCAAGAGAAAAGGGGCATGGAGAGGAGAAGCCCACCCCTTGGGATCCTCTCGCTTTTCTCCCTCCCCCTTACGCCCCCCCTTCCATCGGCACAAGTGGCTGCTCCCCTTTCCCCTTCTTCTGCTTCGGCTCCAAATCCGACAGCCGAGGGAGTGAAAGGGGAAGGGGACCCGATCGATTTAAAGGAACCAAGCCAGCCGGCGCACACTCCTCTTCGCCATGAGTTAAAACGGTTACTAGAAGACCAAGAGAATTTCCCGGCTTTCGAGCAAATCTACCCACCAATACCCCCTCAGGGAGGTTCCTTTGGCTCAAGGGGGAATCGGGTATGTCAACGCTCCCCTTAGCAGCATGGAGGTTAGGAATTTTAAAAGGGAAATGAAGGTTTTAATAGAAGACCCGGAAGGGGTTGCGGAACAAATCAACCAGTTTTTCAGCCCCAACCTGTACACCTGGGCTGAGTTAATGGCAATCCTAGACATACTCTTCACTGGGGAGGAACAGGGACAGATAAGAAGGGTGGCCCTGAGAGTCTGGGATGACGCACATCCCCTTGCCCAGGGAGGACTGAGGGGGGAACAGAAATACCCCCTCCAGGACCCCCGTTGGGACCATAACGATGCCGCTCACCGGGACCACATGAGGGATCTGCGGGGGATAATAATTCAAGGAATAAGGAACGCAGTCCCCAAAAATCAGAACATGAATAAGGTTACCGGGATCAGGCAGGATAGAGAAGAGACACCATCGACCTTCCTCAATAGGCTGAAAGAGCATATGAGGAAATATTCTGGGCTCGATCTATCTGATCCCACTGCCCAGAGCCTCCTGCGGGTGTACTTTGTCATGAACTCCTGGCCCGATATTAACAAGAAACTGCAGAAAATAGAGGGGTGGCAGGCGAGGCCACTGGATGAGCTATTAGGAGCTGCCCAAAAAGTTTACGTGAGTCGTGGGGAAGAGGAGAAGAAAGAAAAAGACAGGCCAGCCAAAGTGATGTTGCAGACTCTAAAGCAACAGGAGGAGCAAACAAGGGTTATGGTACAGACCGTACAGACGGTGGCAGAAGCAGTTAGGGGACCCTATGGAAGAGGGAGAGGGCGCAGGCACCCTGGGATGGGAGAGGCTCCAAGGAAAGGAGAATTCCCTCCTCGTGCCCCACTAACTTGTTTTGGCTGTGGCCAGCAGGGGCATATGAAAAGGGACTGTCCCCAGCAGGAAAAAGAACAAAGAATTTACTCCCTTGTGGAGAAGGAAGATTAGGGAGGTCAGGGGTTCATAGAACAAGCCCACCCTGAGCCCTTGATAATTTGAGGGTGGGCAGAGAAAAACAAGATTATACCTTTTTGATAGACACGGGCGCTACCCGATCGTCTCTAACCAAACTGCCAATCGGGGCCAGAATCGGGAGGGAGACCATTATGATCTCAGGGGTGAAGGGGGAGAACTTCCCGGTCCCTGTTACGGAGACCTTAGAACTGGAATACCTAGGGTCAAACTTCTCGGGAAAGTTCCTAATCATACCCGAAGCCCGGGTAAACTTGTTGGGAAGGGATTTGATCATCTGGATGTCTATCCGGCTTGTCCCAATGGGGTCCCAGATTGTTCCCCAAAGGGTAGTATTATTACAAGAGGAAGATAAGAATCGAATTGATCCAAGGGTCTGGGCGGGACCTCAAAATCGGGGAAAATTGAATATCCCCCCCTTGAAGATTAAGCTACGGGAACCGGGGACTCTGGTGAGAGTAAAACAATACCCCATCTCACTTGAAGGAAGGCAGGGCCTAAAACCTGTGATCCAAGGCTTATTAGAAGACAAGCTACTGGAACCTTGTCAGTCTCCTTATAATTCCCCCATACTCCCGGTCAAGAAAGGGGATGGTACTTATCGATTAGTCCAGGACCTCCGGGAAGTTAATAAAATAGTACTCCCATCCCACCCTGTGGTCCCGGACCCCTAGACCATCCTGGGAAAGATCCCAGTGGAGAGTAAATGGTTTAGTGCTATCAATCAATCAATCAATCAATCAATCGTATTTATTGAGCGCTTACTATGTGCAGAGCACTGTACTAAGCGCTTGGGAAGTACAAATTGGCAACATATAGAGACAGTCCCTACCCAACAGTGGGCTCACAGTCTAAAAGGGGGAGACAGAAAACAAAACCAAACATACTAACAAAATAAAATAAATAAGATAGATATGTACAAGTAAAATAAATAAACAAATAAATAGAGTAATAAATATGTACAAATATATATACATATATACAGGTGCTGTGGGGAAGGGAAGGAGGTAAGATGGGGGGATGGAGAGGGAGACGAAGGGGAGAGGAAGGAAGGGGCTCAGTCTGGGAAGGCCTCCTGGAGGAGGTGAGCTCTCAGCAGGGCCCTGAAGGGAGGAAGAGAGCTAGCTTGGCGGATGGGCAGAGGGAGAGCATTCCAGGCCCGGGGGATGACGTGGGCCGGGGGTCGATGGCGGGACTGGCGAGAACGAGGTACGGTGAGGAGATTAGCGGCAGAGGAGCGGAGGGTGTGGGCTGGGCTGCAGAAGGAGAGAAGGGAGGTGAGGTAGGAGGGGGCGAGGTGATGGAGAGCCTTGAAGCCCAGGGTGAGGGGTTTCTGCCTGATGCGCAGATTGATTGGTAGCCACTGGAGATTTTTGAGGAGGGGAGTAATATGCCCAGAGCGTTTCTGGACAAAGATAATCCGGGCAGCAGCATGAAGTATGGATTGAAGTGGAGAGAGACACGAGGATGGGAGATCAGAGAGAAGGCTGGTTCAGTAGTCCAGACGGGATAGGATGAGAGCTTGAATGAGAAGGGTAGCGGTATGGATGGAGAGGAAAGGGCGGATCTTGGCAATGTTGCGGAGCTGAGACCGGCAGGTTTTGGTGACGGCTTGGATGTGAGGGGTGAGTGAGAGAGCGGAGTCGAGGATGACACCAAGGTTGCGGGCTTGTGAGACGGGAAGGATGGTAGTGCCGTCAACAGTCATGGGAAAGTCAGGGAGAGGGCAAGGTTTGGGAGGAAAGACAAGGAGTTCAGTCTTCGACATGTTGAGCTTAAGGTGGCGGGCAGACATCCAGATGGAGATGTCCTGAAGGCAGGAGGAGATTCGAGCCTGGAGAGAGGCGGAGAGAGCAGGGGCAGAGATGTAGATCTGGGTGTCATCAGCGTAGAGATGATAGTTGAAGCCATGGGAGCGAATGAGGTCACCAAGGGAGTGCGTGTAGATTGAGAACAGAAGGGGACCAAGCACTGAACCTTGGGGAACCCCCACAGTAAGAGGATGGGAGGGGGAGGAGGAGCCTGCAAAAGAGACTGAGAAAGAACGACCAGAGAGATAAGAGGAGAACCAGGAGAGGACAGAGTCTGTGAAGCCAAGGTCAGATAGTGTGTTGAGGAGAAGGGGGGGGTCCACAGTGTCAAAGGCTGCTAAGAGGTCGAGGAGGATTAGGACAGAGTATGAGCCGTTGGATTTGGCAAGCAGGAGGTCGATTTATCGATTTGAAGGACGCGTTTTGGGCATGTCCACTGGACCCGGACAGCAGAGACCTATTTGCCTTTAAATGGGAAGATCCGGACTTGAGTAGGAAACAACAACTAAGGTGGTCTGTCCTTCCCCAAGGATATACTGAGTCGCCTAACCTTTTCGGCCAGGTATTGGAGACCGTCCTCTAGGGTTTCCAGTCCTCCCCTCAGACTCTGGTTCTGCAATATGTTGATGACTTATTGATAGCCAGACCGAAGAGGGAGGATGTGAGCAAAACTTCTACCGAGCTTCTAAATTTCTTGGGGGACCGGGGACTGAGGGTCTCCCAGAAAAAGATGCAACTTGTGGTAAAAGAGGTGAAATATTTAGGCCACCTCATTAGCGAAGGTAGCAAGAAACTAGATCCAGCCCGAATTTCCAGAATACTCCAGATTCAGCTCCCCAAGACTAAGAGAGAATTACGGAAATTCCTGGGGCTGGTGGGCTACTATAGGTTGTGGATTGATTCCTATGCAACCCTGACAAAACCGTTATACCAACACTTGTTAGAGGAAGAACCGGATATCATCCTCTGGGATGAGGACAGTAGGAGCAGTTTTAACGGACTGAAAGAAACCCTGCGGTCACCCCCGGTGCTGGCGTTACCCTTGCTGGAGAAACCTTTTCATTTATTCGTTAGTGTGGACAGAGGGGTGGCTCTGGGAGTGTTAGCCCAACAATGGTGGGGGCAGCGGAGGCCAGTGGCTTTTCTTTCCAAGATCTTAGATCCAGTGGCTTGGGGGTGGCCCACTTGTTTGCAGGCCGTGGCCGCCACGGCCATCATGGTAGAAGAGAGCAGGAAACTGACATTTGGGGGCAGCCTGGTTGTCAGTGTTCCTCATCAAGTCAGATCTATTCTAAATCAGAGGGCTGGGAGATGGTTGACAGATTCAAGGATCCTCAAATACGAGGCCATTCTACTTGGAAGAGATGACCTGATACTCTCCCATGACACTAATCAAAACCCAGCCACCTTCCTGGTGGGAGGGCCTGATGTGGAGCTTAAGGAAGAGACACATAGTTGCATGGAACTGATTGATTTTCAGACAAGAACCCGAGAGGATCTACAGGAGTCTCCCATCCCTGACAGTGTCAATTTGTTTATAGGTGGTTCTTCCCGAGTGGTGGAAGGGAAACGGAGGAATGGCTGTGCAGTTATCGATGGAGACAAAATGGCTGTGGTAGAACTGGGTAAACTTCCCAATCCTTGGTCAGCTCAGACCTGTGAATTATATGCACTAAGTCGGGCCCTGAAACTCCTGAAAGGCAGAGAAGGGGATATATATACAGACCCCAAGTATGCCTGGGGAGTCATACATGTATTCGGAAAAGTTTGGGAGGAAAGAGGCATGATCAATAGCCAGGGAAAAGAGTTAGCCCATACCACTCTATTACAACAGGTACTGAAGGATTTACATCTACCCAAAGCCCTGGCTGTAGTACATGTAAATGGCCACCAAAAGGGAAGCTCTTTCAAGGCTAGAGGGAACCGTCTGGCCGACGAGGAGGCAAGAAGGGCTGAAGAACTCGGGCAAAGAATAACCGACCCTATATTGGTCCTGATTCCTACGTTTCCTTCTAATTTGATGCCTGTATCCCTATCGGAGAAAGAGGCAAATGGGGCCCAAGATTTGGGAGCCCAGAAGGATGAGCAGGTGAAGTGGGTTCTCCCAGATGGAAGGGAGGTCCTTAATGAAGCCACAACAAGACAGGTATTACAGCACCTGCACCAAGGTAGCCATTGGGATGTCCAGAACCTGTGCGATACAGTACTCGGAAAATACATCTGTCCTGGTATCTATACACTCGCCCGGCAAGTAGGAGCTGGCTGTATTATTTGTCGGAAGACGAATAAGAACAGCCAGCGACGCTGCCCCGCTGGTGGCCGGCCTCCGGAAATTCGACCATTCCAGAGCATCCAAGTGGACTTCACCGAGGTATCCCCGGTGGGTAGGCTGAAGTACTTATTGGTGGTAGTGGATCACCTCATGTCCTGGGTGGAGGCCTTTCCCCTGGCTCAGGCCACTGCTACTGCGGTGAGTAAAGCCCTTTTGGAACAAATCATCCCACAATATGGATTGGTAGAGAGAATCGACTCAGACCAGGGAACCCATTTCACTGCTCGAGTCCTCCAGTCTTTGATGAAAGCCTTAGAAATTTCTTGGGATTTGCACACCCCTTGGCACCCTCCGTCCTCAGGCAGGGTGGCAAGAATGAATCAGGAAATTAAGAAACAACTCACCCGATTGATGACAGAAACCCAACTTCCCTGGATAAAATGTTTACCGTTAGCTCTCCTCCGCATCAGGACCAAGCCCCGCCGAGATATAGGATTATCTCCATACGAACTCTTAAACTGTCATCCCTATCCAGCTAGACTGTCCCAACCTCCCCAGTGGGAAACTAAGGACAGGTTTTTAAGGGAATATGTGCAGTCCCTGTCGAGCCATTTGTTTTCCCTACAGAAGAAGGGGATTATCGCTCAAACCCCTCCACTGGGATTTCCCATACATAAGCTTCAGGCTGGCGACTGGATCCTCATTCGGGTGTGGAAGGGGGAAAAGCTGATGCCGACCTGGGAAGGCCCGTTCCAGATATTGCTCACCACAGACACTGCGGTGCGGACTAAAGAAAGAGGTTGGACTCATCATACCAGAGTAAAAGGACCTGTCCGAAAACCATTGGAGTGGACTGTCGCCTCCTGCGATCCTGACCATTTGAAGACCACCATCCGGAGAGGACCCGGAGAGACGGATATCCCTCGAGAATGAGAACGAACTGGTACCCCCGATTTTCCTCCTAATAGGGGCTGCCTCAATCATCTGAATCCAGAGACTACGGACCTCATGACCCCTGAAGGTTACTTTCAGTGGGGACAGTGGGTTCTGGGAGCTTGTTTCGGTTTACCTTTTGTGGTTGGGGGTATAATATTCTTGCTGTGCCTGTACCTTAAGAAGTGTAGACCCATTAGGCAGAACTAGGTGACAAGATTGTAAAGAGAACCAAGCGGCAGAGATGAGGGCACAGACACCACGATTTCGTGGCGCCTGGTCCCTGTGCTGACCCTCATCTTCCTTACCCCCCTATCCACCCCTGCGCCGGCAGGGAAAGAGATCACACCCTGGCGGAAGGAGGAGAATACGTTTACCCAGCTAGGGGAAACGATAACTAACACCCTTAATGTGACCAACTGTTGGATTTGTGGTGGTCTCCGGGAGTTAGAGAGTTGGCCTTGGGTTCCTATACCCCTGGAGCCGGCCTGGATCCTCAGTAACCAGTCAGAGATGCGTAACGGTTCTAAGTTTTGGACATCCTCAGAGAAACATCAATGGGACCTGGCGGAGTTAGTTAAGGGCCAGTATTGCCTGAATCAGTCCGGAGGAGGGGAGTCAGTGGGGGAAAGTGACTGTGTCTGCACCTGTTCCTCCACAGAAAATGAACAGGTTAACTGTACTCCTTACCAGAACTAGCGGAACAATACTCACATTAGGTATCACAATGGGAATTGGATGAGGTGTAATGTCACTGGTACACGATCATTTTGTGCAATCAAGATATTGACTGGTGACTCGGCCGTAGGTTGCTCCCAGTTCACTGGGGCCCAGAGCGCACTGCCAGCATGGTGCCTTAAGTATAATGAGACTGAAGTGAGCTCCAAGAGCACACAAGTGATCTGGGAATGGCATAATTCGACCTGGTGAGGGCATTTTCCTTCTTTTTGGAGTCCATGGAATCGGAGCTCCGATCCCCTTATCAAGACTTGCGTGGAGAATGTCTCCCTAGGTCTATGGGAGTGTTGGTTTTTAGAGGAACAGCTCGGAGGACCTTTGGGGGGGGACCTCTTGGGGACTGGGAAGGTGTCTATTACCCCGTGGCTATAAACAGCACAGAACCCTTCACCATAAGTTGGACTGTAATATCCCGGAATGCGCGGCTGGCGCTAAGGATCCATTATTGGATCTGCGGCCTGACAGCATATACACACCTGCCAGCAAATTGGTCGGGCAGCTGTTATATTGGCATAATTAAACCAACGTTTTTCTTCCTGCCAGGCCAGGAGGGGAGTCACCTAGGAATACACCTCTATGACGATTTGAGGGATAGTGGAGGGAGAGAAAAGAGGTCTCTCGATACTTCATTAACAAGTACCGGGAATGCAAACGGGTGGGGAGACGAGTGGCCTCCGGAAAGAATAATAAGGATCTACGGTCCGGCCACCTGGGCTCAGGACGGGAGTTGGGGGTACCAAACTCCGATTTATATGCTCAACCAGCTAATTAGACTCCAGGCGGTCTTAGAAATTATTACTAACCAAACAGCAAGAGCCCTTGGTCTGTTAGCAGAGCAGGCCACTCAAACCAGGGAGGCTGTCTTACAGCATCGGCTGGTTTTAGACTATTTATTAGAGGCTGAAGGGGGTGTGTGTGGAAAATTAAACCTGTCGAACTGTTGTCTAAAAATTGATGACAATGGGAGGGTGTTCATGGAAATAGCTCAGGAAATAAGGAAACTAGCCCATGTTCCTGTGCAAACCTGGAAGACTCCTTCTGATACCTCATGGATGTCTTGGTTTGGAGGCGCCTGGTGGAGACAAGTCTTATGGTTCTTGCTTCTCGCTATAAGTGGAATAATACTCCTTCTTATTTGTCTGCCCTGTATGATGCAATTAATCACTAGGACAGTACGGAGCTCAATCCAAAAAATGTTGCAAGTATCGGAGAATGGAGAAATTAAAATTATGATCATGAGAGCACAAGGATGGATCCCTTTAAATACTATGGACCCATCTGATTCTGATGAATATCCTCAGGAAGAAATGATCGAGGAGGTTTATCAGCAATGGTGTGAGGAAACTTCAGGGAGGATAAAAGAAAAAGGGGGGATTGTGAGGGATGGAACCCAGTGGAGTTCCCTCATGAGACCCCAGGAGGGAAGGCAGAAGCTATCTCCCCGGGCCAGGCCGGGTTACACCAGATACGAAGAACCGGAAGGAGATAAAGGGAAAGTAAAACAGCCACACTGCAGGACCTGTTTACAAAAAAACAGGATGCAGCTAACCGAAAAACAGGATACAGGTTAAACCGCAAGGCCACTGTCAGCCTTGTACATGAGCGCGTGTTGGGTGCTAGCTGCCGCGATTAAATTGTATGCCCTAAGTATCCAGACAATGGAGAAAGGGGATAGTCGGTGGGAGGAAGAGGATGCGTTAGGGCTATAAA

>ERV_Env-Tac5_consensus

GCACGCTCACACACACACACAGGTGCCCTCTCTCCCTCCCTTTCTCTCTCTCTCTCGTTCTCTCTCTCTTTATTAACCATGTAATCGCTGATAGCAAATAAACGGCAACCAAGCTTTGGACCTCTGGCTGACTCTTCCTGGTGTGAACGCGCGGCCTGCGTCCGGAGACGTCCGAAGACCCGGAGAGGGTAAGAACCCGAGAGTTGCCCCCGAAACCCCGGGGAGCAACGACTGGCGCCCAACGTGGGGCTCGGGTACCCCCATAGGGGAAGACCAGGGAGAGTGCCCCGTAGGCCAGGTAGCTAAAGGTCAGGCGGGAAGATGGGGACAGTACGATCGCTGCCCATATATAAGCCAGAAGATAAGGAATTGTACACCCGGCACTGTTACAAGATATTGAAAAGTAAAGGGGTAAAGATTGAGCTGAAAACAATAAGGGAATTTATAGGGAAGGTAACTATGACCTCCCCGTGGATACTTGATTCCGGGATCACGGAGGAGAGATGGGACGTTATTGGGGAACAGATGACGGCCTATGAAGACTCCCACCCGGGGGAGCTAAAGGACGTGGACTTTCTTATACACGGTATCCTCCGTGCGGCCTTTCAAGGGCCAGAGAGACTCGTTCGTAAGTTTGACGCGACCTGCCAGACTGACAAGGAGGAAGCGGGGCAGGAAGGGGAGAACAGCCGACAACCTGAGGAGACTGGTCAGGCCGAGGCAGAGACACCCCCCCCACGGGCCCCGAGTTACCACCGCATCTATCCTGATCTCGGGCCTTACACCCAGAGTGAAATGCCCAACGCCGAGCGGGAAAGAGGAGGTCAGGCTACACAGACGGAGGTTGAGAATGAGATAGGGGATACGCGGGAACAGTTCAGGCAGATGGATGTGGGAAGCAAGAAACCGATCGTTAAAAGTGGGGAGGGGACGAGAGCGGTTTCAGCCCTACAGCGAGCTTTGGAAGAGGCGGTGGCGAGGGGAGAGGACGTCACGGGATGGGAGGTATTCCCCGTGATAGAAAGACCAGACGGGGGACGAGGTTTCGCCCCGATACCTTGGGCGAAACTCAAAGAGTTGAAGGCGGCGTGTGTCGCCTACGGTCCCAGCTCCCCTTATGTGAGCCAGCTCCTGGACACTATGTCCCTGGAAAGCGTTTTGACCCCGAATGATTGGAAATCCCTTGCCCGCGGGTGTTTGGATCCCGGACAGGGCCTTATATGGATGTCTGAGTTTACCACCGCTGCGAAAGAACTAATATGCAGACGGGGTTTCCCGAACCCCGCCGAGGCTTTTGCAGCTGTGACCGGGACGGGACGATTCGAGACTCCTGAGATGCAGGTCAACTACGAGCCCGAGACGTATATGGTGATTGCCAGGGTTGCCCTAACTGCCTGGCAAAAGGTTCCCGAGAAAGGGGACCATCGCTCCCCCTTAACGCAGATCAGACAGCGCCCGGACGAGGCGATTCAAGATTTCGTTTCACGCATGCAGTCCGCGGTCACCCGTATTATAGGGGATCGGGACGGCGCCGAAATTGTATTGAAGCAGATGATCAGAGAGAACGCAAATAGTGCCTGCAGGAAAGCATTGGCAGGGCTGCCCAGAGAGGCCACATTAGGAGACATCCTGCAAAGATGCGAAGGGGTTGGAGGAGAAGAGTACAAGGCGCAGATGCTTGCGGGGGCAATAATGAAAGGATTATCAGGAGTCGGGGAAAGGGGGCGACAGTGCTTTCGGTGCGGACGGATGGGACACCTGATGGCTCAATGCCGAGCTCAGGACAAAAGTGGCCCCCCAGCACAGCCGAGAAAGGGTGTGACTTGCTTTGAATGTGGGAAGCACGGGCATTATGCGAAACAATGCCGCTCGAGGCGGAGACCCCGAACGGCGTCGGGAAACGGGTGGAGGGGCCCCGCGCGGGCCCCGAATCACGCGTTCCCCGTGTCAGCCGCGGGAGAGAGCCCGAAAAGGGTTTCTCTCATAGAGGAGTACCAGAGGCTTCGGTCCCGGGAGCCACTTGGGCTGAAGTCCGATGCCTGGCCCCAGTGGAAATCCGACCCGGGGACACTGTAAACGTCCCAATACAGCCCCTCCCCTGGAGGGCCCTAGTGGTGGGGCTTCAGACAAGAGCGGCGGGGGTCGTTCATTCGTCCGAACGGGGGGAAGGGCCCGGGAAGATTCCCCTCACCAACCATCATCCGTATGGTATATACCTAGGATCGGGAATGGTCATTGCTCGCGCTACGCCCCTTGACCCCCCGCCCCCTGATCCCCCGCCCCCAGCTATTGGAATAATACAGGAAATAACCCTTGATAAACCGTGGCGGACCCTCTTAGTTGAGGGAAAGCCCCTTAAAGGGCTCCTGGACACCGGGGCCGATCGCTCCGTTATTCAGGATTGCTGCTGGGCAGCGGAGTGGCCACTAGCAAACCACTCGATGGGGGTGCAAGGCGTGGGGGGACTGCAAGCCGCAAGAGAGGCGGGCCGTTCGTTGATTTGGTCATGCCGGGGAAGGCAAGGTGCGTTCGTCCCGCTGTGCGTTCAAGGGCTCCGTATGAATCTGTGGGGTAGAGATGTGCTGCAGGGGCTGGGAGCCCACCTTATAGATGAAGCTGATCCTTTTTAGGTGGGGCCACTGGGTTCCGGGGCTTGTCNACACCCCCACTGGTATGGCTTCCACACCCACCGGTATGGGTGGACCAGTGGCCCCTGACCAAGGACAAGCTGCAGGCGCTGAGGGAACTAGTTGCGCTCCAATTTGCACAGGGGCACCTAGAGGAATCGTTCAGCCCCTGGAACGCCCCCGTATTCGTTATAAAAAAGAAAGCCGCGGGGAAGTGGCGCTTCCTCATGGATTTAAGAAAAATCAATGCGCTCATTGTGCCAATGGGACCCCTACAACCCGGATTGCCCTCCCCTAACATGATTCCAAAAAATCACCAGATCCGGGTCATAGACATTAAGGACTGCTTTTACAGCATCCCGTTACACCCTGACGACAGGGTGAAGTTTGCTTTTACTGTCCCGAGCCCGAATTTCGCGGAACCAGCACTCCGGTACCAGTGGAAGGTGCTGCCACAGGGCATGTCTTGTAGTCCCACCATATGCCAGTGGTTCGTGGGGCAGATACTCGCCCCTTTCCGCAAGGAATACCCGGGGGCGACGATCGTCCATTACATGGACGACATCCTTCTGGGTATGCCCGACCAAGGGCAGGTACAGACGCTCACCCGGCGAGTGGTGGCTGCCCTCGCAGCCCAGGGTTTATTCGTAGCACCNGAGAAGGTACAAGAATCAGCCCCGTACACGTACCTTGGGTTTGACGTCACCGAGACCCGGGTGACTCAAAGACCACCCCAAATTGACCCCCGAAAATATGTCACGTTAAACGATATGCAGGGTTTGGTAGGGAGGATTCAATGGATGCGCGCGAGAACGCCCATCCCCTCCGCCCTCATGCAACCCCTGTACGATCTCCTAAAAGGAGACCCCAATCTTAAATCGCACCGCGAATGGACGGAGTCCGCGAAAAGCGCACTCCGAGAAATCACACAGCGGTTGGCGGGTAGCCATACTTGTCGAGCTGAACCCCACCTCCCCATGGAGGTCACGATATTTCGGGAGGGTAGCCTTTTCGCGGCTATACACCAAGGAGCCGAGATTCTCGAATGGTGTTATCCTCGAAACCCCTCCCGTGTTCTCCCAAAGGAGACTGAGCTTCTCGGTCGCTTTTGCCAGAACGCCATACAGAGGGTAGTGGCGCTTTCAGCCACTTACCCTATAGTCCATGTCGGGATAGCCATGGGAGACCTTGAGGCGATAGCCAGGGACAGCTTCCTGTGGGCCATGCTCCTACAGCAGGCCACCTTTACGGAACGCTCCCCGTTGACTCTTAGCCACTTATACGAGGGATCGGACATTCTGACCCCTCGGGTGATCTCCGATACTCCGGTCGGAGGAGACAATGTCTTTACGGACGCCACCAAGGAACACCGGGCGGCAGTTTTCAATCAGACCACCGGTGCCTTGTCGGTGCTCGACACACCATACGGCTCGACTCAGCGAAATGAACTTTTCGCTATCATATGGGCAATGACTAACTACCCCCAGGCGATCAACATTATCTCGGATAGCCTGTATGCCGTTAATCTTGCCCGGCGAATCGAAACCTCCATACTTTTCGACAGGCACTCGGAGATTGGGAACATGATCTCTCAGCTTCAGGCTGCCGTAGCGGCTAGGGAATGTAAGGTATACCTCATGCACGTCCGTTCCCACACGGACGGACAAGGGCCGATCTTTGACGGCAACCGAACTGTGGATGCCAGCCTACACCCCACAGGGCCGTCACTAATGGGGCTGGATGCCGCGGCTGCGGCGCATAGGGAGTTCCATCTCCCGGCCACCTCGCTCCGCCGGCTGTATGGGGTCACTAGAGAGGAAGCCCGGTCTATTGTCCGGCGCTGTACTCGCTGTCTTCCGTTCACCCCACGGCCGGCAGGGTCGACGGGTGTGAACCCCCGGGGGCTCACGCCGAACGAGCTGTGGCAAATGGATGTCACCCATTGGGGGACCTCCACGATTCATATGACCATTGACACCTTCTCCGGATTCATGCTGGCGACACGCCAAGCAGGAGAAGCCGCAAAACATGTGCAAAACCATTTGTACCATTGCTTTGCCACTATAGGCACACCCCAGGAGATAAAGACGGATAATGGCCCCTGCTATGTCTCCAAAGCCATGTCCCTCTTTTTCTCCTCCTTTGGCATCTCCCATGTTACCGGGATACCTTACAACCCCAACGGTCAAGGTATAGTGGAAAGGGCAAACAGGACGCTGAAAACTCTCCTGCAGAAGCAGGGGGTGGGGAAGCGGGTCACGCAGTGCGCCCTAGACAAGGCAACGTACACCCATAACTTTCTATCTGTTGATCGGGAGACGGGACTCTCCCCCGCCATGCGGCAACTCTGCCACAGGTCCGCGACCGTGGAACCCCGGCGCCACCACCCGCACGCCCGCACTATGGGGCGAGCGATGTGGCGTACTGTGGAGGGCGATTGGCGCGGTCCTGATCCGGTACTAATTCGGGGTCGAGGATATGCTTGTATCTCCACAGGTGACGGCCCCGTTTGGATCTCCTCGCGACACCTCCGCCTCGTCGAGGAAGACGATGCCCAGCCGCAGGGGAAGTCCCAGCGCCCTGATTCCCCTCCTCCTCCCCCTCCTTCCCCTTCCCCTTCTTCTTCCCGGGGGCGCGGTGGGGAGGGAGGCCGGGACGCGGATTAACTGGACTGCTCTGGTAAATAGCCACCTGAAGGATGTTGAGGAGCAGAGCGGGGTTTGGAGGTGGGCCCTCCTCGCCAATGCGCCTTTGTTACGCATGGTCTCCTGGAGGGAGGACTCCCCCCAACTCTCATTTCAAGGGAACGTGACCAAGCTCACGGGCCAGCACCGGGGTCAGTCCCCAGACACTGACCAGCATGAGATCCGTATTCGGGGCGCTATCTTGCGCAGCACTGGTCCCCCCATATGCTGGTACACCGGCAATGGGGCTTTCTCGAACCAGCCCGAGGTGACCCGATCTTGCTTGCGGTTGCTAAATTTTACTTCTACACTGGAAGTCAACATACCCCAAGGTCAGGGGCACCCCACACACCGAAACCTTACCCTGCTGGGGATCCACGGCACACTTCAGTGCACCCCCGGCGGCACAAAATCCAGCGAGGGAAGCGGGGGGTATTTTGGCCCTATCCGAAATGCGAGCTCGCTCCCCGGTCCCGGGAGAATGTGCGCCGAGGGTCGGGGTGGGGTTTGGATGCCTCTAACTGACTGTTCCACGGGCATATTCCGCTGTATTCCCCATAGTGATAACGGTACCCTCCTGCTCTCCTGGGATGACCACATTGGGGACCGCACATATACGGAGGCGGCCGCCCCAGGGCTTTACGCGCTACCGTGGGGGACGCACCACCCCGGTCTCTGGAAAGCCTTTACTTCCCTGCGCTCCCCCACCGCGACCGTCAGGGTCCGCTCGTGCGCCTGCCCTGACGGGGCGACGTGCCAGGATTACCCTTTCTCGGTGTATCGTATTATTTTCCGTGAGGGTCTGAACGTTTCCCTGCACTTGCAGGGCAATTGTGCCGACCATGATTATGTCTTTCAAGGGAACGCCCTTATCCCTGCCCCTTACGCGCTTATAGCCTGCCCAAGTACTGATCCCAGCTGGAACCTTACTGCCTCCCGGTCAGCTTATGGCGACTTTTCCTTGTCCACCACGGGTTCCTCACGATGCGTCCTTATGTCTCGGGCGCGACCCATGACCACGGAGGGTGCGGCAGATTACGCTGTTTGGCTGGTCCGGCGGCCCCGCTACACCCTTATACCCGTTAAGTCCGCTGACCCTTGGTTTTCCCAGGCCTCCGACCATCGATGGCACACCCTAGCGACCCGCTTGCGCGGCTTGCGCACCCGCAGGGAATTATTTGAGGCCATGCTGGGTATCGCGAACCTCGCGTTTTCCGCCTTCCAGGAATGGCAAATCCTGAATCTGTACGATCGGACTGGTTTCCTCGCGTCTCAGCTCTCGGCCTTCATGCACTCCACCCAACTGGCGTGGGGGGTTCAGCGTCACCTGGACGCTTCCCTGCAGCAGGAGCTGATCGGCCTTGAGCACGTGATCGAGACGCTCGGCGACGAGGTTCGCCTCCTGTCCCTACGGCAGGCGGTGAAATGCGACTACCGGTATCGGCATGTCTGCGTGCTTCCGCTCCGCTGCAACGCCACCAACTCTTCGTTCCCTGCCTCCTGGGACGGAGTAAAGAAACACCTGGAGGGACTTTTCCTGGCCGCCAACGTGACCGGCGAGTTACAGGAACTGTGCCGCCTTATTGACGAGCTCAACGCCCAGTCGCTGCGCTTCCCCAAGGCTTGGGACGACCTAGGCCGCGGGGAGGACGCGGTGGAGGAACTTTGGGACTGGATTAAACGCGTGATGCCCGGGGGATTGGCCCCTTGGCTGGTCTCGGTCCTTGCCGCCACTGGGGGGGTTTTGCTTTTGCTCATGTGCCTCCCGTGCCTCCTTCGTCTCTTTTGCTCCATTGTTGCCCGGGTGGTCCGCGAGCTTAAGGTCGAGATGCTCCCATTGCGCGGACGCCCTTAGCGGTCCCCGCCATAACGGGAGGGGGACACGTGGGGAGCGTGGGAAAGATAGGGAGTATGTGCACACAAACCGTACCCAACGGCTCAAAGCCAAACAGACTCAACAACAGGAGATAAAGGGACCAGGGCAAGAAAAACAAGCTTGCACCTGCAAAGCTCGGAGGCAACCTCAAGGCCATATCCACAGCCAGCAATGGTTGGCAACGGGTGGCTGGGGGCCAGCCAGAGCAAACGCTTCGTGATGGACAGAGCTGTTGCTAGGCTAGTGGCTGGAGGGTGGCGAGTGAGTGTGCTACATGGCCTGAGAAAATCCTGCCCCCGGGCAACGGGGGGTATAAGAGACGAACAAGACAAGGGGGGGTGCAAGCACGCACGCTCACACACACACACAGGTGCCCTCTCTCCCTCCCTTTCTCTCTCTCTCTCTTTCTCTCTCTCTTTATTAACCATGTAATCGCTGATAGCAAATAAACGGCAACCAAGCTTTGGACCTCTGGCTGACTCTTCCTGGTGTGAACGCGCGGCCTGCGTCCGGAGACGTCCGAAGACCCGGAAAGGGTAAGAACCCGAGAGTTGCCCCCGAAACCCCGGGGA

>ERV_ENv-Tac6_consensus

GACCTAACATCTGGCCCATCAATTACTGGGCCTAAGTAGTAAGCCAGGCCTTCCGTGCTCACAGGAAGACCTACAAGGAACTGAGAGGAGAGCTGTGGACAGGCAAGAGTTACTTGTTATCGGCCCTACGGCCTTGCATAGACGGGTGTTGCTTGTTACTGGTTTTGAAGGTTACAGAATGGCTGTGTTAGTTATAAACACAGCAACATGGAAAAATACAGAATGTGACTGAAAGATATAAAAGTCTTGCTGCTTAACCCAATAAACAGTCTCCTTGTCCACCTTGAGACTCGCCTGGCCTGCGTTCTTCCATTGCGAGTGCCCGTCTCCCCACTGCGCTGGACGCGTGGAGGCAGACGGCAACAAATGGCGCCCGAACAGGGACAGGGACATGAAGCCTGGACCCTCAGACTCAGCACCGCAACCCTCACGAAGCAACTCATCACTGCGCAGTGACTCACTAAGGAACCGAAGAGGTGAGGGCATTTAAGTTTACTTTCATTTTCAAACACCAGGCTTACAGCAGGGAAGAAAATGGGACAAATTCAATTCCGTTCTCGGGATCATTATATCCAATTACTTAAGGATGTTGTAAAGGAGCGTGGGTTGATTATCACCACTGCCCAGATTGAACAATTTTTGGAATGCGTGGAAAAATGCTGCCCTTGGTTTCCAAAGGAAGGAAAGTGGAATCCTTTCACTGAGCCAGCTCCTTCCGCACCACCNCCTCCACCACCCCCTCCATGGGTCCCGAAACCAGCACAAACACCTCTGCAGCGAGCGATGGGGGCTGCTATGGAAGGGGGAGAGGAGATACCTATTGAATTATTGATGGGTTTCCCTGTGATGGAGCAACCTGACCCTGCTAATCCTGGTCAGTTAATACGTACCCACGAACTCTTGAATTTCAAACTATTAAAAGAAGTGAAACAGGGTGCTGCGACATATGGGCCCACCGCTCCTTATGTGCTGGCACTCATAGAAAGTATTGCTTCCTCAATCCTTTGCCCGTATGATTGGCAAACCTTAGCCAAGACTGTCCTGGATGGCGGAGACTATTTGCTTTGGAAGGCAGAATTTTTTGATTTGTGCGCGGAGCAAGCGCGTCGCAATGATCGCGCCCGACCTCCAGTTCAAATCACTTTCGAGATGCTTACAGGGGCTTACAGGGAGCTACAGACTCGTGAAAGCTTGTCCCAGCCCCTACCCAAAGGGGTACTGATTACATGGAAAACTGTTTTTACGGCGCTGCGGGCGCTGCATCCCGCTGAACGGGTGTTACAGGAGGCGCCGGCGGCGCGCGACTTGGGGGCAGGGAGTGATCAGGAGGACCCCCTGTCCTTTGAAGTTTTGNCNCCAACAGAAGAATTGGAGCCGGTGTACGCTACTGTTCCGGATCCCGAGGACTTGTATCGGGACCAGGATCATAATGACTCTGGCGATCCTTCTGACTCTGGGACTGCAGACCCGAAGGAGGGATATGATCTATACCCGCCTTTGGCCCCTCTTACAACGACGGCTGCTGTCGCACCCACAGCAAAGGACCTTTTGCAGACCCAGNGCCTGCAGGCGGTATCCCGTTCGTCGATCGTTCCCCCTCCCCCCCTTGCTCCCCCTTTGTCTGCTGCGCCGCCGTATGTATATGCTGTGCCTGCTACGGCGCAGCCTGCCTATACCGGGGCCCTTTCGGGCTGCCGGCAGAAGGCATTGCGCCGAGGGGATGTACAGCTACTGCAAGCGATGCCGGTGGNGTATCTTCCCACCGGCCGGCCGGCTCAGTATGACTCTCTCCCTTATGAGTTGATTAAGGAGTTGCGGAAAAGTGTTCGCGATTATGGGTTACAGTCGTCCTATACAATGAATCTAATGGGGTTGATTGGGTGGATGAAGACTATNAANTAGCTGTTAGTGGCCGCTACCAACCCCGAAATTGGCTGCTTTACAAGTGATAGTCGCGGAACAATTAGCGGCGGGGCATATTGAACCCTCCGATAGCCCTTGGAATTCCCCTGTCTTTGTGATAAAAAAGCGCTCGGGGGCCTGGAGGATGTTGACTGATCTCAGAGAAATCAATAAGACGATGCAGCCGATGGGAGCGCTGCAACCTGGGCTCCCGAACCCGGCTATGATCCCTAGGAATTGGCCGATATTAGTAGTAGATATTCAGGATTGTTTTTTCTCTATTCCGCTCCACCCGGACGATCGTATTCGATTTGCGTTTTCAGTCCCTGTGGCTAACAGGACTCAACCTACTACGCGATATCATTGGACGGTCCTCCCCCAGGGAATGAAAAATAGCCCCACAATGTGCCAGACTTACGTTGCGCTTCTGATAGCGCCGCTCCGCGTCAAGCATCCTGAGGCATATATTATTCACTATATGGATGACATATTATGTGCTTGCGCCACCGCAGCCCAACTACACGCCTTACGGCCTGCTCTTCTGGCGGCTCTAGAATCTGGAGGGCTGCGGGTTGCACCCGAGAAAGTCCAGACAAAGCCCCCGATCACCTATCTGGGACATATTTTGACAGATCGGACAGTGTCCCCTGTGACCCCTACCTTGGATGTTTCAAAATTAAAAACCCTTAATGATTTCCAAAAATTACTGGGGGAATTAAACTGGGTGCGCCCTTATCTTGGAATCCCCACCGATCAATTATCTCACCTTTTTGCTACCTTGCGAGGAGCCCCGGAGCTCTCTTCCCCCCGCTCCCTTTCACCACAAGCCGCCCGGGAGTTAGAGCAGGTAGTCAAAAAATTGGCTCAGGCTACGGTGGACCGTGTCACCCCAGGGATACCCTTATCCGCCGTTGTCGTTCCCACCTCTGTCATGCCTACAGGGGTTATATGTCAAGAACAACAATTGGTTGAGTGGATGCATTTGCCCACTCAGCCCCACCGCTCTGTTTCCCCATATCCGACTTTGGTAGCCCAGTTATTGGGTCGTTTAATTCGGCGCATTACCGTCCTTTCTGGGACCGGACCCTCTGCATTATACCAACCATATACTTCAGATCAACTTGCCACACTTCTTGGCAACGATCCCGATTGGCAGATCCTTTTGGCAATGCATACCGGGGCTTGGGTTACTGGTCTCCCAGATATTCCACGATTAAAGGTTTTCTCACCTTTTGCATGGGTATTCCCCTGCATCACGGTTCCGGACCCAATACCGGGGGCTACCACAGTTTTTACAGATGGAACGAAAGGCCGAGCCGCCTATTATTGTCCTCCGGACACTTCCCGCGTTATCCCCATTCTGTCCCATTCCGCTCAACACGCGGAACTTGTAGCTGTGATGTCAGTCCTTCGCGATTTCCCTCAATCACTTAATATTATCTCAGATAGCCGGTATGCATGCTTTGTTACTCTTCACATTGAGACTGCTGCGCTCCGGTTTTCCGAGGCAGCCATTTTTCCTTTGTTCCAGGAACTTCAAGCAACGATTCGCGCTCGTTCTCATCGTTTTTATATTTCTCATATTAGGGCGCATTCGTCCCTCCCCGGACCATTGGTTGAAGGTAATGCTTGTGCGGACCTACTTACTAGGGAAATTGCTGGCGCACCAAAGGCGATTAATTTTGTTGCACCACTGTTGGCTCCGGCTCTTCTCCCTTTAGATGCTACAGCCGCAGCCTCGCAAGCCCACCGGCGCTTCCACCAAGGGGCAGGAATGTTGCGCCGTCAATTTGGCATTTCTCGGGAAAGCGCGCGGGCTATTGTGAAAGCTTGTACCTCCTGTGTTACTACATTACCTGCGGCCTCTGAAGCCACGAATCCCCGTGGTCTTGCGCCAAGAAAGCTGTGGCAAATGGACGTCACGCATTACCCTCCCTTTGGCAGATTGGCTTTTATTCATGTTACTGTTGATACCTGTACATTATTTACTTTTGCGTCGGCTCACACTGGTGAATCAGCCAAACATGTTATGGATCATTTGTTTCTTGCTTTTGCCTTTATGGGAATTCCACAGGTATTGAAAACCGACAATGGCCCCGCATACACTTCTAAGGCGTTCCGGTCTTTCTGTAATACTTTTGCTATCCGCCTTGTTACGGGTATTCCATACAATCCCACGGGTCAAGCCATTGTCGAAAATCGTCATCGTTGGCTCAAAGCCTTTTTGGAAAAACAAAAAGGGGGAGATGAACTGACCTCTCCCCATAGACAATTACAATCCGCTATATATACCCTTAATTTTTTAACTATGGATGACAAAGGTCTCACCCCCGCTGAAAAATGGGGTGGTAGGGAAACTAAAGAACACCCAGAAAGAGTAAAATGGAAGGACCCCCTCACGCGACAGTGGCGAGGGCCAGATCCCTTATTAACGTGGGGCCGAGGTTATGCTTGTGTCTTTCCAGAAACTGAGGACCGCCCGATCTGGATACCTACAAAGAACATACGCCGCGTCCTTTCTACAGTGCGTTCTCCCTCACCCTGCGTACTTCAGGAGCTGACAGAGAACCAGGATACAGACTCCGCGACCCTTCCTCTGGAGGATCAGCAGCAAAGTTCCTAAAAATGATGATCGCTTTCCTGATTCTCGCCTCCTCCACTGTTGTGATTACTGAGGACTATTGGACTTTTGTTCCGTACCCTCCGTGGGTGCGGCCCGTTACGTGGCAGGACCCGGTGGGAGATATTCTCACTAATGCTTCTGATCTTGTAGGGGGGGATGCGATTCCATGGGAAGGACCAGGGGAAGAGGCACTTGTCAATGTCTCCCTCCTTGCTACTCGCACTCCTATATGTTTCTCTGCAGCTCCGACCCCAAGTCCGTGTCTGTCTCTGCTCCCCCTCACTACTTGGGCTCAAGAAAGAGAATTGCGTGAACCCCAGCCCCGTTCGGTGTACTGTGGGATTACCCGTTTGACTGCTGTATCCCATCGTTTGGGAACCTCACCTTCGGTACCCGAACTTCCTTCTTGTGTGTTTGCCAGCTCGTCTTGGCCACCATCAGAGTCTGTTTTGCCTATTTGGCAAACGTGCCGACAACCTTTTCCCACGGTTATTCATTTGTCCTCTCACGTATTCCCAGCCACCTTACTGAATTGGGGCGGACATGACAACCTTACTTATTTATCCCCTTTGCATGGTTATCTTGGGTTTCTGCCCACGGATGCTTTCCAGGCTTTAGGTTCCTCTCGTGCCGGAACTTCCTCCCCTTTGTACCCACACCTTTGGCGGGCTTTTGCTGCTTTGGGTCCAATCACACTTGCTTTTGCTCGATTACAGGTCCCTTTCGATTCTATCCCAGTCCATATGGAAGAACGGCAAATAAGAGCATGTATATTGCCACCGCAAGCCTTCCTTGTTGGCTACCCGTCCCTTATCCCCTTCCCTTCCCTCTGGAATATTACATGTGATAATTGTAGACTTACCCAATGTATATCCGCCCATGATGCCACTGCTACTATTCTTATAGTAGAACAACCTCGTCATGCTATACTACCCACCTCATTATCCCACCCTTGGGCTGAGTCCCCTGCGGATTCTGCCCTGCAGATGCTAGCTGATCATTTTCGTATTCCAAAACGGCGACATCGTCGGTTTGTTGGATTGTTAATCACTGGGTTTATCGCTTTGGCATCCCTTATTATTGCTACCACCTTATCCTCTATTACCCTGCATACTGAAATAACACAGGCCTCCTTTGTTAATTCCCTTGCAAGAAATGTTTCTGATGCTCTGCAAACACAGTTACGATTAGATAAAGCCTTTACCGCTCGCCTCTCTCTTCTACAGCATTCTCTTATAGAATTAGGAAATGAGGTTGATGCTCTGCGCACTCAACTTTCTTTTCAACGTTGTGATTATCGTTACCCTCATATCTGTGTGACCCCATATTATTATAATGCCACATTTCCCGGGTTTTCGTGGTCCGCGATCCGAAATCGGCTCCAAGGAGCCTGGCATTCTTCGAATGTCTCCCTTGATTTGTCACACCTTGCCTTACTCTCCGATGCTATGCAAAATACCCCTCCCTTGACCATTCCTGCCGATATTGCTTCACCGATCATTTCTGCCCTTGAGGCGGTTAATCCTTTTAATTGGCTCTCAATGCTGCTTCATAGTCTCCCATATATTATTTTGTTGATCTTGGCCCTGTTTTGCCTGCCCTGCCTTGCCCGATGGCTATCTGTCTTCCTTCGCCCGCTTCTTACTCAACTCCATGCTTTGCAATTGTTTCAAAAACAAAAAGGGGGACTTGTGGCAGACCTAACATCTGGCCCATCAATTACTGGGCCTAAGTAGTAAGCCAGGCCTTCCGTGCTCACAGGAAGACCTACAAGGAACTGAGAGGAGAGCTGTGGACAGGCAAGAGTTACTTGTTATCGGCCCTACGGCCTTGCATAGACGGGTGTTGCTTGTTACTGGTTTTGAAGGTTACAGAATGGCTGTGTTAGTTATAAACACAGCAACATGGAAAAATACAGAATGTGACTGAAAGATATAAAAGTCTTGCTGCTTAACCCAATAAACAGTCTCCTTGTCCACCTTGAGACTCGCCTGGCCTGCGTTCTTCCATTGCGAGTGCCCGTCTCCCCACTGCGCTGGACGCGTGGAGGCAGAC

>ERV_Env-Tac7.1

GGATCAGTGGAAAAGAGCATGGGCTTTGGAGTCAGAGGTCATGGGTTCAAATCCCTGCTCTGCCAATTGTCAGCTGTGTGACTTTGGGCAAGTCACTTAAATTCTCTGTGCCTCAGTTACCTCATCTGTAAAATGAGGGTTAAGACTGTGAGCCCCCCGTGGGACAACCCGATCACCTTGTAACCTCTCCGGCGCTTAGAACAGTGCTTTGCACATAGTAAGTGCTTAATAAATGCCATTATTATTATTATCATTATTATTATATGACTATCCCCTAAGGCTTACAACATCCAATCCTTCCATCCTCCTATGGCATCAGGGACACTGGGGCCTCTGTTGGACATTAGACAGGCATCCCCACAAAGTTGTTGAAGGTAGAGCCAGCTGAACACTCCACTGTCCACTAGTGGACAACCAAGCCAGGGCTGGTTTCTGCACGGAGCCCCAACCCAAAAGCCTTGTATAGCAAGGGGCTGGGGGATGTACCAGGCCATGGAAACTGGGATAGGGACAAGGTAAGTAAGTGAACATACTTGTCCTCAACTTCTCCCTGCCCCTTCCTCCTCAATCCTAATTCCCTTCTTGGACATAACATCTCCTCCAAGAGGCCTTCCCCAATTAAGCCCTTTTTTCCCCTGGCTCCCTCTCCATTCTCCCTTGTCTAGGCACTTGAATCTATGACCTTTGGGCAATTTGATATTCACTCCACCTCCAACTCCTCAATACCTTGTACATTTCTTTAAATTATATATTATGAATTATCTATTTATAATATTATCTGTCTGCCCCTCTTGACTGTAAGTTTGCTATGGGTGGGGAACATTAACCACTAATTCTGCTGTATTGGACTCTCCCAAGGGCTTAATACAGTGCTCTGCACATAGTAAGTGCTTAATAAATACCACTGACCTGTTTACTGATCCCAGAGTTTGAACTTTCTCCTTCCTAGTGCGCTTCTCAAGAGTTCTGCTCAAGGGACTATGTGTTTTCCTGGGGTTTGGGGAAGTGAATATATTAGTCTGTTAGAGAAAGAGGTATGAGAATTTCCGTGCCGTGACCCATCCATTCCCACATCTTCTCTGCTACTACCACCAGTCCCCATAGGAGTGTTCGAAAGTTTCAGAGGGTGCAGACTAGTTGGGGTGAGAGCAGGGTCCGAAGGTGGAGCCAGACAGGGTAGGAACACTGTCATAAAGACTTATCAATAGGAGAAGCAGCGTGGCTTTGTGGAAAGAGCACGGGTTTGGGAGTCAGAGGATGTGGATTCTAGTCTCTGCTCTGCCACTTGTCTGCTGTGTGTCCTTGGGCGAGCTGATTAACTTCTCAGTGCCTCAGTTACCTCATCTGTAAAATGGGGATTGAGACTGTGAGCCCCACATGGGACAGCCTGATTACCTTGTATCTAGTCCAGTGCTTAGAACAGTGCGTGGCACAGAGTAAGCGCTTAACAAATACCATAACAATGATAATAGTAGCAATGGTTCCTGGCTAGGAGATCTCCACCAAGGGGCAAAGAGTCGCACTAGGACTGAAGGATGGGTGGTCGATTCAACCTGCTCAACCTGTTTGTGACCAGGCTGCAGGAGCCCAGGATGCTGGGCATCTTAATTGCGTGTGAATCATTGGTTCAACGTGGTGAGGGCCAGGCCTACCTGTGATTACTGCTTCCCTGGTGCACCTCCATCTGTCCTAGGATGCATGTGGAGACCACAGCAGTCGGTGTGGGTACTTGTTTTGAAGACAGTAGGTAGAGACAGTGAAATTGAATGAAGACCTGGCTCTCCTGACAGTAGGGCTCCCTCTGCAAGACTGCAAAGGCAGCAGCCGAGAGGGCCAAGGCAGCCGTTTCGGGGAGTGGAAACTCATCGGCACCCTTTCACTGACCACAATGGTAACAATATTGCAGCGTGGTTGCCAATCAGCACCCTGCAATATCCTGAAATCCAAGAGTGGCCTGCTGACTATCTCTGTAGCTGGAAAATGCTGTTTGAATGGGGTGGAAAATCCACAGTCCCTGAAAGACTTAATGAACCATTGGAGGACTGGGAAATGTTGAAGGTGGCAGTGCCAACAGGAACTATTGGGCTGGTTGAGGTATAAATCTTAGGATCTGCTAGACATAGGTCTTCGGGGGAAAATAATTAAGGTTGAGCAGAAAGCCTGAGGTAGGGAAGGTGATGGTAGCAGGAAGAGAGGATTATCTTCAGGGTAAAACAGATTGGGAAGCAAGAACAGGGAAGGTGATTAATCAGTCAAGCAAACAATCTGTGTACAAAGTATTGTACTAAGCTTTGGGGAAAGCACAATTATAATTGTACTTTCCTGCCCATAAGGAGGAGGTTACAGTCTCCAAGAGGAGACAGATATTCAAATAAAATACAGATAGGGGACATAGTTAAGTATCAAGATATTTACGTAAGTTCCATAGGACTCTTATGAGTATCAAAGAGCTTAAAATGTACACAGAAAAGTACATAGTTGATGGTGAGAAGAGTGGAGATAAGCAATATAAAGGCTTAGTCAGGGAAGGCCTCTAGGAAGAGATGTGATTATAGGAGGATGCCTCTAGGAAGAGATGTGATTATAGGAGGATTTTGAAGGTGGAGAGAGTTGTGTTCAGTGATATATGAGTAATAACAATAAAAATTATTGTATGTGTTAAGCGCTTACTATGTGCCAGGTACTGTACTTGGGGCTGGGGTGGATACAAGCAAATCGGGTTGGACATAGTCCCTGTTCCATGAGGGGCTCAGTCTCAATTCCCGTTTCACAGATGAGGTAGACTGAGGTACACTGTGAGCCCATTGTTGGGTAGGGACCGTATCTATATGTTCCCAACGTGTACTCCCCAAGCGCTTGGTACAGTTCTCTGCACACAGTAAGCGCTCAATAAATACGATTGAATGAATGATGAAAGGTAAATGAAGCACGGAGAAGTTAGTGTCTTGCTCAGGTCATACAGCAGACAAGTGGCGGAGCCAGGATTAGAACCCGTAACCTTTTGATTCCCAGACCCGTGCTCTATCCACTACACCATGCTGAGTTCCAGGCCCAAGGGAGGATATGGGCAAGGGGTTGGCGACAAGATAGATGAGATCAAAGTACATAGAGTAGGCTGGCAATAGAGGAGCTGAGTGTGCATATTGGGTCATAGTAGAAGATTAGAAGGGTGAAATACAATGGAGTGAGCTGATAGAGAAATAGCGTGGCTTAGCAGAAAGAGCACGGGCTTGGGAGTCAGAGGTCATGGGTTCTAATCACGGCTCTGCAACTTGTCAGCTGTGTGACTTTGGGCAAGTTGATGATGATGATGAAGTCACTTAACTTCTCTGGGCTTCGGTTACCTCATCTGTAAAATGGGAATTAAGACTGTGAGCCCGTCGTGGGACAACCTGATTATCTTGTATTTACCCCACTGCTTAGAACAGTGCTTGGCACAGAGTAAGTGCTTAACCATGAAGGCGCTGAAACAAGTCCCTGATGCGGGTCGGCGGGAAACATCTTTTGCTGCCGTTCGTCAGGGTGCTCAGGAACATTATGTTACTTTTCTGGATAGGCTCCAAATTGCCATACAGCGTCAGATTGATTAGGGGGAGGCTCGCGGTTATTATTATGTCAATTGGCAATTGAAAATGCCAATGTTGATTGCAGGAAGGCATTGGATCCCCTACGGAACAAGGCAAAGACGATATCAGACCTAATTAAGGCGTGCCAGAATGTTGGCTCTGAGCACTTTAAAGCTGAAATGTTAGCCTCTGCCCTGGCGCAACAACTTACGGTGGCCCGGGCGGCGGTCAAGTGTTTTTCATTTGGACAGGAAGGGCATATAAGGCGTGACTGTCCTAAGAGGTTTAGAACCCGGCAGGTAAGGCGGGACGTTGTGTCCTTACAGCCATGTTCCAGATGTCACAAAGGATTCCACTGGGGTAGTCAGTGCAAATCCAAATTTGATGTGCAGGGCAACCCAGTGTGGCAGCCGGGAGATGGCTGGAGGGGCGTGAAGCCCGGCGCCCCGAGGACGGCACACAGGGCTACGAGAAATTTCGTGGGTCAGGCGGGAGCTCCGCAGTCGGAGGGTCTCCGCGACATGTCTCTGACCTCCGCCCAGCCACCGCAGTGAGCGCCGGGTTGGACCTGGCCACTGCCAGACCAACCGTGATTTCCAATATTAGTGTACATTTGCTGCCGACGGGCGTATTTGGACCCATGCCTCTGAACACCATGGCCCTTTTGATAGGTCGATCTTCAACTAGTAGAAGTGGGCTGTTTGTCTTACCCGGGGTTATTGACCCTGATTATACGGGGGAAATCAGGATTATGGTCTGGACACCGACACTGCCTTGTACGGTTCCCCCGGGTGAACGCACTGTTCAACTTGTTTTGCTCCCAAGTAGGGGAACGGGTGGTAGTGTTCAGACACGGGCTGGAGGTTTTGGCAGTACCGGTCCACCACGGATATTTTGGACGCACAAGGTCTCCCCAGGACAGCTCTTGCTGACCTGTTTGGTCAATAATCGCCCTTTCACTGGCCTGGTTGATACGGGTACCGATGTTACTATCATTCAGCAGTCCCAGTGGCCTCCCGATTGGCTCCTAGTTTGCTCAGCGGCTGCTGTGGCGGGGTCGGTGGGATGCAAGTGTCATGGCAGAGCGCTCATTCCTTGCTGGTTAAAGGGCCAGAGGGCAAACAGGGCGTATTACAACCTTATGTCCTGGCAGTGCCTTGCACCTTGTGGGGACATGACTTATTAGTGCAATGGAACGTGACACTTATGACAAATTTATAGCGGGGGCCACTGCAGTTCAACCTCGCCTTAAACTGACCTGGAAGACTGACAGGCCGGTTTGGGTAGATCAGTGGCCCTTGAAGGGCGAGAGGTTGGTTAAGGCATGATAACTTGTACAGGAACAACTGGCCTTGGGGCATATAGTTCCTTCTACCAGTCCCTGGAACACCCCCATTTTTGTCATACCCAAAAAATCGGGAAAGTGGCGACTACTGCAAGATTTGAGGGCGATTAATGCAGTAATGACACAAATGGGGCCCTTGCAGCCAGGCACGCCTTCCCCGTCCATGCTTGCTGAATCCTGGCATTTAAGGGTGATTGACTTGAAGGACTGTTTTTTCACGATCCCACTTCATCCGGATGATTGTTCGCATTTCGCCTTCTCGGTGCCCACAATCAATAATCAAGGACCACTGGACAGATACCATTGGGTAGTATTACCGCAAGGCATGATGAATAGCCCTACTATTTGTCAGATGGTGGTAGGCTCGGCGCTGGATCGGGTACGGGCAGTCCACCAGCATGCTATTACTTACCATTATATGGATGATATTCTTATTGCAACACAAGATTCGCGAGCCCTCTCACTCGTGTGTCAGAATGCCAACCAGCAACTACAGCACCAAGGGCTGCCGATAGCAGAGGAAAAGGTTCAGACTGTTGCACCCTGGAAATACTTGGGATGGAGACTGTATGAATCTGAAATCCATCTACAGCCCCTGCAAATCGCTAATACCATAAATACCTTGAACGATTTACAGAAACTCTTAGGTACCATTAACTGGCTTCGGCCGATTTTGGGCATAACGATGGAGGAATTATCTCCGCTTTTTCACCTGCTACGAGGTGATCCTGACTTGACATCTTCGCGTACTCTCACCTCGGCAGCATCTACCGCGTTACAGCGCATTGCCACAAGAATACAGACTAGTTATGGTCACCGACGCTCGAACTCCCTCCCAATTCATTTATTGGTGATTTACCACACCTTCCAGCTGTATGCGGTGCTGGGACAATGGACAGAAGGAGCCGTCTCTCCTACTGTGACACTGCGGACCTTGGAGTGGCTTTTTCTCCCTCATTCCTTCTCCAAAACTGTGACAACCCCCATTGAGATGATGGCCCGTCTTATTTCTCACGGGCGCATCAGATGTCAACAATTAATGGGGGATGATCCCTCTTTTATCCACATATCGGTTACTCGTGCCGACCTTGACACCTGGCTATCACAAAGCCTTGTCCTGCAAATTGCACTTGTGGACTTTCTAGGAACGATCGAATACACCCTGCCTAGACACAAATTGTTGCAGTCGTTGCCCCAGGTGCCCTTGCAGCCGCGGATCCTACTTTCACATGTTCCGTTGCCTAAGGCATGCACAGCCTTTGTAGATGGGTCCGGAAAGACAGGAAAAGCGGTAGTCGTCTGGAAAGGTGTCTCAGACGCTTGGGAATCAGACGTTTTCCAAGTAGAGGATCAACACAGATTGTGGAATTGGTCGCCGTTGTACCCGCCTTTGAATTATTTACATCCGAAGCTCTGAATCTGATGGTAGACTTTGCGTACGTCTCGGGTGTAGTCCAATGCCTGGAAGGGGCCTTTATTAAGGAAGTGGAGAATCGCCTCTTATTCCAGTTGCTTTTGCGATTATCACGCCTGTTAACACAAAGGTCTCACCCATATTTTGTCGTCCATATTGGATCACATACGGCATTGCCTGGTCCGCTGACCGAGGGCAATGCTGTAGCGGATGCTTTAAAAATGCATGTAGTACGGCCAAATCTGTTAACACAGGCCCGGCTTTCACACGACTTTTTCCACCAGAATGCACGGTCTCTGCACAAACAATTTTTGCTGACTATTCAACAAGCACGGGATATTGTTCGCGCTTGTCCTGATTGTCAGCAACTGTCCTCAGTACCTATTCCTGCAGGGGTAAATCCCCGCGGCCTTCGATCCAATGACATTTGGCAATCCGATGTTACTCATGTGGCCGAATTTGGTAGAATGCATTATGTTCATGTTACTGTTGATTCCTTCTCGCATCTTGTTGTGTCCACCGCCCATGCAGGTGAAAAGGCTTGTGATGTGGTTTGACATTGGTTGCACTCTTTCGCCGTCATGGGGGTCCCGGTTACTGTTAAAACTGATAATGGCCCCACCTATGTCTCTCGTAGGGTACAACTCTTTCTCCAAGATTGGGGGGTGCACTATATCACTGGTATCCCTTACTCCCCTACTGGGCAAACCATTGTGGAACGGATGCACCACACGCTTAAAGCTTTGTTGTTTAAACAAAAGAGGGGGAATCCCACGGGAATGACTCCCCAGGAACGGTTATACAACGCTACATACGTTTTAAATTTTTTAACCTTGTCTGATTTGTCCCTTACAGCAGCACAGCACCACTTTGGACGAGACACTACGTGGCCCGAACGGCCCCGTGTGTACTATAGACCTCTTGGGTCGGAACAGTGACAAGGACCTGCACCTCTGATCACCTGGGACCGTAGGTACGCCTGTGTTTCTCTGCCTTCTGCCCCCTACTGGCTGCCGGCGCGTTGTGTCAACCCCTTCAAGACGAGATACCACAGTCTGCGGAGGGCCACCAGGATGGCGAAGACGCGGGCGGCGCCCCCTGATACAGGGCTCAACACGGTCAGAATGTTTATGATCTGGAGCCTGATGGGAAGTCTTTGCCTGCTTGTCACGGGGAATGGACTTTACTGGGCACATATTCTTGATCCACCCATCTTCAAACCTATAACCTGGTGGGATCCTACACCACCTATCGGGAATAATGACACTGAATGGATAGGTGGGCTATGGATGCCCACGACGATAGACGGAGGGTCTAATAATACTAGTTTTTTTGCTATCAATAATACTGAGTTTGTAACTGACCTTCCACCCCTTTGTCTGACAACTAGTAATGACAGCGACATCGGCTGTATCCTGCTGTCGCCTCAGCAACACCTGATAGTGAGGGACAAACGTGGAACAAAGAATTCCACTTTAATAAGCCTCCCGGGTGTGTCTGGAGCATCGTCTCTCCACAACCTGTTGGAAGTGGGGATATATACCCCCGACTTGCCCCGTTGTACGGAGGACCCACTCACAAACAACACATGGATAAACTGGCAGCCATGTCACGGGCTCGGTCCTGATCCTGTTACTGTCAACGGAACTACGGGAACCATTTTGGACTGGGGCCCTCATGGGGTTTTGTGGGACTCGGCAAGCAACCACTCTGTTGGGTTCGGCATCCATAACCATAGTGTATCCTGGCATGGTGGTGGGTTAGCTGGGCCTCTTTTACAGTTTTATATGCCCGCCAACAACACTGCCTCCCCATTACATATGGGTATTTGGCATTTAGGCTTCGCATTTATTAATCGTACCTTATGGAATCTCACCCTGTCTAATCATACTGCTAACGCTACACAGTTACAAACTGATCCAATTATGATTTGCACTTCACATCCCTACATCTTTGCAATGGCACAAGTGGCTACCATAAGCCACTGCAACCATACTTACTGTACTAACCTGACCAATGTATGGCATGGAAATTGTTTCTCTTCTTTAAATGCACTACGCAACAATCTTACCATTATATTTCCCCTGTGTCAGTGCCCGGAGCTGTGGCTTCCTGTTAACCTTACACGTTCATGGGAAGGTGAGAGCGGTCTAGGTCATTTTGCGCGAATAATCCAGGAAGAGGTGGATAGCTCTAATAAGCGCCGAAAACGATTCATAGGGTGGTTGATATTTGCACTTGTATCTGCTATTGTCATTCTGGCTTCTGCGACCACTGCCATCGCCTCACTGGCACAGTCAGTGCAGACGGCAAAGGTGGTAGAGGAAACACTAACCAACGTCACGCAGGAATTTGAGATGCAGGAACATATTGACGAGGAAATCATGGTGAGATTACAGGCCCTCGAGGCAGCGGTAATCTGGCTCGGAGATCGACAGACAGCCTTAAAGACCCGCCTTTCGTTGCATTGTGACTGGGAGCATATCTCTGGAGGCCTTTGTGTGACCCCCTTGCCCTGGAATGCTACCACGCATCCATGGGATACGGTCAAAGAACATCTGCAAGGCGCGTTTGACATGTCCTTACAGCACGATGTCCGATTCTTGCATGATCAATTAAAAGGCCAGATTCAGCAGCTGCAGGCATTAAACACTCAGAATGTTCTCGAAACCCTTCAACGGGATATGTCTTGGTTGAACCGAAAACGTGGTTCACGGGGTTAAATCTGCACGTCTGGGTATTTGTCGGTATAGCAGGAGCCTTGCTTTTCTTCCTTTTGCTCTTTTCCTATATGACATGCTTGCTTATTAGGGCCACACGCACGGTTGAAGCTCGTGTGATGGCCACGTTGCTCATTAATGGTATAGTTATAAATAAAGGAGGGGGAGATGTGGGGAACAGCAGGGCCAAGTAAAGAAGGAACTGAGAGGAGAGCTGTGGCCAAAAGGCTGGTCTCTGTTATTGCAGCCTTGTACAGGCAAGAGTTACTTGTTATCGGCCCAATGGCCTTGCATAGATGGGTGTTGCTTGTTGCTGGCTTTGAAGGTTATGGAATTGCTGTGTTAGTTATGAACACAGGAACATGGAAAAATACAGAATGTGACTGAAAGATATAAAAGTCTTGCTGCTTAACCCAATAAACGATTTCGTGCTTACCCTAGCCGGAGTCCGTGCCTTCATACACCACACTTAACAAATACCAAGATCATTATTATTGAGTGCCTTCAAGCCTATTTTAGGGAATTTTTATTTGATGTAGCAGTAGGTAGGCAATCATTGGAGGTTTTGAGTAGTGAGGAGATGTGGACTGACCATTTTTTCAGAAAAATGATACGGGCAGCAGAATGAAGTATGGACTGGAGAAGAAGAGATAGGAGACAGAGAGATCAGCGAGTTGGCAGATTCAGTAGTCAAGGCAGAGAAACAAAACAGTCTAGTCGATAGGGCATGGGCCCGGGAGTCAGAAGGACCTGAGTTATAATCCCAGCCCTGCCATTGGTCTGCTTTGTGACCTTGGACAAGTCATTTAACTTCTTTTCAGCGTGGCTCAGTGGAAAGAGCACGGGCTTTGGAGTCAGAGGTCAGGAGTTCAAATCCCGGCTCCACCAATTGTCAGCTGTGTGACTTTGGGCAAGTCACTTAACTTCTCTGTGCCTCAGTTACCTCATCTGTAAAATGGGGATTAAGACTGAGCACCCCTTGGGACAACCTGATTACCTTGTAACCTCCCCAGCGCTTAGAACAGTGCGTTGCACATAGTAAGTGCTTAATAAATGCCATCATCATCATCATCCTTCTTTGG

>ERV_Env-Tac7.2

AGTGTGGGGAACAGCCGGGCCAAGTAAACAAGGAACTGAGAGGAGAGCTGTGGCCAAAGGGCTGGTCTCTGTTATCACAGCCTTGTACAGGCAAGAGTTACTTGTTATCGGCCCTACGGCCTTGCATGGACGGGTGTTGCTTGTTACTGGCTTTGAAGGTTACAGAATTGCTGTGTTAGTTATAAACACAGGAACATGGAAAAATACAGAATGCGACTGAAAGATATAAAAGTCTTGCTGCTTAACCCAATAAACGATTTCGTGTTTACCCTAGCCGGAGTCCATGCCTTCATACACCACAAATGGCGCCCAAATCGGGAATGCAAATCTGAATATATGATCCGGTGGTGTACCCGCAGCTGGACGGAGGATATGAGTTCGTAAAAACTCCGGCCGGAGACGTTAGGACATCACCATCCGCAGGACGGAGAGATGAGCAGCCGTGGAAGAATTGACAGCATGGGGCAATCAGTCACGAGAGAGCAGAAGTTGCAGCTTGAGATTTTGCAACGTATTTTAAAGGAGCAAGGGTTTAAGGTGCAACCCATACCACTGGTCCAGTTATTAGTATGGATTCGGGATCATTGTATATGGTTCCCTGAAGACGGTTCTTATAATCTTACCTTATGGCAAAAGGTGGGCTAGGAGCTACAGACTCGTGACAGCTTGTCCCAGCCCCTCCCCAAAGGGGTACAGATTACGTGGAAAACTGTTTTTACGGCGCTGCGGGCGCTGAATCCCGCTGAATAGGTGCTGCAGGAGGCGCCGGCGACGCGCGACTTGGGGGCAGGGAGTGATCAGTGTCCTTTGAAGTTTTGCCTCCAACAAAAGAATTGGAGCCGGTGTACGTTACTGTTCCGGATCCCGAGGACTTGTATCGGGACCAGGATCATAATGACTCTGGCGATCCTTCTGACTCTGGGACTGCAGACCCGAAGGAGGGATCTGATTTATACCCTCCTCTGGCCCCTCTTACAACTACGGCTGCTGCCGCACCCACAGCAAAGGACCTTTTGCAGACCCAGCGCCTGCAGGCGGTATCCCGTTCGTCGATCGTTCACCCTCCGCCCCTTGCTCCCCCTTTGTCTGCTGCGCCGCCGTATGTATATGCTGTGCCTGCTACGGCGCAGCCTGCCTATACCGGGGCCCTTTCGGGCTGCCCGCAGGAGGTATTGCGCCAAGGGGATGTACAGCTACTGCAAGCGATTCCGGTGGTGTATCTTCCCGGGCGGCTGGCTCGGTATGACTCTCTCCCTTATGAGTTGATTGAGGAGTTGCGGAAAAGTGCTCTAGATTATGGGTTACAGTCGTCCTATACAATGAATCTAATAGTGGCTGTCTCTGAATCGTATGTGATGACACCCCATGATTGGCGTACCTTGTTCCGTTTATTGATCAGTCCGGCCCAGTTTTCTGTTTGGGACTCTGAGAATCGGGAAGCTGTTACCTTGCAAGTTATGGATAATCTTGCTAACAATATTAATCTTGGGGTGGATGAGTTAGTAGGGCAAGGGAAGTTTGCCACGCCCCAAGCACAGGTTCAACTGAACCGGGTGGCCTTTACCCAGGCTGCCTCCTTAACCATGAAAGGCGGTAGTCGTCTGGAAAGGTGCCTCGGACGCTTGGGAATCGGATGTTTTCCAAGTACAGGGATCAACCCAGATTGTGGAATTGGCCGCCGCTGTGCGCGCCTTTGAATTATTTGCATCAGAAGCTCTGAATCTGATAGTAGACTCTGCGTACATCTCGGGTATAGTCCAACGCCTGGAAGGCGCCTTTATTAAGGAAGTGGAGAATCGCCTCTTATTCCAATTACTTTTGCGATTATCGCGCCTGTTAACACAAAGGTCTCACCCGTATTTTGTCGTCCATATCAGATCACGTACGGCATTGCCTGGCCCGCTGACCGAGGGCAATGCTGTAGCGGATGTTTTAACAATGCAGGTCGTACGGCCAAATCTGTTAACACAGGCCCGGCTTTCACACGAGTTTTTCCACCAGAATGCGCGGTCTCTGAGCAAACAATTTTCGCTGACTATTCAACAAGTGCGGGATATTGTTCGCGCTTGTCCTGATTGTCAGCAACTGTCCTCAGTACCCATTCCTGCGGGGGTAAATCCCCGGGGCCTTCGATCCAATGACATTTGGCAATCTGATGTCACTCATATGGCCGAATTTGGTAGACTGTGTTATGTTCATGTTACTGTTGATTCCTTCTCGCATCTTGTTGTAGCCACTGCCCATGCAGGAGAAAAGGCTCATGATGTGGTTCGACATTGGTTGCACTCTTTCGTCATCATGGGGGTCCCGGTTACTGTTAAAACTGACAATGGTCCCGCCTATGTCTCTCGTAGGGTACAACTCTTTCTCCAGGATAGGGGGGTGCGCCATGTTACTGGTATCCCTTATTCCCCTACTGGGCAAGCTGTTGTGAAACGGATGCACCACACGCTTAAAGCTCTGTTGTTTAAACAAAAGAGGGGGAATCCCACGGAAATGACTCCCCAGGAACGGTTATACAAGGCTACATACGTTTTAAATTTTTTAACCTTGTCTGATTTGTCCCTTACGGCAGCACAGCACCACTTTGGACAAGACACTACGGGACCCGAATGGCCCCGTGTGTACTATAGACTTCTTGGGTCAGAACAGTGGCAAGGACCTGTACCTCTGATCACCTGGGGCCGTGGGTACGCCTGTGTTTCTCTGCCTTCCGGCCCCTACTGGCTGCCGGCGCATTGTGTCAAGCCCTTCAAGACAAGGATAGCACAATCTGCGGAGGGCCACCAGGATGGCGAAGACGCGGACTGCGTCCCCTGGTACGGGGCTCAACACGGTCAGAATGTTTATGATCTGGAGCCTGATGGGAAGTCTTTGCCTGCCTACCACGGGGAATGGTCTTTACTGGGCACATATTCTTGATCCACCCATCTTCAAACCTATAACCTGGTGGGATTCTACACCACCTATCGGGAATAATGACACTGAATGGATAGGGGGGGTATGGATGCCCCCGACGATAGACGGAGGATCTAACAATACTAGTTTTTTTGCTCTCAACAATACTGACTTTTTAACTGACCTTCCACCCCTTTGTCTGACAACTAGTAATGACAGCGACATTGGCTGTATTCCGGTGTCACCTCAGCAACACCTGATAGTGAGGGACAAACGTGGAACAAAGAATTACACTTTAATAAGCATCCCGGGTGTGTCTGGAGCATCGTCTTTCCACAACCTGACAGAAGTGGGGATATATACCCCCGACTCGCCCCGTTGTACGGAGGACCCACTCGCAAACAATGCACGGTTAAACTGGCAGCTGTGTCGCGGACTCGGTCCTGATCCTGTTAACCTCAACGGAATTACGGGAACTATTTTGGACTGGGACCCTCATGGGGTTTTGTGGGACCCGGCAAGCAACCACTCTGTTGGGTTCGGCATCCATAACCATAGTGTATCCTGGCATGGCGGTGGGTTGGCTGGGCCTCTTTTACAGTTTTATATGCCCGCCAACTACACCGCCTCCCCATTACATACGGGAATTTGGCGTTTACGCTTCGCGTTTATTAATCGTACCTTATGGAATCTCACCCTGTCTAATCATACTGCTAACGCTACACAGTTACGAACTGATCCAATTATGATTTGCACTTCACATCGCTACGTCTTTGCGATGGCACCAGTGGCTACCATAAACCACTGCAAACATGCTTACTGTATTAACCTGACCAATGTATGGTATGGAAATTGTTTCTCTTCTTTAGATGCACTATGCAACAATCTTACCTTTATATTTTCCCTGCCTCGGCGCCCGGATCTGTGGCTTCCTGTTAACCTTACACGTTTATGGGAAGGAGAGAGTGGTCTAGGTCTTTTTGCGCGAATACTCCAGGAGGAGGTGGATAGCTCTAACAAGCGCCAAAAACGATTCATAGGGTGGTTGATATTTGCACTTGTATCTGCTATTGTCATTCTGGCTTCTGCGACCACTGCCGTCACCTCACTGGCGCAGTCAGTGCAGACGGCGAAAGTGGCAGGGGAAACACTAACCAACGTCATGCAGGAATTTGAGATGCAGGAACGTATTGATGAGGAAATCATGGTGAGATTACAGGCTCTCGAGGCAGCGGTAATCTGGCTCGGGGATCGACAGACAGCCTTAAAGACCCGCCTTTCGTTGCATTGTGACTGGGAGCATATCTCTGGAGGCCTTTGTGTTACCCCCTTGCCCCGCATCCATAGGATACGGTCAAACAACATCTGCAAGGCGCGTTTGACACGTCCTTACAGCACGATGTCCGATCCTTGCATGATCAATTAAAAGGCCAGATTCAGCGGCTGCAGGCATTAACCACTCAGAATGTTCTCGAAACCCTTTAATGGGATATGTCGTGGTTGAATCCGAAAACGTGGTTCACGGGGTTAAATCTGCGCGTCTGGGTATTTGTCGGTATAGCAGGAGCCTTGCTTTTCTTCCTTTTGCTCGTTTCCTATATGACATGCTCGCTTATTAGGGCCAAACGCACGGTTGAAGCTCGTGTGATGGCCACGTTGCTCATTAATGGTATAGTTATAAATAAAGGATGGGGAGATGTGGAGAACAGCAGAGCCAAGTAAACAAGGAACTGAGAGGAGAGCTGTGACCATAAGGCTGGTCTCTGTTATCACAGCCTTGTACAGGCAAGAGTTACTTGTTATCGGCCCTACGGCCTTGCATGGACGGGTGTTGCTTGTTACTGGCTTTGAAGGTTACAGAATTGCTGTGTTAGTTATAAACACAGGAACATGGAAAAATACAGAATGCGACTGAAAGATATAAAAGTCTTGCTGCTTAACCCAATAAACGATTTCGTCCTTACCCTAGCCGGAGTCCGTGCCTTCATACACCACA
